# A method for genome-wide genealogy estimation for thousands of samples

**DOI:** 10.1101/550558

**Authors:** Leo Speidel, Marie Forest, Sinan Shi, Simon R. Myers

**Affiliations:** Department of Statistics, University of Oxford, Oxford, UK; Université du Québec à Montréal, Montréal, Canada; Wellcome Centre for Human Genetics, University of Oxford, Oxford, UK

## Abstract

Knowledge of genome-wide genealogies for thousands of individuals would simplify most evolutionary analyses for humans and other species, but has remained computationally infeasible. We developed a method, Relate, scaling to > 10,000 sequences while simultaneously estimating branch lengths, mutational ages, and variable historical population sizes, as well as allowing for data errors. Application to 1000 Genomes Project haplotypes produces joint genealogical histories for 26 human populations. Highly diverged lineages are present in all groups, but most frequent in Africa. Outside Africa, these mainly reflect ancient introgression from groups related to Neanderthals and Denisovans, while African signals instead reflect unknown events, unique to that continent. Our approach allows more powerful inferences of natural selection than previously possible. We identify multiple novel regions under strong positive selection, and multi-allelic traits including hair colour, BMI, and blood pressure, showing strong evidence of directional selection, varying among human groups.

---

Large-scale genetic variation datasets are now available for a variety of species, including tens of thousands of humans. In principle, all information about a sample’s genetic history is captured by their underlying genealogical history, which records the historical coalescence, recombination, and mutation events that produced the observed variation patterns. In practice, several key existing approaches (e.g., Refs. [1,2]) leverage an underlying coalescent model, because this provides a flexible modelling framework and is the limiting behaviour of a variety of finite-population models^3,4^. However, inference under the coalescent is complicated by the structure of the model, uncertainty over the correct genealogy conditional on observed data, and the large resulting space of possible sample histories^5^. Other approaches^6–11^ use more heuristic approximations to the coalescent, sometimes reducing accuracy: regardless, all published existing methods scale to tens or a few hundred samples at most.

As a result of these issues, the use of direct genealogy-based inference to detect recombination events, date mutations, and reveal evidence of positive selection has been limited to smaller datasets^1,2^, while for larger datasets approaches based on data summaries^12–14^ or downsampling^15,16^ have predominated. A diverse set of tools have detected genetic structure that is in good agreement with geopolitical separation over generations^17^. Admixtures of ancient populations have been identified and dated^18^. Other applications have found bottlenecks in population sizes that are consistent with anthropological evidence of initial human migration from the African continent^15,19–21^ and evidence of subsequent introgression with archaic humans, such as Neanderthals^22^.

We have developed a scalable method, Relate, to estimate genome-wide genealogies (see **Figure 1; Methods;** URLs for implementation). Relate separates two steps; firstly identifying a genealogical framework at each site in the genome, which describe ancestry relationships among sequences but not the times of particular events. Secondly, these times are estimated after mutations are mapped to branches of these trees, allowing for variable population sizes which are inferred from the data, to produce complete genealogies. These are then used directly for downstream inferences. Our approach approximates the coalescent model, but performs as well as or better than existing approaches in our simulations, whilst being thousands of times faster.

**Figure 1.**
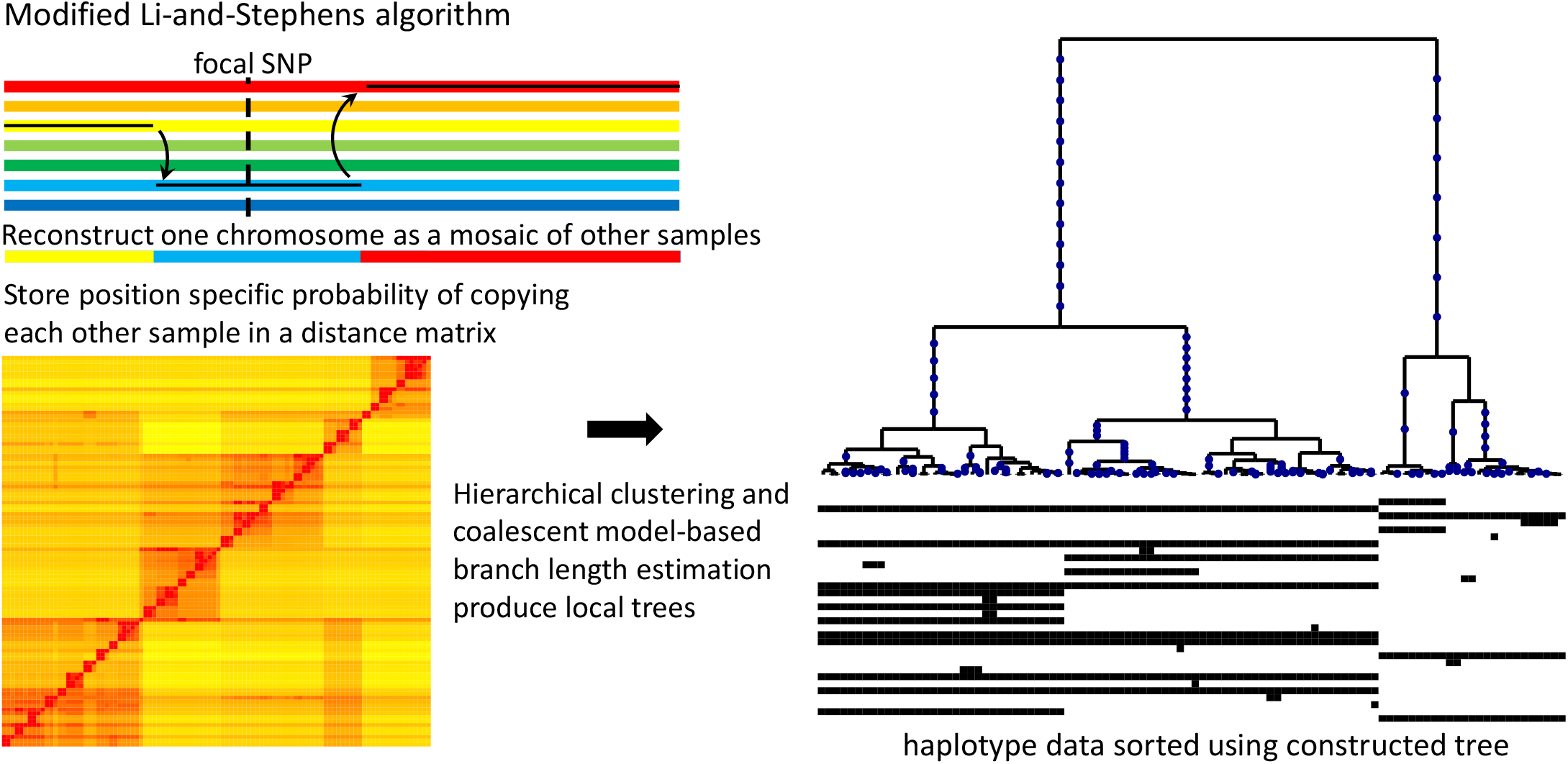
Relate Method overview: Our method applies a version of the Li-and-Stephens algorithm^15^, modified to take ancestral and derived states into account, to calculate at a focal SNP (dotted vertical line) a position-specific distance matrix (bottom left) containing log-likelihoods of copying from each other sample, Our tree builder uses the resulting inferred distance matrix to coalesce haplotypes (right-hand side). After mapping mutations to their corresponding branches, we estimate branch lengths using an MCMC algorithm that employs a coalescent prior model.

We demonstrate the utility of a genealogy-based analysis by applying Relate to 4956 haplotypes of the 1000 Genomes Project dataset^23,24^. We estimate population sizes of all 26 populations in the dataset and their split times using cross-coalescence rates between populations. In agreement with a previous study, we identify an increase in the mutation rate of TCC to TTC mutations, which we date at around 10,000 to 20,000 years ago^25^. The estimated genealogies contain a strong signal of introgression between Neanderthals and modern humans in Eurasia, and a weaker introgression signal between modern East and South Asians and Denisovans, alongside other signals specific to African groups. Finally, we suggest a test statistic that can identify loci under positive selection by tracking frequencies of mutations through time. We demonstrate that, for biologically plausible scenarios of selection on complex traits, where selection is relatively weak, this test is more powerful than the integrated Haplotype Score (iHS)^26^, and we identify genomic regions under strong positive selection that were previously unreported. We find a remarkable enrichment of SNPs identified in genome-wide association studies (GWAS) among these targets of selection, and identify evidence of widespread directional polygenic adaptation, using SNP-trait associations identified in GWAS.

## Results

### Overview of the Relate approach

At each particular position along the genome, Relate first identifies a non-symmetric distance matrix whose rows each estimate the relative order of coalescence events between a particular sequence and the remaining observed sequences, at that position. To do this, Relate uses the posterior probabilities output by a hidden Markov model (HMM) similar to that proposed by Li and Stephens^27^, but leveraging knowledge of ancestral and derived status at each single nucleotide polymorphism (SNP) to improve speed and accuracy. The distance matrix is then used to construct a rooted binary tree using a bespoke algorithm. Mathematical arguments demonstrate, encouragingly, that if the “infinite-sites” model is satisfied so that each observed mutation occurs exactly once, our approach is guaranteed to generate a set of genealogies exactly producing the observed data, in the limiting cases where either there is no recombination, where the recombination rate is very large, or where all recombination occurs in intense widely spaced hotspots (Supplementary Note: Method details). Because the distance matrix is position specific, these binary trees adapt to changes in local genetic ancestry that arise due to recombination. In practice, we save computational time by only rebuilding trees at a subset of sites along the genome (**Methods**).

The binary tree construction step does not estimate branch lengths. To achieve this, while allowing for variable population sizes over time, we first map mutations onto each genealogical tree and then apply an iterative Metropolis-Hastings type Markov Chain Monte Carlo (MCMC) algorithm to estimate branch lengths under a coalescent prior. We simultaneously estimate a stepwise varying effective population size through time, using the genome-wide collection of estimated genealogies (**Methods**). Our final time estimates then account for changes in population size, assuming an unstructured population. We can also explore population stratification within a large sample, by leveraging Relate’s estimates of the coalescence rate through time of any pair of sampled sequences. By averaging pairwise coalescence rates within and across groups, we obtain estimates of effective population sizes for sub-populations and cross-coalescence rates between populations. As we show in the next section, this can provide accurate estimates despite the fact that our tree-builder does not account for such population stratification.

### Simulations

We evaluated Relate in terms of its speed, accuracy of inferred trees, robustness, and ability to infer evolutionary parameters, by simulating data under the standard coalescent with recombination using msprime^28^. We compared performance to ARGweaver^2^, which samples from a time-discretised approximation of the standard coalescent with recombination and a constant population size, and which we therefore expect to perform well on the simulated datasets. Relate was >4 orders of magnitude faster than ARGweaver, for cases we were able to apply the latter, and also much faster than RENT+^11^ (**Figure 2 a,b**). Our approach scales linearly in sequence length and quadratically in sample size *N*, allowing it to be applied to e.g. 10,000 human samples genome-wide, using a compute cluster.

**Figure 2.**
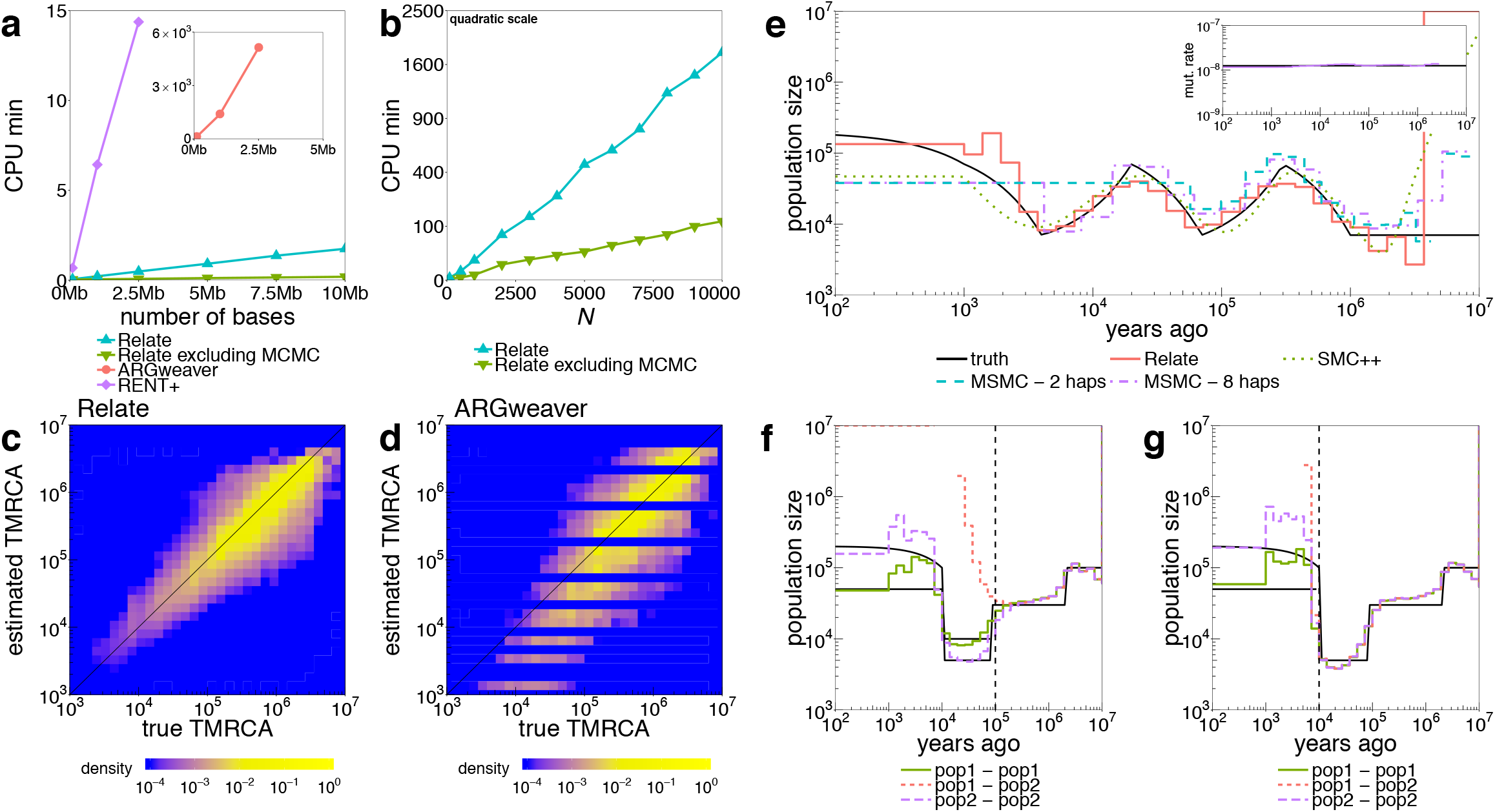
Simulated Data: **a**, Runtimes of Relate, RENT+ and ARGweaver in CPU minutes as a function of the number of bases simulated with *N* = 200, *μ* = 1.25 × 10^−8^, 2*N_e_* = 30,000, and recombination rates taken from human chromosome 1. We also show the runtime of Relate excluding the estimation of branch lengths. **b**, Runtime of Relate in minutes as a function of sample size *N*, where we simulate 2.5Mb for each data point. Other parameters are the same as in **a;** y-axis is on a quadratic scale. **c**, True TMRCAs for pairs of haplotypes (x-axis) versus those estimated by Relate (y-axis). **d**, As **c**, except showing results for ARGweaver. **e**, Comparison of population size estimates across methods, for simulation with an oscillating population size^15^ Inset shows the mutation rate over time estimated by Relate. **f,g**, Population-specific estimates of population size and cross-population coalescence rates for a simulation with a discrete bottleneck for two populations that separated 80,000 YBP (**f;** vertical dashed line), or 10,000YBP (**g**).

To evaluate accuracy of tree topology and branch lengths, at each locus and for each of the 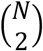 pairs of haplotypes, we compare the estimated time to their most-recent common ancestor (TMRCA) to the truth (**Figure 2c** and **d**), observing improved performance relative to both ARGweaver and RENT+. Compared to these approaches, Relate also showed excellent robustness to errors in the data, as well as to misclassified ancestral alleles, and was able to estimate times well in the presence of changes in population size (Supplementary Figure 3). Other accuracy measures yielded similar results (Supplementary Note: Simulations). We next compare the accuracy of Relate’s inferred population sizes to those based on applying two leading specialist approaches, MSMC^20^ and SMC++^21^. Relate obtains more accurate estimates than these methods, particularly in the recent past, for a variety of previously tested^20,21^ population size histories including oscillating population sizes and bottleneck scenarios similar to those observed in out-of-Africa events of modern humans (**Figure 2 e** and Supplementary Figure 3). While our method assumes a single population when estimating branch lengths, when applied to a combined sample from two diverged populations, it still performs well in recovering their distinct population histories and estimating their split time(s) (**Figure 2 f,g**).

### Genome-wide human genealogies

We applied Relate to data of the 1000 Genomes Project, comprising 2478 individuals with diverse genetic ancestry and approximately 81 million SNPs (see **Methods** for details on data pre-processing). Our method terminated after 4 days using a compute cluster with up to 300 processors (Supplementary Table 1). 86% of all SNPs (>96% of SNPs at >0.2% derived-allele frequency (DAF)) map uniquely to trees, falling to 76% of CpG dinucleotides, which are known to possess strongly elevated mutation rates (Supplementary Figure 5). The number of trees constructed in a genomic subregion is correlated to recombination distance (*r*^2^ = 0.63) and the average tree has 3883 SNPs mapped to it, reflecting block-like structures of human haplotypes between recombination hotspots (Supplementary Figure 5).

We estimated within-group and pairwise coalescence rates for pairs of groups, by first extracting the genealogy for members of a particular subsample of interest embedded within the full genealogy, and then re-estimating coalescence rates through time for this genealogy. We observe a clear out-of-Africa bottleneck following the migration of Asian and European populations (CHB: Chinese in Beijing and GBR: British in England and Scotland shown), and a split from African populations (YRI: Yoruba in Ibadan, Nigeria shown) already visible at 200,000 years before present (YBP) and lasting to around 60,000YBP (**Figure 3 a,b**). This is consistent with recent studies^15,29^ and supports a slow separation between African and non-African groups that might reflect e.g. several out-of-Africa dispersal events. Asian (CHB shown) and European (GBR shown) populations separate more recently, with a clear and much more sudden separation visible at around 30,000 YBP (**Figure 3c**). We are also able to detect, and date, very recent separations <10,000 YBP, such as between CHB-JPT (JPT: Japanese in Tokyo) or FIN-GBR (FIN: Finnish in Finland) (**Figure 3 d,e**). We find a second bottleneck in Finnish samples, occurring around 3000 to 9,000 YBP and after separation from GBR^30,31^, alongside other very recent population-specific events including in Peruvians and Gujarati individuals (Supplementary Figure 6). The Finnish bottleneck is thought to have caused enrichment of certain disease-causing gene variants, commonly classified as Finnish heritage diseases^30,31^. The absence of a strong bottleneck in African populations (LWK: Luhya in Webuye, Kenya and YRI shown) following the departure of European and Asian populations can be seen in **Figure 3f**. All populations show a remarkable increase in inferred population size, often to >1,000,000, in the recent past (Supplementary Figure 6).

**Figure 3.**
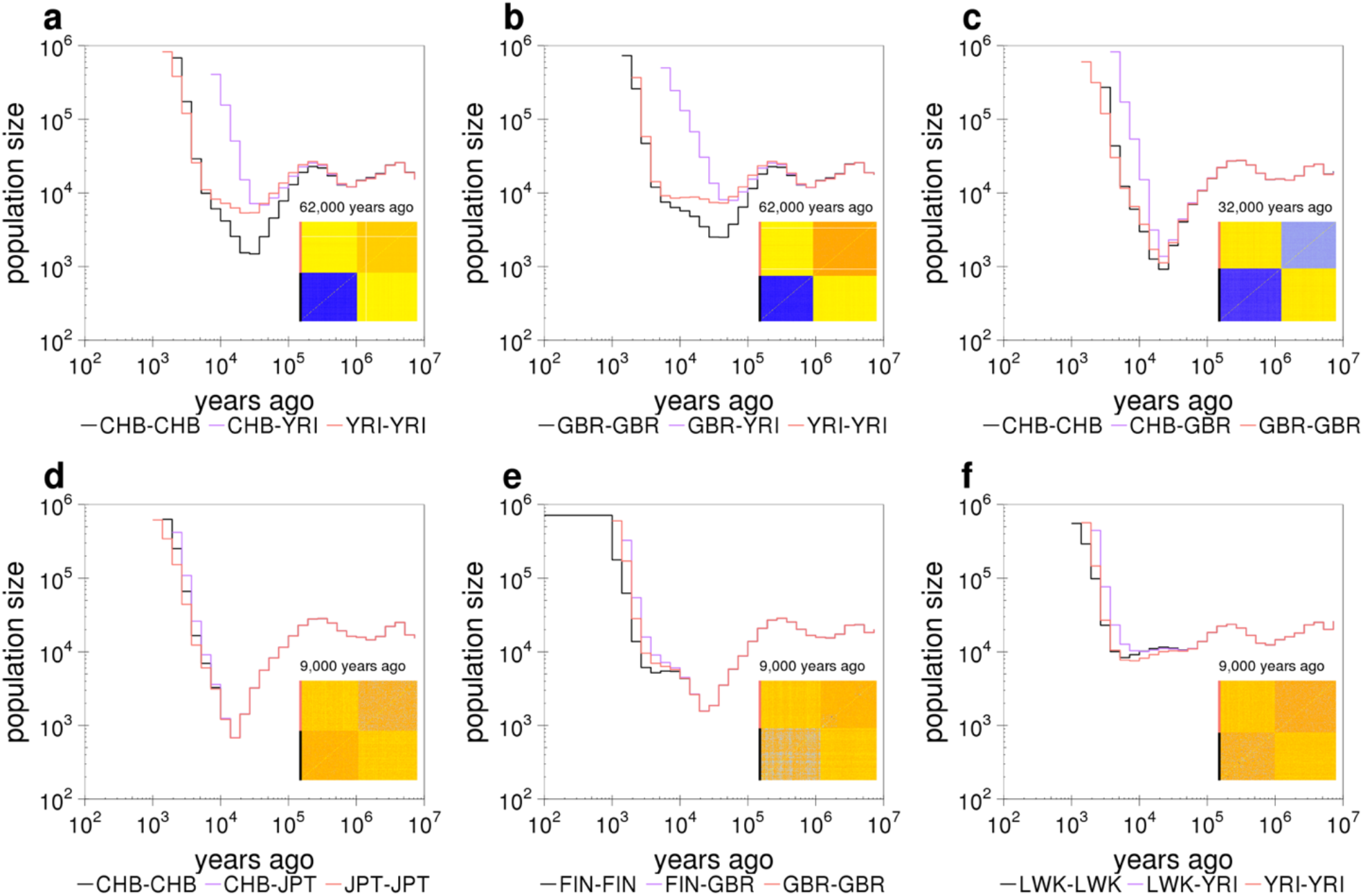
Population sizes and split times in 1000 GP: Relate-based population-specific estimates of population size and cross-population coalescence rates using genome-wide genealogies for CHB and YRI **(a)**, GBR and YRI **(b)**, CHB and GBR **(c)**, CHB and JPT **(d)**, FIN and GBR **(e)**, and LWK and YRI **(f)**. Insets show the matrices of coalescence rates between pairs of haplotypes at the indicated time. Rows and columns are sorted by population labels of haplotypes, as indicated by the colour on the left of each matrix.

It is straightforward to explore the relative mutation rate of particular classes of mutations through time (**Figure 4a**), and as reported previously^25^, this confirms a strong elevation in the rate of mutation types including TCC->TTC trinucleotide changes in West Eurasian groups, which we date to 5,000-30,000 YBP, but infer to be weak or absent in the present day. This approach might usefully be applied in a range of other species in future. Overall, these results support accuracy of our inferred historical relationships, including the timing of a range of different historical events, identified within a single analysis framework.

**Figure 4.**
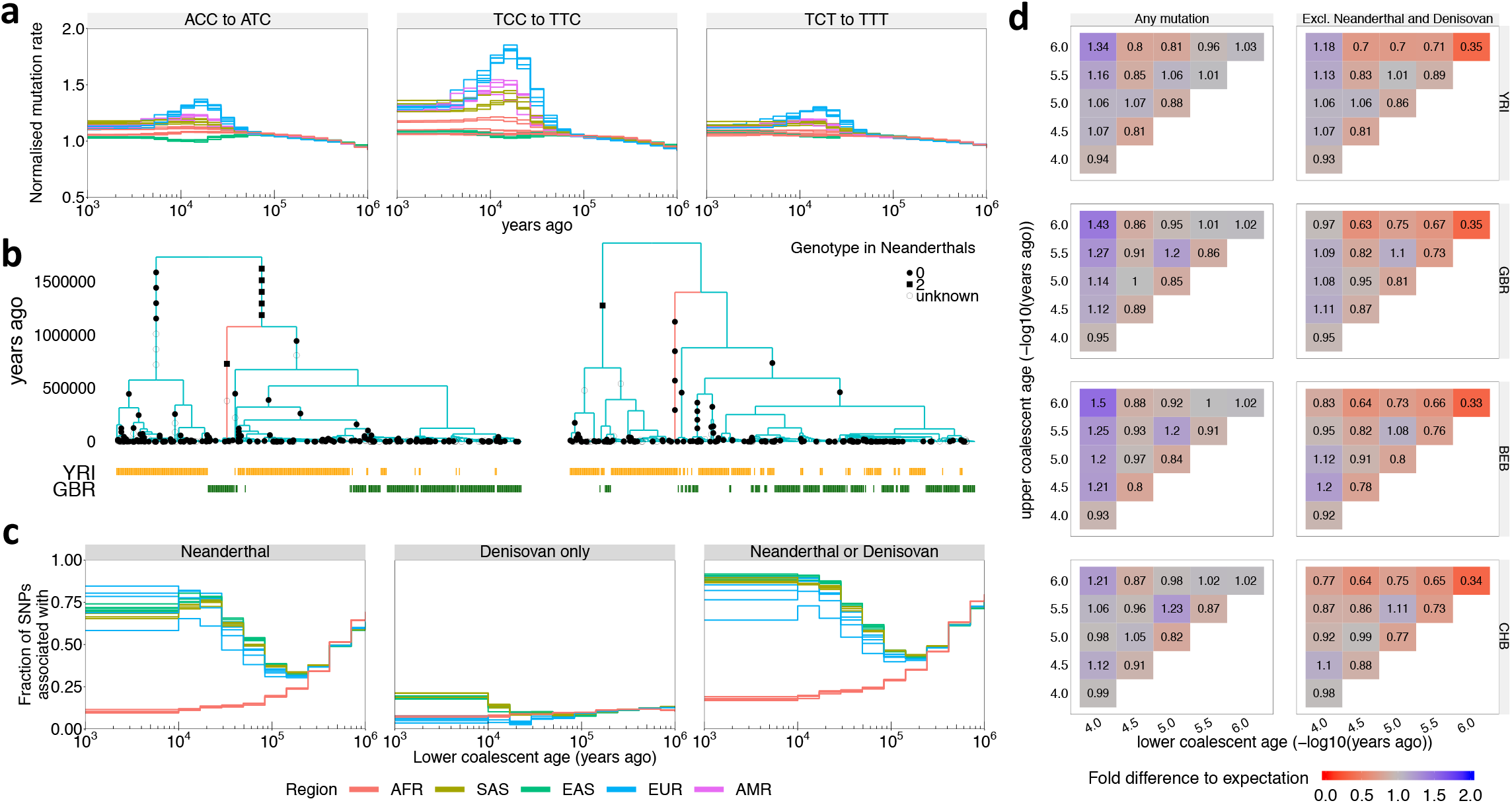
Evolution of human mutation rates and evidence for introgression: **a**, Evolution of mutation rates for three triplet mutations ACC to ATC, TCC to TTC, and TCT to TTT. We elimin ate temp o ral trends share d across different mutation categories by dividing by the mean mutation rate across mutation categories in each epoch. For each population, we then normalise the mutation rates such that the average rate over time equals 1. **b**, Marginal trees for a subregion on chromosome 14 (left) and chromosome 11 (right). The tree on the left contains a long branch with descendants only in GBR (red) consistent with Neanderthal introgression into GBR. The tree on the right contains a long branch with descendants only in YRI (red) consistent with introgression in YRI involving a hominid not closely related to Neanderthals. **c**, Fraction of SNPs on branches with an upper coalescent age >1M YBP that are shared with Neanderthals (left), Denisovans and not Neanderthals (center), or Neanderthals or Denisovans (right) (**Methods**). In **a** and **c**, colours encode geographic regions (AFR: Africa, EAS: East Asia, EUR: Europe, SAS: South Asia, AMR: Americas). **d**, Number of mutations binned by age of upper and lower coalescent event, relative to the expected number of mutations when randomising topology while fixing ages of coalescence events (**Methods**). Right column shows mutations not present in Neanderthal or Denisovan samples.

### Embedded signals of Neanderthal and Denisovan introgression, and unexplained events, within the trees

Tree-based approaches should, in principle, provide information regarding ancient admixture and introgression events. Introgression from distantly related groups in the past is expected to introduce lineages which forward in time can then expand in the tree, and backward in time remain distinct from other lineages, resulting in long branches with unusual numbers of descendants associated with particular times. We therefore identified such branches (>1 million years (MY) in age and with varying lower end), in different human groups (**Figure 4b,c**). It is established that all non-African human groups possess similar levels of Neanderthal introgression, and specific Asian and Austrolasian groups possess admixture from a group related to Denisovans^22,32^. We therefore annotate long branches possessing at least two mutations as being shared with these groups if they possess at least one mutation shared with them, leveraging genome sequences of the Vindija^22^ and Altai^33^ Neanderthals, and a Denisovan^32^ (**Figure 4b** shows one example of likely introgression from Neanderthals into European GBR, but not African YRI individuals). After classifying old branches based on their lower-end times – which provide a lower bound on the introgression time - for branches originating within the last 10,000YBP, 85-90% are shared with Neanderthal or Denisovan for most European and Asian groups (**Methods**). An exception is IBS, which has more long branches shared with African populations (Supplementary Figure 6**b**). In East and South Asian groups, Neanderthal sharing remains high for branches ending up to ~30,000YBP, then slowly declines, while the data instead suggest a very recent arrival of Denisovan DNA (mainly <15,000YBP). This substantial sharing of long branches with Denisovans is restricted to East Asian and South Asian groups. This suggests that aside from groups closely related to Neanderthals and Denisovans, no strongly diverged hominid group has left a major, recent impact in non-African populations studied here. The coalescent-based arrival date of Neanderthal DNA agrees with previous estimates based on LD^34^, and direct evidence of hybrids^34,35^ around 40,000YBP. Moreover, elevation in the sharing of quite deep haplotypes with Neanderthals steadily increases from ~100,000 YBP, which is suggestive of introgression beginning from this time in non-African individuals, although it is important to note that our date estimates for individual events might be over- or under-estimates in some cases.

In contrast to non-African groups, sharing with Neanderthal/Denisovans is very low (<20%) in African populations, and declines towards the present, suggesting minimal recent interactions^22,32^. This is despite the fact that the largest number of long branches observed come from African populations (on average, there are 42,434 mutations on branches with a lower coalescent age < 30,000 YBP and upper coalescent age > 1M YBP in African populations, compared to 7012 such mutations in non-African populations). In fact, 98% of mutations on long branches are unique to populations in Africa, indicative of separate events occurring in non-African and African populations (Supplementary Figure 6**b**). We observe a strong enrichment of mutations with an upper end >1M YBP and lower end <40,000 YBP in YRI, GBR, BEB, and CHB, where this enrichment can be almost entirely explained by Neanderthals/Denisovans in the non-African populations, but not in YRI (**Figure 4d**). In panmictic simulations with matched population size histories, we observe no such enrichment (Supplementary Figure 6**c**). This may be consistent with ancient but uncharacterised population structure within Africa, for which there is increasing evidence^36,37^. **Figure 4b** shows one example consistent with an introgression event in YRI with a hominid not closely related to Neanderthals.

### Powerful tree-based approaches to study natural selection

By directly modelling how mutations arise and spread, genealogical trees offer the potential to powerfully investigate different modes of natural selection, using novel approaches (e.g., Ref. [38]). For example, a recent method, SDS, tests for the presence of positive selection acting on a focal SNP by testing for differences in tip branch lengths between carriers and non-carriers using the density of singletons around this SNP^39^. A tree-based analogue of SDS (trSDS) directly compares tip branch lengths^40^. Here, we propose a class of approaches (Relate Selection Tests) based on estimating the speed of spread of a particular lineage (carrying a particular mutation), relative to other “competing” lineages, over some chosen time range. Rapidly-spreading lineages may carry positively selected mutations. To test for selection over the entire lifetime of a mutation, we condition on the number of lineages present when it first arises, and use as a test statistic the number of present-day carriers. Assuming no population stratification within a group, the null distribution of this statistic can be calculated analytically and is robust in principle to changes in population size through time (**Methods**).

Simulated data (**Figure 5a**) show a close match in null no-selection scenarios of our p-values *p_R_* to the expected uniform distribution, in scenarios (1000 haplotypes) incorporating human-like recombination hotspots and variation in population size through time. In simulations across a range of selective advantages and SNP frequencies (**Figure 5b, Supplementary Figure 4**), our approach increases power relative to SDS and trSDS in all settings, as well as relative to iHS for weaker selection in particular. trSDS is more powerful than SDS, while applying the Relate Selection Test to the *true* genealogical trees yields a test that is uniformly more powerful than other approaches (**Figure 5b**), indicating the strength of using tree-based approaches. In practice, there is therefore some decrease in power from the need to infer trees via Relate. The increased power of our statistic for detecting weak selection might be particularly beneficial when investigating selection on complex, polygenic traits, where small effect sizes mean the selection coefficients on single loci are expected to be small^41^.

**Figure 5.**
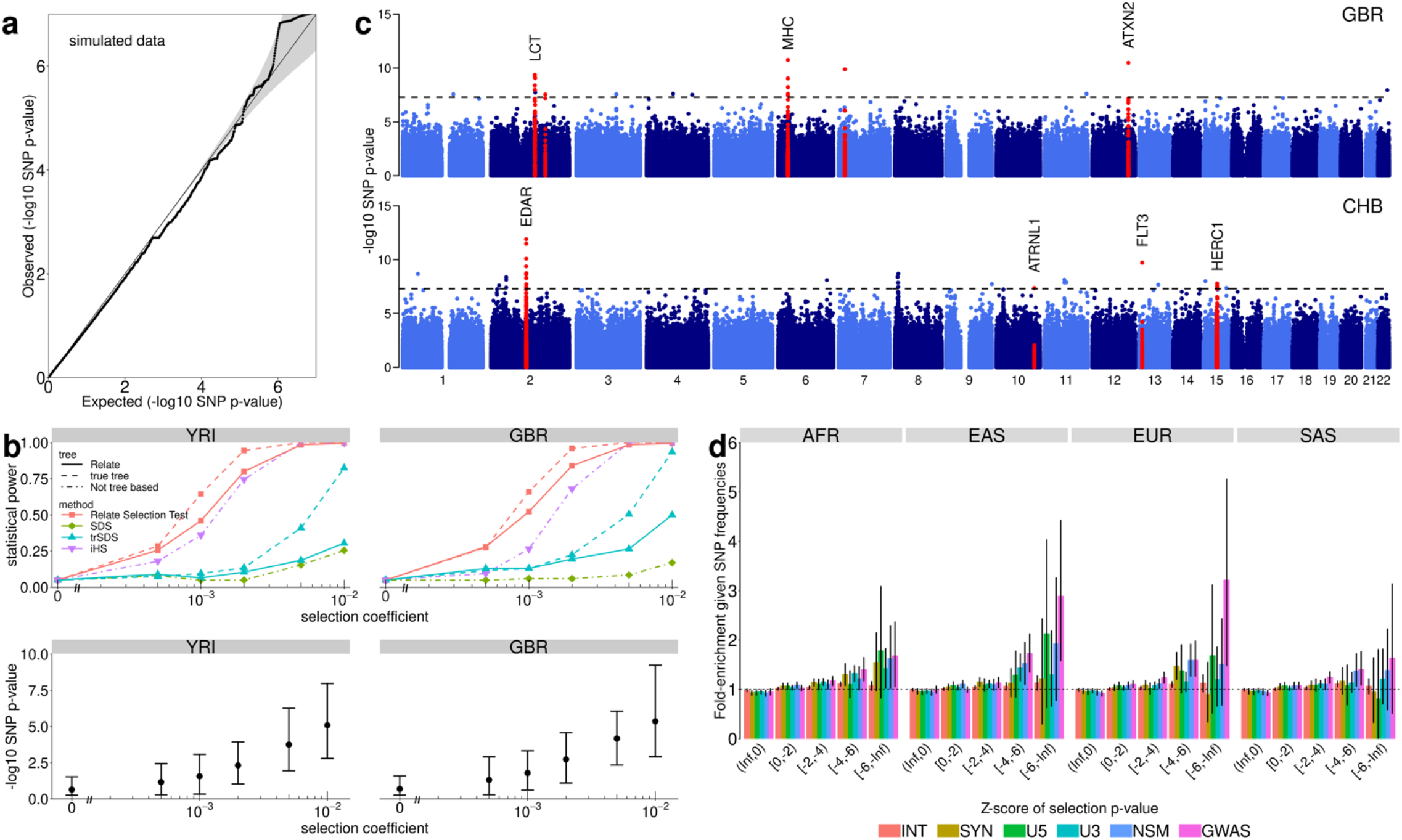
Natural selection: **a**, QQ-plot of p-values for selection evidence of SNPs. We simulated 250Mb for *N* = 1000 haplotypes using the recombination map of chromosome 1 and a bottleneck population size resembling that of non-African populations. **b**, Power simulations using *N* = 1000 haplotypes. We use historical population sizes estimated by Relate for YRI (left) and GBR (right). **c**, Manhattan plot showing p-values for selection evidence of SNPs, for GBR and CHB. We highlight regions containing a SNP with *p*_R_ < 5 × 10^−8^ in at least three populations (see Supplementary Table 3 for a full list), as well as the MHC region in GBR. **d**, Enrichment of functional annotation among targets of selection, conditional on allele frequency. Error bars show 95% confidence intervals estimated from 1000 iterations of a block bootstrap resampling (**Methods**). We group SNPs by mean regional Z-score corresponding to the log p-value for selection evidence, where a smaller Z-score indicates stronger selection evidence. SNPs are binned by partially overlapping functional annotations: intronic mutations (INT), synonymous mutations (SYN), mutations at the 5’ end and 3’ end of a gene (U5, U3), non-synonymous mutations (NSM), and GWAS hits (GWAS).

Calculating *p_R_* for each bi-allelic SNP across 20 populations within the 1000 Genomes dataset (**Methods**) identified 35 regions containing genome-wide significant signals (*p_R_* < 5 × 10^−4^), using the stringent criterion that this threshold is reached separately in each of three or more groups (Supplementary Table 3). Of these, 11 have been previously reported, including the LCT region associated with Lactose tolerance in Europeans, and a mutation in the EDAR gene in East Asian populations^42,43^. In both cases, the causal variant is in high linkage disequilibrium (LD) to the mutation with lowest *p_R_* (*r*^2^ ≥ 0.8). We also observe a previously-detected strong signal of positive selection in the MHC region in GBR^44^ (**Figure 5c**). Among unreported regions, we identify the EDARADD gene – which interacts with the EDAR gene^45^ in the formation of hair follicles, sweat glands, and teeth^43^ – as exhibiting selection evidence in all South Asian populations, as well as the Finnish population and reaching *p_R_* < 10^−6^ in all European populations. In 16 of 35 regions, we identify GWAS catalogue hits (OR=6.44; p=0.01), non-synonymous mutations (OR=2.49; p=0.16), or eQTLs (OR=1.74; p=0.1), in LD with the mutation with strongest selection evidence (*r*^2^ ≥ 0.8, **Methods**), suggesting functional effects, reaching statistical significance for the case of GWAS hits despite the small number of cases tested. Only 8 of the 35 regions are attributed to European populations, and 18 regions are found only for African populations.

Overall, SNPs in functional parts of the genome are significantly enriched among targets of positive selection (**Figure 5d**, see Supplementary Note: 1000 Genomes Project for details). In particular, we find strongest enrichment for GWAS hits, across all considered populations, encouragingly supporting a link between evidence of selection and SNPs with detectable influences on phenotypes at the organism level. Multiple previous studies^46–49^ have attempted to test individual polygenic traits for evidence of directional polygenic selection, but confounding due to population stratification^50,51^ is potentially problematic in practice. To leverage potential power gains of the Relate Selection Test for this purpose, we tested whether derived mutations that increase (or decrease) a trait show increased evidence of directional selection relative to randomly sampled control mutations of the same frequency (Wilcoxon test; **Methods**). For each tested trait we thin GWAS hits to account for LD (Methods) and examined only SNPs showing “genome-wide significant” associations (*p* < 5 × 10^−8^), because confounding due to population stratification is thought to operate through relatively small - but systematic - biases in effect size estimates^50,51^, but is not known to produce false-positives that are genome-wide significant. At each such SNP retained, we use only the association direction, rather than its strength, to offer additional robustness to potential confounding.

If positive selection occurs so as to advantage SNPs influencing a trait in a certain direction, e.g. trait-increasing, we would expect positive selection on trait-increasing mutations, and negative selection on trait-decreasing mutations. In general, we expect our test to be sensitive mainly to the former, because selection will increase frequencies of such SNPs, and the Relate Selection Test has reduced power to identify selection at rarer markers (**Figure 5b**), particularly if selection has varied through time so as to impact mainly standing variation. However, as described further in the Discussion, for traits with a large number of hits, and strong selection, it is theoretically possible for our approach to observe some selection evidence in both directions^52,53^, because to avoid ascertainment effects we condition on SNP allele frequencies at trait-influencing sites. Therefore, we additionally test for differences in present-day DAFs between trait-increasing and trait-decreasing mutations, which can provide orthogonal evidence of polygenic adaptation, aiding interpretation of results we observe (**Methods**). We also note that caution should be exerted when interpreting polygenic adaptation^54^. For instance, a selection signal associated to a particular trait may not necessarily imply a phenotypic change in the same effect direction, because, for example, selection could have occurred to counter-act a large change in phenotype caused by another mutation. Additional issues are considered in the Discussion.

As a positive control, we applied our test to GWAS for hair colour conducted for the UK Biobank^55^ (**Figure 6a**). As in previous studies^49,56,57^, we find a signal for SNPs associated with blonder hair colour among European populations, which is absent in South Asian populations^49,56^. Moreover, we observe strong selection signals for a decrease in black hair colour, as well as an increase in light brown hair colour among all European populations, and weaker signals in South Asian populations, while East Asian and African populations show no evidence of selection. Testing based on iHS scores identifies only some of these signals, and with significance decreasing around 4 orders of magnitude (**Figure 6a**). Next, we applied the same to 84 traits: 6 from the UK Biobank, and 78 with at least 10 genome-wide significant GWAS catalogue association signals in each effect direction. We tested all populations except recently admixed groups; 61 of these 84 showed nominal evidence for selection (p<0.05) in at least one population (**Figure 6b**), with strong geographic clustering, and the most significant signal (*p* = 6 × 10^−14^) for SNPs associated with decreased BMI in CEU. The largest number of selection signals are observed for Europeans, possibly because many GWAS were conducted in these groups. Interestingly, East Asians have the fewest selection signals and no enrichment of low p-values (Supplementary Figure 7) which may partly be explained by their stronger population bottleneck, which would theoretically be expected to weaken selection signals.

**Figure 6.**
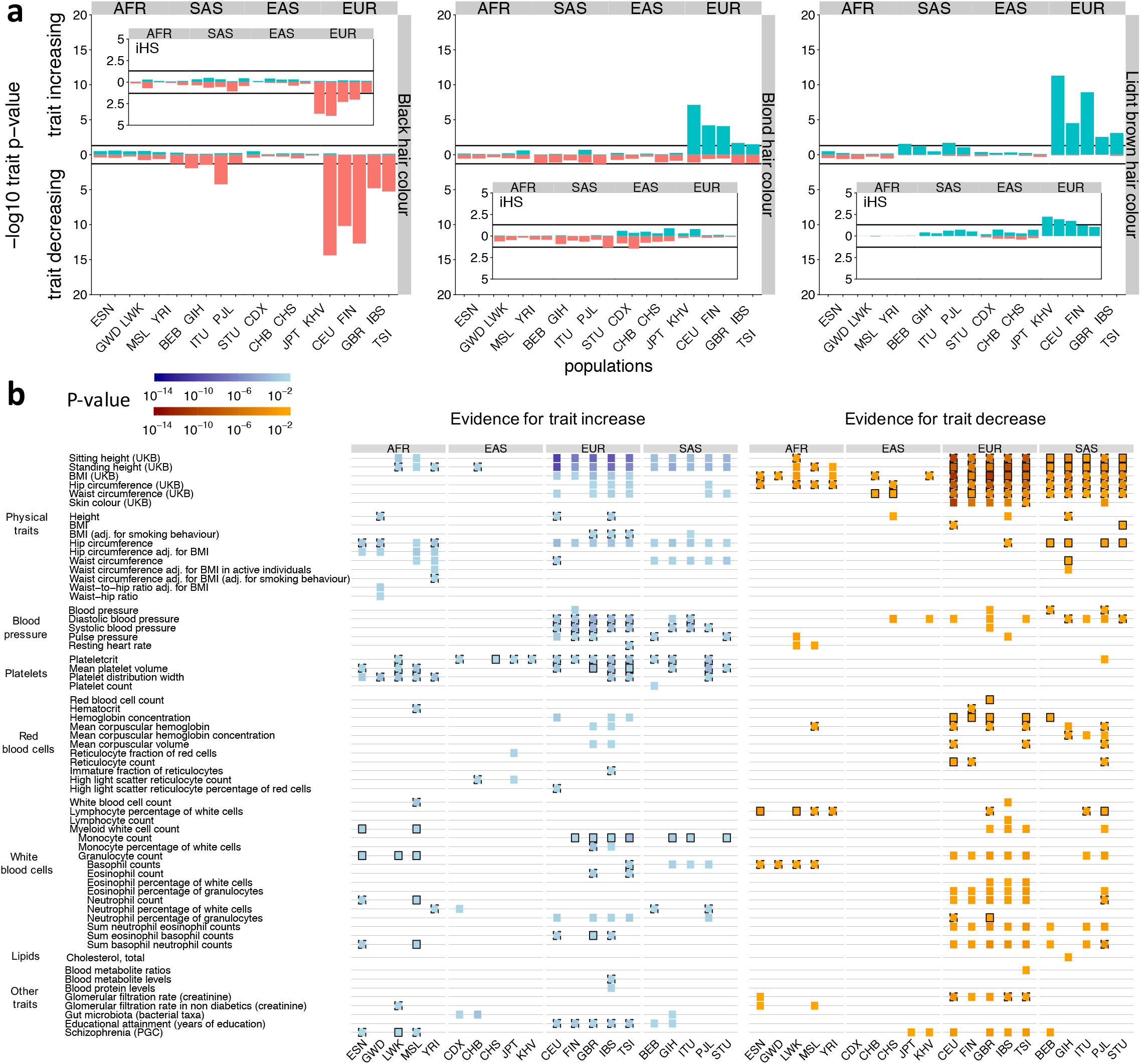
Evidence of selection on traits: **a**, P-values for evidence of directional selection of black, blond, and light brown hair colour (see **Methods** for calculation of p-values). Insets show p-values for the same test but using iHS scores instead, where iHS scores are calculated for each population separately for any variant with a minor allele frequency >5% in that population. **b**, Evidence for directional or bidirectional selection on multi-allelic traits. Each trait is associated with at least 10 SNPs in both effect directions in each of the considered populations. We show evidence for a trait increasing over time (left) and evidence for a trait decreasing over time (right) if *p* ≤ 0.05. Black boundaries indicate consistency with an additional test that tests for shifts in the DAFs (solid: *p* ≤ 0.05, dashed: *p* ≤ 0.5, Methods).

Height, Body Mass Index (BMI), and Schizophrenia have been studied previously and show a large number of association signals^58^. While several studies have reported genetic differentiation between populations for these traits^59–61^, evidence for selection remains controversial^40,47–51,59,60,62^. It was recently reported that recent selection on increased height in Europeans has been overestimated and that estimates have been confounded by subtle population stratification^40,50,51^. Our test finds an enrichment of selection evidence for *both* effect directions for height, across most populations except East Asians, using the large collection of UK Biobank associations. DAFs tend to be larger towards the height-decreasing direction. This complex picture may be a consequence of both negative and positive selection acting on height, as well as pleiotropy; SNPs impacting other traits might also impact height. We also identify strong evidence of selection towards decreased BMI across all populations, with agreement of DAF shifts, indicative of directional selection. For both traits, we detect little evidence of selection using associations in the smaller GWAS catalogue collection. For Schizophrenia, we find evidence of selection towards decreased risk in Europeans, and some South and East Asian populations, while African populations show selection evidence towards a risk increase.

Overall, although we find selection evidence for a range of traits, we observe little overlap with traits identified in Ref. [49], which focuses on very recent selection specific to the British population. Among other phenotypes, we see selection evidence for a variety of blood-related phenotypes, with congruent DAF signals. In Europeans and some South Asian groups, we detect a strong signal towards increases in diastolic and systolic blood pressures, contrary to previous studies showing selection for decreased blood pressure in these groups^56,63^. We moreover find evidence for selection towards decreased hemoglobin concentration and other related traits, while platelet-related traits appear to be selected to increase across many populations.

Interestingly, we observe differences between the frequency conditioned selection signal and shift in DAF for some traits related to white blood cells (**Figure 6b**). For instance, we detect a signal towards increased granulocyte counts in African populations, but decreasing counts in some European and South Asian populations. While DAFs are strongly different (*p* < 0.003; Wilcoxon test) and in agreement with the effect direction for the African groups, DAFs remain slightly lower for SNPs associated with decreasing granulocyte counts even in Europe and South Asia. (*p* < 0.09 for EUR, *p* < 0.12 for SAS; one-sided Wilcoxon test). However, these DAFs for granulocyte-decreasing variants are increased relative to those in African groups (*p* < 0.0013; one-sided paired Wilcoxon test), while DAFs for granulocyte-increasing variants were not significantly different (*p* > 0.4; two-sided paired Wilcoxon test), so a possible resolution is that this trait (or a related trait) was selected to increase in the past, and has more recently been selected to decrease in some non-African groups.

## Discussion

We have developed a scalable method, Relate, for estimating genealogies genome-wide and demonstrated its accuracy, as well as utility, on a diverse set of applications, building histories for thousands of samples. In many of these applications, we have improved in accuracy or resolution on existing state-of-the-art methods, each of which have previously required separate analyses. Although we have focused here on data for humans, Relate should work equally well in other recombining species. A strength of approaches based on building such genealogies is that results are automatically self-consistent: all our inferences are derived from the same genealogy, making results across different applications easier to compare. We note that this approach is highly modular, in the sense that the methods developed for genealogy-based inference should be applicable regardless of the specific algorithm used for estimating marginal trees.

In our analysis of 1000 Genomes data, we provide several examples whereby Relate-based trees are able to capture evolutionary processes that are themselves evolving through time: “evolution of evolution”. Changes through time in mutation rates, population size, migration, and archaic admixture, are simultaneously inferred, as are population-specific signals of natural selection. Genealogies thus provide a powerful way to study these complex, interacting phenomena, and we believe studies of other evolutionarily and temporally dynamic processes – for example of evolution of recombination rates through time^64,65^ - will yield new insights.

Interpretation of our findings regarding natural selection requires some care. A strength of our Relate Selection Test to identifying candidates of ongoing selection is that it provides p-values, which are naturally calibrated, even if population sizes vary through time. In common with previous studies, we find a relatively small (<40) number of clear signals of strong, ongoing selection across multiple human populations. In contrast, we find a much larger collection of phenotypes where – based on published GWAS data – there is evidence that mutations influencing a phenotype in one direction or another show evidence of directional selection. These traits include BMI, blood pressure, and white and red blood cell counts, and more generally we see an enrichment of selection evidence at loci shown to associate with human phenotypes in GWAS studies. While these findings appear highly consistent with the polygenic nature of most human phenotypes - which are expected to impose very weak selection, but on a large collection of loci^41^ - it remains challenging to assign selection signals to specific phenotypes. For example, a directional signal might be partly driven by selection on other phenotypes correlated to those studied. Moreover, even if mutations e.g. increasing WBC counts have been generally favoured in a group, this does not imply that WBC count *itself* has increased evolutionarily; if for example a selective sweep has fixed a single SNP of major effect on this phenotype (such as Duffy negativity in Africa, associated both with malaria resistance and decreased WBC count^66^), then selection might be acting on other SNPs to compensate this change. Environmental influences might have similar impacts. Differences between populations must also be interpreted carefully: aside from impacts of demographic history, most human GWAS’s to date have been conducted in European populations, so that recently arisen phenotype-influencing mutations in other groups might not have been observed, reducing power in those populations. Finally, we note that we only utilise the direction of association signals in testing for selection evidence, and test derived mutations, in order to increase robustness to residual population stratification still present in a GWAS, even after attempts to correct for such stratification. We believe that this is likely to resolve the most serious known issues, except in a setting where residual stratification (which can correlate with selection evidence^67^) improves power to observe effects that are genome-wide significant in one direction vs. another. Implicit in our approach is the idea that stratification issues are relatively far weaker for potentially genome-wide significant SNPs (of relatively large effect size) compared to directly using effect size estimates - which may be comparable to the strength of bias – across many or all SNPs genome-wide.

The fact that Relate is able to provide age estimates for mutations and other events in the tree is central, because these estimates enable us to construct initial statistics to understand ancient migration and admixture events, as well as evidence for natural selection either on individual mutations or collections of mutations. We note that it is important to account for variation in past population sizes (Supplementary Figure 3) for accurate age estimation. We regard the selection statistics applied here as initial approaches along a path towards a richer inference framework; it should be straightforward to develop related approaches to target e.g. background selection, full selective sweeps, or balancing selection. Because trees allow the spread of individual mutations to be inferred through time, more sophisticated approaches (e.g., Ref. [40]) should be able to examine temporally fluctuating selection, for example by using statistics similar to those we have introduced, but only testing for more rapid spread of particular mutations from some chosen time onwards. Another important direction for future work will be the development of techniques to understand migration events and ancient admixture. As one example, our results suggest a large impact of ancient substructure and/or archaic admixture specific to African populations, as has been previously hypothesized^36,37^. More generally, we believe that by following particular lineages, it should be possible to gain additional information (beyond e.g. cross-coalescence rates that we presented here) on the direction of past migration events. In principle, these analyses and those of selection could be done using the trees already constructed: we hope that methods will be developed providing tools to perform statistical analyses on a set of trees generated either by Relate, or other approaches. Other analyses might use estimated mutational ages obtained here directly, for example in understanding the properties of mutations influencing human disease^58^.

There are several natural extensions to the tree-building algorithm itself. A particularly useful extension might be allowing for increasing sample sizes. We note that a different approach to ours, tsinfer, has recently been developed^68^. This method has impressive scaling with sample size, and might readily extend to even millions of samples, while Relate can currently only handle at most a few tens of thousands of samples genome-wide. While tsinfer currently only infers tree topologies (as part of a full ancestral recombination graph structure), and so cannot infer tree times or model varying population sizes through time, it would be possible to use tsinfer-based tree topologies in our framework, allowing full tree-based inference for huge sample sizes, and this – or another approach to achieve a similar scale-up - represents an important direction for future work. Additionally, it should be possible to extend Relate to incorporate ancient DNA sequences, in order to leverage direct observation of ancient haplotypes. One complexity here is that such samples may have substantially higher error rates or more missing data than modern-day individuals, and an approach to handle this might involve “threading” of additional (ancient) sequences through genealogies initially built using sequenced individuals^2^. Such an approach might also be useful for efficient statistical phasing and/or imputation of individuals only typed at a subset of markers.

## Acknowledgements

We thank Nick Barton, Molly Przeworski, Guy Sella, Jonathan Terhorst, Pier Palamara, Gerton Lunter, Jonathan Marchini, Sile Hu, Christopher B. Cole, Thaddeus Aid, Clare E. West for helpful comments, ideas, and suggestions. L.S. acknowledges the support provided through the Engineering and Physical Sciences Research Council (EPSRC) [grant number EP/G03706X/1]. M.F. acknowledges the support provided through the Natural Sciences and Engineering Research Council of Canada (NSERC, PGS D) and the Clarendon Scholarship. S.R.M. acknowledges the support provided by the Wellcome Trust Investigator Award [grant number 098387/Z/12/Z and 212284/Z/18/Z]. This work was funded by the Wellcome Trust (WT090532/Z/09/Z).

## Author contributions

S.R.M. designed the study. L.S. and S.R.M. developed Relate with contributions by M.F. in the development of the algorithm for estimating coalescence rates. L.S. and S.R.M. performed the analysis, S.S. provided supplementary data and L.S. and S.R.M. wrote the manuscript.

## Data and software availability

The software Relate can be downloaded from https://myersgroup.github.io/relate under an Academic Use Licence.

## Datasets used in the current study were obtained from the following URLs

1000 Genomes Project phased dataset, https://mathgen.stats.ox.ac.uk/impute/1000GP_Phase3.html (13 Jan 2017); Genomic mask, ftp://ftp.1000genomes.ebi.ac.uk/vol1/ftp/release/20130502/supporting/accessible_genome_masks/ (20 Jul 2017); Human ancestral genome, ftp://ftp.1000genomes.ebi.ac.uk/vol1/ftp/phase1/analysis_results/supporting/ancestral_alignments/ (20 Jul 2017); GWAS catalogue, https://www.ebi.ac.uk/gwas/api/search/downloads/full (9 Nov 2017); PGC GWAS study, https://www.med.unc.edu/pgc/results-and-downloads (23 Nov 2018); HaploReg, http://archive.broadinstitute.org/mammals/haploreg/data/haploreg_v4.0_20151021.vcf.gz (21 Oct 2017); GTExeQTL https://storage.googleapis.com/gtex_analysis_v7/single_tissue_eqtl_data/GTEx_Analysis_v7_eQTL.tar.gz (13 Jan 2019); UK Biobank GWAS summary statistics, http://www.nealelab.is/uk-biobank (4 Oct 2018); PopHumanScan, https://pophumanscan.uab.cat (13 Jan 2019)

## External software used in the current study were downloaded from the following URLs

ARGweaver, https://github.com/mdrasmus/argweaver (24 Jan 2017);RENT+, https://github.com/SajadMirzaei/RentPlus (2 Oct 2017); msprime, https://github.com/tskit-dev/msprime (22 Jul 2017); msmc, https://github.com/stschiff/msmc2 (14 Oct 2017); SMC++, https://github.com/popgenmethods/smcpp (14 Oct 2017); simuPOP, http://simupop.sourceforge.net/ (27 Jun 2018); mbs, http://www.sendou.soken.ac.jp/esb/innan/InnanLab/ (27 Jun 2018); SDS, https://github.com/yairf/SDS (27 Jun 2018), selscan, https://github.com/szpiech/selscan (31 Jul 2018); hapbin, https://github.com/evotools/hapbin (11 Dec 2018)

## Methods

### Relate overview

We estimate genealogies as a sequence of rooted binary trees, where each tree captures the genealogy for a subregion of the genome. This representation serves as an approximation of an Ancestral Recombination Graph (ARG)^4^. We estimate local ancestry without global constraints on tree topology, thereby transforming genealogy reconstruction into a feasible and highly parallelisable problem.

Our approach can be divided roughly into three steps, which we detail below (also see **Figure 1**, Supplementary Figure 1, and Supplementary Note: Method details).

### Calculating position specific distance matrices

While trees vary along the genome, our method heavily utilizes ancestry information from nearby SNPs to reconstruct the tree at a specific position. We achieve this by using a HMM similar to that first proposed by Li and Stephens^27^ (see Supplementary Figure 2 for parameter choices). Intuitively, this HMM reconstructs a haplotype as a mosaic of other sample haplotypes along the genome (Supplementary Figure 1), allowing for mismatching in the copying process, and viewing changes in haplotype as recombination events. After applying the HMM, at a focal SNP *ℓ* each of the other haplotypes *j* therefore has some probability *p_ijℓ_* of being copied from, to generate haplotype *i*. After rescaling *p_ijℓ_* appropriately (see Supplementary Note: Method details), we obtain a position-specific distance matrix *d* whose entry (*i,j*) converges to the number of mutations derived in *i* and ancestral in *j* in the limit of no recombinations. In the presence of recombination, this *d* can be interpreted to store a local number of derived mutations, where more closely related haplotypes tend to have fewer mismatches over longer stretches, therefore receiving a smaller distance in this matrix.

We modified the Li-and-Stephens HMM to account for ancestral and derived states, a modification that guarantees our approach will construct the correct tree topology under the infinite-sites assumption with no recombination, while simultaneously speeding up the calculation of posterior copying probabilities.

### Tree builder

The distance matrix is turned into a binary tree using a hierarchical clustering algorithm. This hierarchical clustering algorithm is motivated by the observation that each row of the distance matrix should indicate the order in which this haplotype coalesced with other haplotypes of the dataset. This can be shown mathematically in some limit conditions, such as the case with no recombination (Supplementary Note: Method details).

Our algorithm iteratively merges clades of haplotypes, corresponding to past coalescences. After merging clades, we update the distance matrix by combining the corresponding rows and columns using a weighted sum, with weights determined by the size of clades. In each step of the algorithm, we merge the pair of clades that coalesce with each other before coalescing with any other clade, as determined using rows of the distance matrix. If multiple pairs of clades satisfy this condition, we choose the pair with minimum symmetrised score in the distance matrix. If the data are consistent with a binary tree under the infinite-sites model, such a pair always exists. In practise, errors in the data, complex recombination histories, or violations of assumptions made by our model, may result in a distance matrix that is inconsistent with a binary tree. To be robust to such cases, we relax the conditions for identifying pairs of clades to coalesce.

### Mapping mutations to branches and estimating branch lengths

Once tree topology is estimated as above, where possible we map mutations to the (unique) branch that has the identical descendants as the carriers of the derived allele in the data. To be robust to errors, where necessary we use an approximate rule for such mapping; however some mutations, e.g. repeat mutations or error-prone loci, may still not map to a unique branch. For these loci, we determine the smallest collection of branches, such that the data can be fully recovered. If a mutation maps to the tree only after reinterpreting the derived allele as the ancestral allele (and vice versa), we “flip” ancestral and derived alleles at this locus. For computation efficiency, to avoid having to construct a new tree at every locus we construct trees starting at the 5’ end of a region or chromosome, and move along the region constructing a new tree only when a SNP is flipped or cannot be mapped to a unique branch. Finally, we apply a Metropolis-Hastings type Markov Chain Monte Carlo (MCMC) algorithm to estimate branch lengths. The MCMC algorithm has a coalescent prior assuming a single panmictic population^3^.

### Estimating coalescence rates through time

We estimate the effective population size, defined as the inverse of the coalescence rate, by applying the following iterative algorithm. We initially estimate branch lengths using a constant effective population size. We then calculate a maximum-likelihood estimate of the coalescence rates between pairs of haplotypes given the branch lengths (Supplementary Note: Method details). By averaging coalescence rates over all pairs of haplotypes and taking the inverse, we obtain a population-wide estimate of the effective population size. We then use this population size estimate to re-estimate branch lengths, which requires only the final MCMC step of the branch-length estimation. By repeating these two steps until convergence (in practice, we use only 5 iterations as this provides good performance), we obtain a self-contained algorithm for jointly estimating branch lengths and the effective population size. We can average pairwise coalescence rates in different ways to obtain rates for sub-populations and cross-coalescence rates between populations.

### Pre-processing of the 1000 Genomes Project data set

The 1000 Genomes Project data set comprises 2504 individuals, from 26 populations. We obtained a phased version of the data set (**URLs**). We next excluded multi-allelic SNPs, and we exclude one individual (two haplotypes) from each population for future applications, and analysed the remaining 2478 individuals (Supplementary Table 2). We use a genomic mask provided with the 1000 Genomes Project dataset (**URLs**) to exclude regions in the marked as ‘‘not passing” in the pilot mask, to remove loci with low certainty of genotypes. We also exclude any base for which the fraction of ‘‘not passing” bases within 1000 bases to either side exceeds 0.9. To account for this filtering, we readjust the number of bases between SNPs at which we could have potentially observed a SNP. We use an estimate of the human ancestral genome (**URLs**) to identify the most likely ancestral allele for each SNP.

### Identifying branches indicative of Neanderthal and Denisovan introgression

We use genome sequences of the Vindija^22^ and Altai^33^ Neanderthals (NEA), and a Denisovan (DEN)^32^ to identify branches indicative of Neanderthal and Denisovan introgression into non-African populations. To identify branches that remain segregated from other human lineages for a long time, we use the world-wide genealogy of 2487 samples. To identify whether a branch is shared with NEA or DEN, at least one mutation needs to be mapped to that branch. We therefore exclude any mutation that has not been genotyped (or does not pass the genomic masks) in these ancient genomes. We further restrict our analysis to branches with at least two mutations mapped to them, as well as having an upper end that is older than 1M YBP. Of any SNPs that map to such branches, we calculate the fraction of SNPs that map to a branch with at least one NEA or DEN mutation. In **Figure 4c**, we plot these fractions as functions of the lower coalescent age of the branch onto which the SNP is mapped. Because the same branch may persist over multiple trees, we identify equivalent branches (Supplementary Note: Method details) and average ages of lower and upper ends across these equivalent branches. We assign a SNP to a population if at least one haplotype of that population carries the derived allele.

In **Figure 4d**, we observe an enrichment of branches indicative of introgression. This enrichment is identified by comparing the observed number of mutations in bins divided by upper and lower coalescent age to that expected in a panmictic history. To calculate the expected number of mutations in each bin, we fix the ages of coalescence events in each tree but randomise the topology assuming a panmictic population. The probability of upper and lower coalescent ages falling into bins *s* and *r*, conditional on the mutation arising while *k* lineages remain, is given by 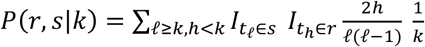 where *I* denotes the indicator function.

Assuming neutrality, a mutation is equally likely to have arisen anywhere on the branch it maps to. We therefore calculate the weighted average 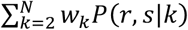, with weights *w_k_* defined as the proportions of a branch while *k* lineages remain. Summing this over all SNPs yields the expected number of mutations with upper and lower coalescent age falling, respectively, into bins *s* and *r*. In **Figure 4d**, age bins are defined by [−∞, 4.25), [4.25,4.75), [4.75,5.25), [5.25,5.75), [5.75, ∞).

### Tree-based statistic for detecting positive selection

Positive selection is expected to result in favourable mutations spreading rapidly in a population. One approach to capture this is via the number of lineages ultimately descending from the potentially favourable mutation(s): although we note that this is not the maximum likelihood approach, it has the benefit of making calculations straightforward. Under a null model of the standard coalescent model without selection, it is known that while *k* lineages remain, the joint distribution of the number of descendants of these *k* lineages is uniform in the partitions of *N* haplotypes to *k* lineages (see e.g., Ref. [69]). Using this property, we analytically calculate the marginal distribution that two of *k* lineages have more than *f_N_* descendants, where *f_N_* is the present-day DAF of the mutation. Here, we choose *k* to be the number of lineages remaining when the mutation of interest increased from frequency 1 to 2 (see Supplementary Note: Method details for the mathematical details).

To remove false-positive selection hits due to poorly inferred genealogies, our analysis for the 1000 Genomes Project data set is based on a subset of all SNPs mapping to trees, and present in 3 or more copies in the dataset. Specifically we remove SNPs failing any of the following filters: (i) the number of mutations mapping to that SNP’s tree is in the bottom 5^th^ percentile, or (ii) the fraction of tree branches having at least one SNP is in the bottom 5^th^ percentile. This excludes approximately 7% of SNPs.

### Simulation of positive selection

To simulate positive natural selection, we adopt the pipeline outlined in Ref. [49]. We first simulate the trajectory of the DAF using simuPOP^70^. We vary the selection coefficient between *s* = 0.001 and *s* = 0.05 and assume that the selected allele is beneficial throughout its history. We fix the present-day DAF to 0.7 (see Supplementary Figure 4 for other present-day DAFs). We then use mbs2^71^ (mutation rate *μ* = 1.25 × 10^−8^, constant recombination rate *ρ* = 5 × 10^−9^) to simulate a region of 20Mb, given the DAF trajectory for the central selected SNP. For each non-zero selection coefficient, we perform 200 simulations, and we perform 500 simulations for the neutral case. We assume a population size history as for our estimates for YRI and GBR, in separate simulations.

We compare to iHS, SDS, and a tree-based variant of SDS (trSDS) proposed in Ref. [40]. For iHS, SDS, and trSDS, we standardise scores using the mean and standard deviation in the neutral case, which is an idealised setting that should favour the power estimates of these methods. We then determine a critical standardised score that corresponds to a given type I error rate in the neutral case to estimate the statistical power. For Relate, we use frequency-conditioned p-values, by calculating a critical p-value that yields the desired false-positive rate in the neutral case (for the statistical power using raw p-values, see Supplementary Figure 4).

### Enrichment of SNPs with functional annotation among targets of positive selection

We merge selection evidence for SNPs by region (AFR: Africans, EAS: East Asians, EUR: Europeans, SAS: South Asians) by first calculating Z-scores of the logarithm of selection p-values within populations, and then averaging these Z-scores across populations. We exclude groups expected to be highly admixed^72^ (ACB, ASW, CLM, MXL, PEL, PUR (Supplementary Table 2)), because recent admixture may confound selection signals. We further exclude SNPs with a DAF <5% in the region of interest.

To assess statistical significance for the observed enrichment of GWAS hits and functional mutations in groups of SNPs showing evidence of selection, we used a block bootstrap with a block size of 1Mb. This will account for LD at scales below this threshold. In each bootstrap iteration, we resample blocks containing SNPs with a selection Z-score within the range of interest, with replacement, and calculate the ratio of the number of SNPs with functional annotation obtained using the HaploReg database^73^ (**URLs**) and the GWAS catalogue to the expected number of such SNPs, conditional on DAF. We condition on frequency, to account for the possibility that skewed frequency spectra in functional SNPs could be driving the signal.

### Pre-processing of GWAS

We use SNP-trait associations documented in the GWAS catalogue^74^ (**URLs**) to study polygenic adaptation. We use only association signals whose GWAS p-value is smaller than 5 × 10^−8^. For each trait, we also remove any duplicate SNPs.

For every combination of population, trait, and effect direction, we compile a set of approximately independent GWAS signals as follows.

For each pair of population and trait, we remove associations that are in close physical proximity and may therefore be in linkage disequilibrium (LD). For this, we first group SNPs into approximately independent blocks, such that any two GWAS hits in separate blocks are separated by at least 100kb and there are no intervals larger than 100kb with no GWAS hit inside a block. We then choose one GWAS hit from each block uniformly at random. We remove any SNP with a DAF <5%. To determine the effect direction of a SNP, we use the annotation in column “95% CI (TEXT)” combined with the indicated risk allele. We then realign the effect direction to the derived allele. We only consider SNPs for which an effect direction can be determined with this procedure. As described in the main text, we only analyse traits with at least 10 independent hits in both effect directions in all populations. This results in 76 traits and a total of 7302 GWAS hits (before filtering for SNPs in close proximity in each population).

For Schizophrenia, we are unable to obtain an effect direction using the procedure described above. Instead, we downloaded results for a large-scale GWAS conducted by the Psychiatric Genomics Consortium^75^. We considered SNPs reaching a GWAS p-value of 5 × 10^−8^ of which there were 9138. We intersected this set of SNPs with SNPs segregating in each of the considered populations. As for the GWAS catalogue, we identified approximately independent blocks. We then chose the SNP with lowest GWAS p-value in each block, resulting in 81 to 89 hits per population.

In addition, we use GWAS conducted as part of the UK Biobank^55^, focussing on highly polygenic physical traits. Our pre-processing protocol is analogous to that for schizophrenia detailed above. The number of approximately independent hits per population range from 272 hits for waist circumference to 989 hits for standing height.

### Trait selection test

For every combination of population, trait, and effect direction, we test whether p-values are smaller than expected. For this test, we first sample SNPs that we use for comparison. For each SNP associated with the population, trait, and effect direction tuple of interest, we sample 20 SNPs uniformly at random with replacement from SNPs, with the same present-day DAF in the population of interest. We then use a one-sided Wilcoxon rank-sum test to test whether the p-values of SNPs associated with tuple of interest tend to be smaller than those for the frequency-matched set of SNPs. We repeat this test 20 times and report the mean p-value of the Wilcoxon rank-sum test.

Our primary test identifies selection evidence conditional on DAF. However shifts in DAF can themselves serve as orthogonal evidence of polygenic adaptation, complementing our inferences. Therefore, we conducted a one-sided Wilcoxon rank-sum test to test whether DAFs of SNPs associated with the effect direction with selection evidence tend to exceed those associated with the opposing effect direction, and compared to our results conditional on SNP frequency. We note that we expect to lack power to reliably detect selection with this test, given that there are typically only tens of SNPs independently associating with each trait In addition, the relationship between selection and SNP frequencies can be complex if selection strength varies through time and/or geographic locations.

