## Supplementary Information for "A method for genome-wide genealogy estimation for thousands of samples"

February 14, 2019

2 Number of 1000 Genomes Project samples used in our analysis by population label. . .
3 Genome-wide significant hits for positive selection. . . . . . . . . . . . . . . . . . . . . . .

### Supplementary Note: Method details

#### 1 Overview of Relate

We first describe Relate in the absence of recombination. In this case, a single tree describes the genealogy of the whole genome. We define the number of *derived* mutations $d(i,j)$ as the number of mutations carried by haplotype $i$ and not by haplotype $j$. Notice that $d(i,j) \neq d(j,i)$. Assuming that every mutation happened exactly once, we can determine the order in which haplotype $i$ coalesced with other haplotypes by ordering them in ascending order of derived mutations $d(i,j)$ ($j = 1,\dots,i-1,i+1,\dots,N$). Once we know the relative order of coalescences, we reconstruct the tree topology using the hierarchical clustering algorithm described in Section 2. The resulting tree topology is guaranteed to be consistent with the truth in the sense that the constructed tree is a subtree of the gene tree describing the data (see Section 2.3).

In the presence of recombination, the relative order of coalescences changes along the genome. We apply a modified version of a Li and Stephens type hidden Markov model (HMM) [2] to calculate a local number of derived mutations $d(i,j;\ell)$ at every SNP $\ell$ (see Section 3). We modified the Li and Stephens model to take ancestral and derived states into account, which is necessary for $d(i,j;\ell)$ to converge to $d(i,j)$ in the limit of no recombination. We use $d(i,j;\ell)$ to reorder haplotypes at every SNP and apply the tree building algorithm to reestimate tree topology. Our method builds trees that are consistent with the truth if $d(i,j;\ell)$ orders haplotypes correctly. This is guaranteed for a recombination map consisting of zero and infinite recombination rates, which can be seen as a limit case of a hotspot recombination map (Section 2.3).

It is computationally inefficient, but possible, to reestimate tree topology at every SNP, because trees are unchanged if no recombination event occurred between SNPs. Instead, Relate initially estimates the tree topology at the first SNP of the 5' end of a chromosome. It then only reestimates the tree topology if a mutation cannot be uniquely *mapped* to a branch of the tree describing the previous SNP or is potentially *flipped*. A mutation is mapped to the branch for which the descendants coincide with the carriers of the alternative allele. Such a branch exists as long as the mutation occurred exactly once in human history and the estimated tree topology is correct. To be robust to errors in the data and the inferred tree, we relax this requirement such that the descendants of the branch only have to approximately coincide with the carriers of the mutation (see Section 2.2 for details). A mutation if potentially flipped, if it maps to a branch only after reinterpreting non-carriers as carriers and vice versa. While this introduces a small bias in the placement of recombination points, we note that this bias does not propagate along the genome because marginal trees are constructed using the distance matrix for the genomic position at which tree topology is reestimated.

Finally, once tree topologies are estimated, we estimate the branch lengths of each tree using an MCMC approach with a coalescent prior (see Section 4). We developed an algorithm that jointly estimates coalescence rates and branch lengths if historical coalescence rates are unknown (see Section 5).

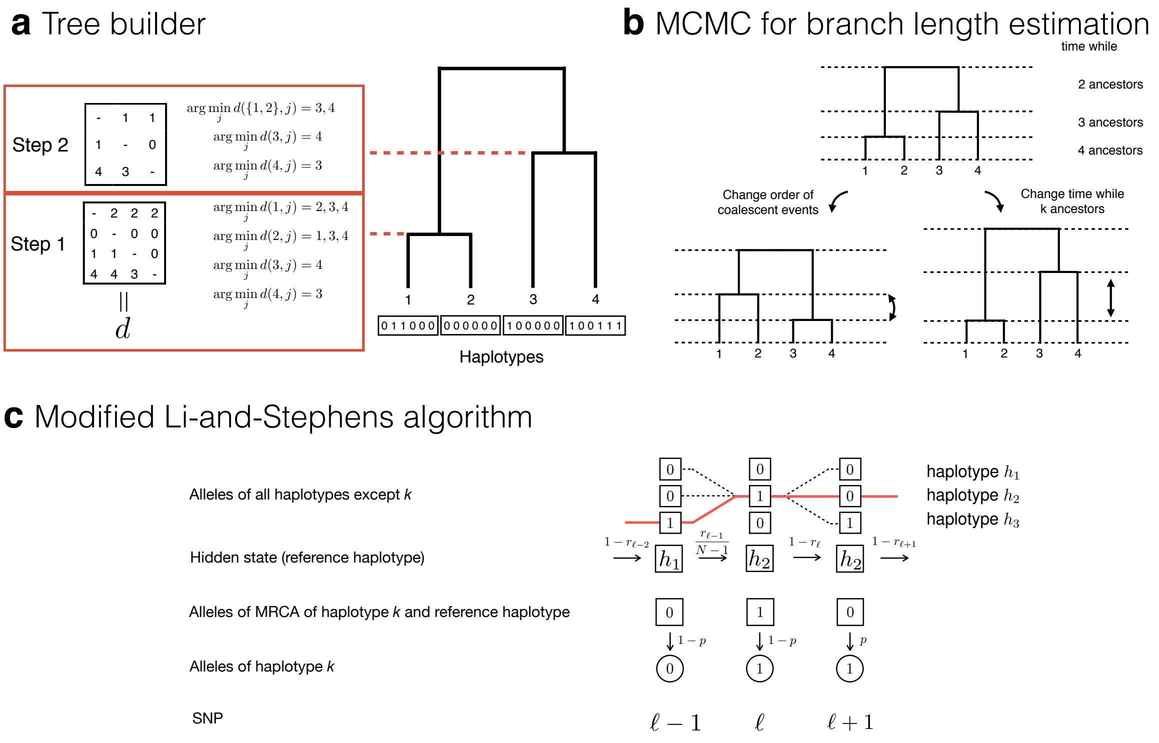

**Supplementary Figure 1: Schematics of the tree builder, branch length estimator, and modified Li-and-Stephens algorithm.**
**a**, Schematic of the hierarchical clustering algorithm for estimating tree topology. In the case of no recombination, the algorithm obtains matrix $d$ containing the number of derived mutations as input. Row $i$ of this matrix determines the order in which haplotype $i$ coalesced with other haplotypes. Using Eq. (1), the algorithm finds the pair that coalesces with each other before coalescing with any other sequence. In the example shown here, we can coalesce haplotypes 0 and 2 or haplotypes 3 and 4. We choose to coalesce haplotypes 1 and 2 first because the symmetrised distance is smaller for this pair, however this choice does not affect tree topology in this case. The resulting tree topology is consistent with the gene tree describing the data. In contrast, when the hierarchical clustering algorithm is applied to the symmetrised matrix $(d(i,j) + d(j,i))_{i,j=1,\dots,N}$, haplotypes 2 and 3 are coalesced first and the constructed tree topology is wrong. This is equivalent to applying the UPGMA algorithm to the symmetrised matrix of derived mutations [1]. **b**, Schematic of possible proposal moves in the MCMC algorithm for estimating branch lengths. We propose either a change in the order of coalescence events or a change in the time while $k$ ancestors remain. **c**, Schematic of the modified Li-and-Stephens algorithm applied to haplotype $k$, which has alleles 0, 1, 1 at loci $\ell - 1$, $\ell$, $\ell + 1$. The emission and transition probabilities shown correspond to the path indicated by the red solid line. At SNP $\ell - 1$, the reference haplotype is $h_1$ which has allele 1. Because the allele of haplotype $k$ is 0, the allele of the MRCA with $h_1$ is also 0 assuming that every mutation is unique in history. Therefore, the emission probability equals $1 - p$, where $p$ is the probability of a mutation. At SNP $\ell$, the reference haplotype has changed to $h_2$. The alleles of haplotype $k$ and $h_2$ are 1. Therefore, the MRCA has allele 1 and the emission probability is given by $1 - p$. At SNP $\ell + 1$, haplotype $k$ has allele 1. The allele of the reference haplotype $h_2$ is 0 and so is that of the MRCA, such that the emission probability equals $p$. Using this HMM, we calculate the likelihood $P_{\mathrm{m}}(H_\ell = j | D^{(k)})$. This is the likelihood of copying from reference haplotype $j$ at SNP $\ell$, conditional on observing $D^{(k)}$. We notice that $P_{\mathrm{m}}(H_\ell = j | D_\ell^{(k)})$ is obtained as the sum of all possible paths when $H_\ell = j$ is fixed (indicated by the dashed lines).

#### 2 Tree builder

##### 2.1 Hierarchical Clustering

We begin by describing how we construct a tree using a distance matrix $d = (d(i,j))_{1 \leq i,j \leq N}$ as input. We note that in general $d$ does not provide a distance metric, and $d(i,j) \neq d(j,i)$. A schematic is depicted in Supplementary Fig. 1**a**.

The tree builder is initialised by placing each haplotype in a separate cluster. The algorithm proceeds by finding pairs of clusters that coalesce with each other before coalescing with any other haplotype. Such a pair satisfies

$$\begin{aligned} \mathcal{A} &= \arg\min_{\mathcal{A}'} d(\mathcal{B}, \mathcal{A}') \\ \mathcal{B} &= \arg\min_{\mathcal{B}'} d(\mathcal{A}, \mathcal{B}'), \end{aligned} \tag{1}$$

where the distance $d(\mathcal{A}', \mathcal{B}')$ between two clusters $\mathcal{A}'$ and $\mathcal{B}'$ of cardinality $|\mathcal{A}'|$ and $|\mathcal{B}'|$, respectively, is given by

$$d(\mathcal{A}', \mathcal{B}') = \frac{1}{|\mathcal{A}'||\mathcal{B}'|} \sum_{x \in \mathcal{A}'} \sum_{y \in \mathcal{B}'} d(x,y). \tag{2}$$

There might be more than one pair satisfying Eq. (1), in which case we choose the pair with the smallest symmetrised distance $d(\mathcal{A}, \mathcal{B}) + d(\mathcal{B}, \mathcal{A})$. The chosen pair is then combined to a new cluster comprising all haplotypes of both clusters. The algorithm is terminated when all haplotypes are in one cluster.

A pair satisfying Eq. (1) is guaranteed to exist as long as there exists a tree consistent with the order of coalescence events implied by matrix $d$. Sometimes such a tree cannot be constructed. To make our algorithm robust to such situations, we replace Eq. (1) by

$$\begin{aligned} \mathcal{A} &\in \{\mathcal{A}' : |d(\mathcal{B}, \mathcal{A}') - \min_{\mathcal{C}} d(\mathcal{B}, \mathcal{C})| < \varepsilon\} \\ \mathcal{B} &\in \{\mathcal{B}' : |d(\mathcal{B}', \mathcal{A}) - \min_{\mathcal{C}} d(\mathcal{C}, \mathcal{A})| < \varepsilon\}, \end{aligned} \tag{3}$$

which allows for a tolerance $\varepsilon > 0$ in finding feasible pairs. In our implementation, we set $\varepsilon = 0.2$. In addition, in case pairs satisfying Eq. (3) cannot be found, we choose the pair with the smallest symmetrised distance.

##### 2.2 Deciding when to build a new tree

It is computationally inefficient, but possible, to reestimate tree topology at every SNP, because trees are unchanged if no recombination event occurred between SNPs. Instead, Relate initially estimates the tree topology at the first SNP of the 5' end of a chromosome. It then only reestimates the tree topology if a mutation cannot be uniquely *mapped* to a branch of the tree describing the previous SNP (or the mutation is potentially *flipped*, see below). While this introduces a small bias in the placement of recombination points, we note that this bias does not propagate along the genome because marginal trees are constructed using the distance matrix for the genomic position at which tree topology is reestimated.

We use present-day genome data of outgroups, such as chimpanzees and other primates for humans, to determine the ancestral and derived alleles. Occasionally, the ancestral allele can be confused with the alternative allele due to repeat mutations between species or sequencing errors. We can infer such cases if a SNP maps onto the tree only after non-carriers of the mutation are reinterpreted as carriers and vice versa. We refer to these SNPs as *flipped* SNPs and reinfer tree topology whenever we detect a potentially flipped SNP.

A mutation is mapped to the branch for which the descendants coincide with the carriers of the derived allele. Such a branch exists as long as the mutation occurred exactly once in human history and the estimated tree topology is correct. To be robust to errors in the data and the inferred tree, we relax this requirement as follows.

By placing a mutation on a branch, we indicate that all descendants below that branch carry the mutation. Let us denote the set of haplotypes that carry the mutation by $C_\mathrm{t}$ and the set of haplotypes that do not carry the mutation by $N_\mathrm{t}$. Similarly, let us denote the set of haplotypes that (do not) carry the mutation in the data set by $C_\mathrm{d}$ and $N_\mathrm{d}$. For a mutation to map to a branch, it needs to satisfy

$$\frac{|C_\mathrm{t} \cap C_\mathrm{d}|}{\max\{|C_\mathrm{t}|, |C_\mathrm{d}|\}} > 0.7$$
$$\frac{|N_\mathrm{t} \cap N_\mathrm{d}|}{\max\{|N_\mathrm{t}|, |N_\mathrm{d}|\}} > 0.7. \tag{4}$$

These conditions should identify suitable candidate branches and should prevent mapping infrequent mutations, such as doubletons, to a unique branch, when in fact they cannot have arisen by a single mutation. Out of all remaining candidate branches, we calculate the fraction of missclassified haplotypes given by

$$\frac{|N_\mathrm{t} \cap C_\mathrm{d}| + |C_\mathrm{t} \cap N_\mathrm{d}|}{N}. \tag{5}$$

We also calculate the same quantity for branches that satisfy Eq. (4) after reinterpreting carriers as non-carriers and vice-versa. We then accept the branch with the minimum score given by Eq. (5) if it is also less than 0.03. We only flip a SNP if this leads to a smaller score (i.e., in case of a tie, we do not flip the SNP). These rules, though heuristic, allow approximate mapping for mutations in 4 or more copies in the data set (Supplementary Fig. 2a).

If such unique branch cannot be found, we map the mutation to more than one branch. In this case, we find the smallest set of branches, such that all carriers of the mutation $C_\mathrm{d}$ are below one chosen branch and such that the summed score given by Eq. (5) equals zero. We do the same after flipping the SNP. We choose to flip the SNP only if this leads to a smaller set of branches. We then add one over the number of branches chosen to the number of mutations on that branch. For many analyses, we only consider mutations mapping to a unique branch, however we note that e.g., CpG mutations often occur on multiple branches.

##### 2.3 Consistency of estimated tree topology in the absence of recombination

We prove that the use of the number of derived mutations $d(i, j)$ as a distance matrix always guarantees estimation of a tree topology consistent with the truth, assuming that every mutation is unique in history, and no recombination. We recall that we defined the number of derived mutations $d(i, j)$ as the number of mutations carried by haplotype $i$ and not by haplotype $j$. We assume that there is a gene tree that is consistent with SNPs in the data in the sense that every SNP can be mapped to a unique branch of the gene tree [3]. Such a gene tree contains polytomies reflecting branches unresolved by observed mutations, and always exists and is unique assuming the infinite-sites model. We say, that a binary coalescence (sub-)tree, is consistent with a gene tree if all carriers of any mutation coalesce before non-carriers of a mutation.

We prove that our tree builder described in Section 2 constructs a tree that is consistent with the gene tree. The input matrix is the matrix of derived mutations $(d(i, j))_{i,j=1,\dots,N}$ and we set $\varepsilon = 0$ in the tree builder. The tree builder therefore coalesces haplotypes according to Eq. (1).

**Proposition 2.1.** *The tree builder cannot coalesce carriers and non-carriers before it has coalesced all carriers of any SNP.*

*Proof.* Assume without loss of generality (w.l.o.g.) that $1, \dots, k$ are carriers of a SNP and $k+1, \dots, N$ are non-carriers of the same SNP. We first observe that if a branch in the gene tree has a mutation, all

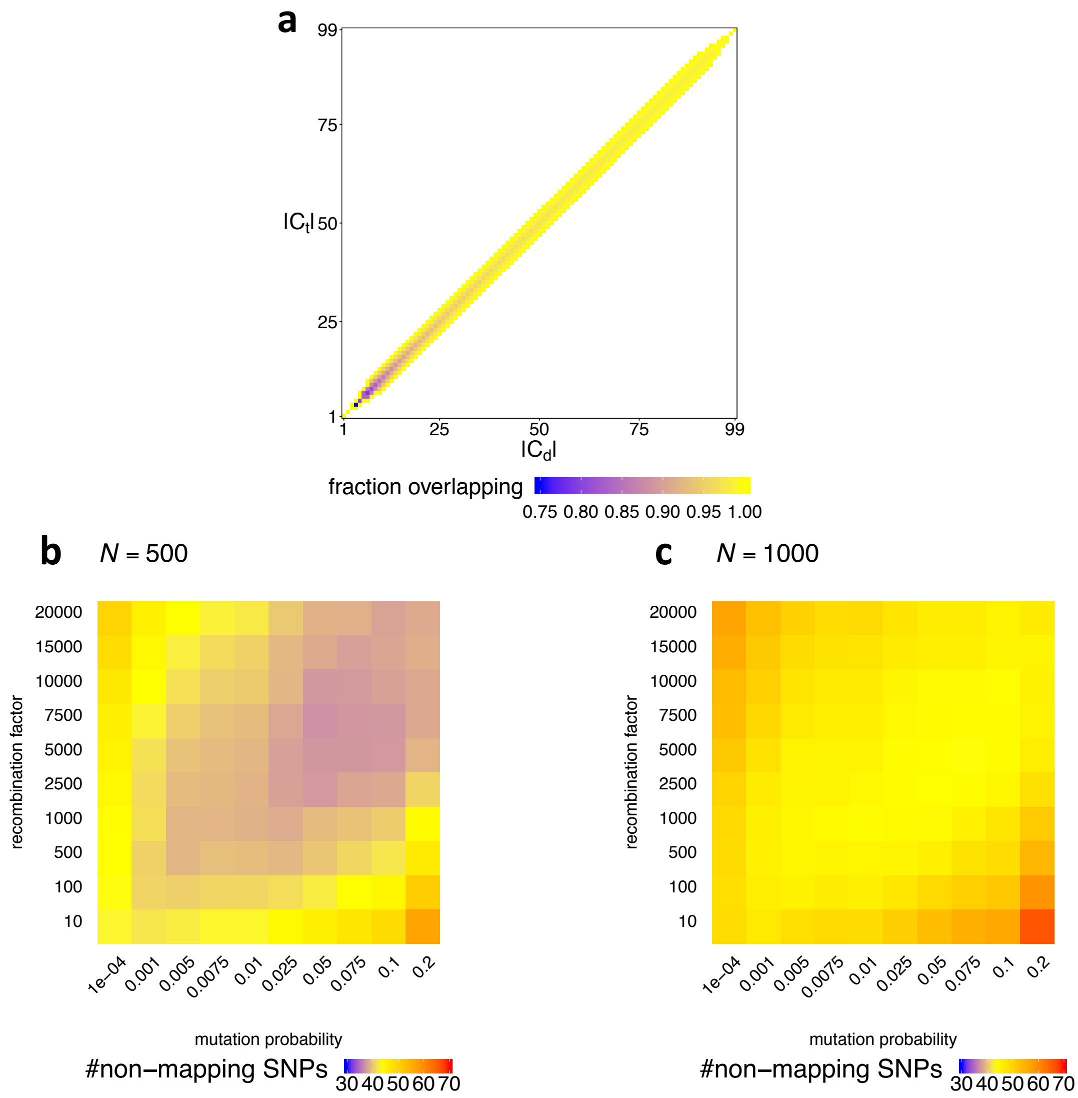

**Supplementary Figure 2: Mapping rule for mutations and sensitivity of the modified Li-and-Stephens algorithm to parameter choice.**
**a** Heatmap showing the necessary and sufficient overlap between the set of descendants of a branch ($C_{\mathrm{t}}$) and the set of carriers of the derived allele ($C_{\mathrm{d}}$), given $|C_{\mathrm{t}}|$ and $|C_{\mathrm{d}}|$, with $N = 100$, as determined by Eqs. (4) and (5). Colours show $|C_{\mathrm{t}} \cap C_{\mathrm{d}}| / \min\{|C_{\mathrm{t}}|, |C_{\mathrm{d}}|\}$, where white indicates that a mutation can never be mapped for the corresponding combination of $|C_{\mathrm{t}}|$ and $|C_{\mathrm{d}}|$. b, c, Number of non-mapping SNPs for different values of $p$ (horizontal axis) and $R$ (vertical axis) for $N = 500$ (**a**) and $N = 1000$ (**b**). The subsets of haplotypes are chosen uniformly at random from all haplotypes. We calculated the mean over 50 randomly chosen subregions of length 1200 SNPs on chromosome 20. In our implementation, we fixed $p = 0.025$ and $R = 2500$.

descendants of that branch carry at least one more derived mutation to any non-carrier of the mutation than to a carrier of the mutation. Using the this observation, we obtain

$$d(i,j) < d(i,h) \text{ for } i,j \in \{1,\ldots,k\} \text{ and } h \in \{k+1,\ldots,N\}. \tag{6}$$

This propoerty is why it is important to distinguish derived mutations in determining distances; the equivalent to Eq. (6) does not hold if $d(i,j)$ is determined as the number of differences between sequences $i$ and $j$.

It follows that coalescing a carrier and a non-carrier in the first step of the algorithm is not feasible in Eq. (1). Let us assume that we have not coalesced carriers and non-carriers until the $x$'th step of the algorithm and that there still exist more than one cluster of carriers. We prove that it is still not feasible to coalesce a cluster of carriers and non-carriers in the $x+1$st step. Let us denote by $\mathcal{A}$ and $\mathcal{B}$ clusters containing only carriers and by $\mathcal{C}$ a cluster containing only non-carriers. We obtain from Eq. (6),

$$d(\mathcal{A},\mathcal{B}) < d(\mathcal{A},\mathcal{C}), \tag{7}$$

because an average of $d(i,j)$ with $i,j \in \{1,\ldots,k\}$ is always smaller than an average of $d(i,h)$ with $i \in \{1,\ldots,k\}$ and $h \in \{k+1,\ldots,N\}$. Therefore coalescing $\mathcal{A}$ and $\mathcal{C}$ does not satisfy Eq. (1). □

With Proposition 2.1, we know that the tree builder never violates the gene tree and therefore, if the algorithm successfully coalesces all haplotypes, we obtain a tree that is consistent with the gene tree. It remains to prove that the tree builder can always find a next coalescence event and therefore, by induction, terminates with all haplotypes in one cluster.

**Proposition 2.2.** *Assuming there is a gene tree consistent with the data, the tree builder terminates with all haplotypes in one cluster.*

*Proof.* Assume that we are in the $x$th step of the tree builder. We prove that we can coalesce a pair of clusters to proceed to the $x+1$st step of the tree builder. We have already proved that the tree constructed until the $x$th step is consistent with the gene tree. We can therefore find two clusters $\mathcal{A}$ and $\mathcal{B}$, such that coalescing these clusters is consistent with the gene tree.

Let $\mathcal{C}$ be a third cluster distinct from $\mathcal{A}$ and $\mathcal{B}$. For $\mathcal{A}$ and $\mathcal{B}$ to satisfy Eq. (1), we require

$$\begin{aligned} &d(\mathcal{A},\mathcal{B}) \leq d(\mathcal{A},\mathcal{C}) \text{ and} \\ &d(\mathcal{B},\mathcal{A}) \leq d(\mathcal{B},\mathcal{C}). \end{aligned} \tag{8}$$

We show $d(\mathcal{A},\mathcal{B}) \leq d(\mathcal{A},\mathcal{C})$ and note that $d(\mathcal{B},\mathcal{A}) \leq d(\mathcal{B},\mathcal{C})$ can be shown analogously.

Let us define by $d_\ell(\mathcal{A},\mathcal{B})$ the distance considering only SNP $\ell$, such that $d(\mathcal{A},\mathcal{B}) = \sum_{\ell=1}^{L} d_\ell(\mathcal{A},\mathcal{B})$ with $L$ denoting the number of SNPs. We note that any SNP with no derived sequences in $\mathcal{A}$ yields $d_\ell(\mathcal{A},\mathcal{B}) = d_\ell(\mathcal{A},\mathcal{C}) = 0$. Let us therefore consider the possible types of SNPs with at least one derived mutation in $\mathcal{A}$. We note that we cannot have SNPs with both carriers and non-carriers in more than one cluster, because we would have violated Proposition 2.1. Therefore, the possible SNPs are as follows.

- If a SNP is only derived in sequences in $\mathcal{A}$, then $d_\ell(\mathcal{A},\mathcal{B}) = d_\ell(\mathcal{A},\mathcal{C})$.
- If a SNP is derived in all sequences of $\mathcal{A}$ and $\mathcal{B}$, but no sequences in $\mathcal{C}$, then

$$d_\ell(\mathcal{A},\mathcal{B}) = 0 < 1 = d_\ell(\mathcal{A},\mathcal{C}). \tag{9}$$

- If a SNP is derived in all sequences of $\mathcal{A}$ and $\mathcal{C}$, but no sequences of $\mathcal{B}$, then this SNP violates the assumption that coalescing $\mathcal{A}$ and $\mathcal{B}$ is consistent with the gene tree.
- If a SNP is derived in all sequences of $\mathcal{A}$, $\mathcal{B}$, and $\mathcal{C}$, then $d_\ell(\mathcal{A},\mathcal{B}) = d_\ell(\mathcal{A},\mathcal{C}) = 0$.

By summing over all SNPs, we obtain

$$d(\mathcal{A},\mathcal{B}) = \sum_{\ell=1}^{L} d_\ell(\mathcal{A},\mathcal{B}) \leq \sum_{\ell=1}^{L} d_\ell(\mathcal{A},\mathcal{C}) = \sum_{\ell=1}^{L} d(\mathcal{A},\mathcal{C}), \tag{10}$$

as required. □

We conclude with a few observations. First, we notice that all branches that have a mutation in the gene tree are guaranteed to exist in the tree built by our tree builder. In particular, if all branches in the gene tree have a mutation, the tree builder is guaranteed to build the correct tree. The tree builder therefore constructs the correct tree in the limit of an infinite mutation rate (and under the infinite-sites assumption). Second, we notice that if recombination rates are only either infinite or zero, we can divide the genome into regions of zero recombination rates. In this case, the tree builder constructs trees consistent with the gene trees of these regions. Finally, we notice that our proof did not depend on the absolute values of the input distance matrix, and that the only requirement was that rows of the distance matrix are perfectly correlated to the order of coalescences of a haplotype with other haplotypes. Therefore, we expect our tree builder to construct an accurate tree as long as rows of the distance matrix obtained from the modified Li-and-Stephens algorithm (Section 3) is well correlated with the order in which a haplotype coalescences with other haplotypes.

### 3 Calculating distance matrices

#### 3.1 Assumptions about the input data

We apply a version of the Li-and-Stephens algorithm to calculate distance matrices. To apply this algorithm, we assume haplotype SNP data as input, which can be inferred by phasing genotype data [4, 5]. We assume a high coverage of SNPs along the genome and no bias with respect to the frequency of a mutation in the population. We also assume that at most one mutation has occurred at any given position along the genome since the divergence of humans from chimpanzees and other primates. This assumption, known as the infinite sites model, is justified by a small average mutation rate in humans [6].

Additionally, we require knowledge of the ancestral allele for every recorded mutation. The ancestral allele can be determined by aligning the human genome to present day genomes of other primates [7]. Assuming that it is unlikely to observe a mutation at the same genomic position in humans and other primates, the allele carried by other primates is declared to be the ancestral allele. For humans, we use the ancestral genome that was inferred as part of the 1000 Genomes Project (see URLs in main text).

For the robustness of Relate to genotype errors and occasional confusion of ancestral and alternative alleles, please refer to the Supplementary Note: Simulations.

Under these assumptions, we can represent SNP data as a binary matrix $D$ with each row corresponding to one haplotype. The matrix $D$ therefore has dimensions $N \times L$, where $N$ is the number of haplotypes and $L$ is the number of SNPs. In this representation, a haplotype is a binary vector, where the ancestral allele is denoted by 0 and the alternative allele is denoted by 1. We denote row $i$ of $D$ by $D^{(i)}$.

#### 3.2 Modified Li-and-Stephens algorithm

A recombination event may change the order of coalescence events between haplotypes. Therefore, mutations close to the SNP of consideration are more informative than SNPs further away. To capture this, we apply a Li and Stephens type HMM [2]. We modified the original HMM by changing emission probabilities such that they take ancestral and derived states into account. A schematic of the HMM is depicted in Supplementary Fig. 1**c**.

The HMM can be interpreted as a generative model for haplotype $D^{(i)}$ using all other haplotypes as inputs. To generate the allele $D^{(i)}_\ell$ at SNP $\ell$, we first choose one reference haplotype $H_\ell$ from all other haplotypes. This reference haplotype is the hidden state of the HMM. In Ref. [2], the emission probabilities are defined by

$$P_{\mathrm{c}}(D^{(i)}_\ell | H_\ell = j) = \begin{cases} p & \text{if } D^{(i)}_\ell \neq D^{(j)}_\ell, \\ 1-p & \text{if } D^{(i)}_\ell = D^{(j)}_\ell. \end{cases} \tag{11}$$

Here, $p$ is the mismatch probability. In this conventional definition of the emission probabilities, a mutation may have occurred on the branch from $i$ to the MRCA with $j$, or on the branch from $j$ to the MRCA with $i$. Therefore, information about on which branch the mutation occurred is lost. To preserve this information, we change the emission probabilities to

$$P_{\mathrm{m}}(D^{(i)}_\ell | H_\ell = j) = \begin{cases} p & \text{if } D^{(i)}_\ell = 1, D^{(j)}_\ell = 0, \\ 1-p & \text{if } D^{(i)}_\ell = 0, D^{(j)}_\ell = 0, \\ 1-p & \text{if } D^{(i)}_\ell = 1, D^{(j)}_\ell = 1, \\ 1-p & \text{if } D^{(i)}_\ell = 0, D^{(j)}_\ell = 1. \end{cases} \tag{12}$$

We can interpret $p$ as the probability of a mutation since the MRCA. The emission probability equals $p$, if haplotype $i$ carries a mutation at site $\ell$ which is not carried by the reference haplotype $j$, such that the mutation must have occurred on the branch from $i$ to the MRCA with $j$. Otherwise, no mutation occurred on this branch and the emission probability equals $1-p$ (see Supplementary Fig. 1 **c**). The hidden state may change between neighbouring SNPs according to transition probabilities $r_\ell$. The transition probabilities are proportional to the recombination probabilities obtained from a recombination map. We describe how to choose $p$ and $r_\ell$ in Section 3.3.

Using the modified Li-and-Stephens algorithm, we can calculate a distance matrix at every SNP which we will use as input for the tree builder described in Section 2. We first derive a distance matrix for the case when transition probabilities are set to zero, which corresponds to no recombination. In this case, the reference haplotype remains the same along the genome. The likelihood of observing $D^{(i)}$ given reference haplotype $H_\ell = j$ is given by

$$P_{\mathrm{m}}(D^{(i)} | H_\ell = j) = p^{d(i,j)}(1-p)^{L-d(i,j)}, \tag{13}$$

where $d(i,j)$ is the number of derived mutations defined in Section 1. By taking logarithms on both sides of Eq. (13), we obtain

$$\log P_{\mathrm{m}}(D^{(i)} | H_\ell = j) = d(i,j) \log\left(\frac{p}{1-p}\right) + L\log(1-p). \tag{14}$$

By rearranging Eq. (14), we obtain

$$d(i,j) = \frac{\log P_{\mathrm{m}}(D^{(i)} | H_\ell = j) - L\log(1-p)}{\log\left(\frac{p}{1-p}\right)}. \tag{15}$$

We can generalise Eq. (15) to the case of non-zero recombination rates to define, for each SNP $\ell$, the *local* number of derived mutations

$$d(i,j;\ell) = \frac{\log P_{\mathrm{m}}(D^{(i)} | H_\ell = j) - L\log(1-p)}{\log\left(\frac{p}{1-p}\right)}. \tag{16}$$

Equation (16), corresponding to a local number of derived mutations, orders coalescence events locally at every SNP. In practice, we use $d(i,j;\ell) - \min_{j \neq i} d(i,j;\ell)$ as our distance matrix, which only affects our tie-breaking heuristic in case no pair of haplotypes (or clusters) satisfy Eq. (3) and we choose a pair with minimum entry in the symmetrised matrix. By subtracting the minimum entry from each row, we remove residuals interpretable as the number of derived mutations on the tip branches, which could confound the symmetrised distance if some haplotypes have many more mutations at their tips than other haplotypes. The quantity $\log P_{\mathrm{m}}(D^{(i)}|H_\ell = j)$ in Eq. (16) can be calculated using the forward-backward algorithm. We describe how to efficiently implement the Li-and-Stephens algorithm in Section 3.4.

We notice that with the conventional definition of the emission probabilities (Eq. (11)) and no recombination, we obtain

$$d(i,j) + d(j,i) = \frac{\log P_{\mathrm{c}}(D^{(i)}|H_\ell = j) - L\log(1-p)}{\log\left(\frac{p}{1-p}\right)}. \quad (17)$$

In this case, we obtain a symmetric matrix that stores the number of mutations that differ between two haplotypes. Using the matrix $d(i,j) + d(j,i)$ as input to our tree builder is equivalent to applying the UPGMA algorithm [1], which is an alternative hierarchical clustering algorithm, to the matrix $d(i,j) + d(j,i)$. We show in Supplementary Fig. 1**a**, how this can lead to estimation of a tree topology that is inconsistent with the data. Intuitively, this is because $d(i,j) + d(j,i)$ combines the number of mutations of two branches, such that information about the number of mutations on each branch is lost. Therefore, $d(i,j)$ preserves more information about the tree topology than $d(i,j) + d(j,i)$.

#### 3.3 Choosing parameters for the modified Li-and-Stephens algorithm

The transition probabilities in the HMM are determined by the recombination map, multiplied by a constant factor $R$. The mutation probability $p$ should reflect the true biological mutation rate and errors in the data set. In practice, we find that the tree topology constructed using distance matrix $d(i,j;\ell)$ (Eq. (16)) is very robust with respect to choice of $p$ and $R$. To illustrate this, we evaluate the performance of our algorithm for different choices of $p$ and $R$. In our implementation, we have fixed $p = 0.025$ and $R = 2500$.

To evaluate how well a pair $(p,R)$ captures ancestry information in a subregion of the genome, we sample subregions of lengths 1200 SNPs at random. We first apply the modified Li-and-Stephens algorithm on the chosen subregion. For every SNP $400 < \ell < 800$, we then do the following. We first calculate the distance matrix as described in Section 3. We modify the distance matrix by hiding the focal SNP $\ell$ which can be done by subtracting 1 from any entry $(i,j)$ where $i$ is a carrier and $j$ is not a carrier of the mutation. We then build the tree with this modified distance matrix using the hierarchical clustering algorithm described in Section 1 . Finally, we attempt to place the focal SNP on branches of the tree and record whether a branch exists such that the descendants of that branch coincide with carriers of the SNP.

We therefore evaluate a pair $(p,R)$ by how well we can build trees at focal SNPs using information from surrounding SNPs only. As $R$ becomes larger and $p$ becomes smaller, only SNPs close to the focal SNP will influence the distance matrix. As $R$ becomes smaller and $p$ becomes larger, SNPs further away will influence the distance matrix. We should find an optimal pair $(p,R)$ such that we include enough SNPs to be able to build a tree onto which the focal SNP can be mapped and such that we prevent SNPs that map onto different trees from distorting the signal.

We applied this method to 50 subregions of chromosome 20 of the 1000 Genomes Project data set. In Supplementary Fig. 2b and c, we show that the number of non-mapping SNPs is relatively robust to the choice of $(p,R)$, particularly along the axis $p/R = 10^{-6}$. We therefore fix $p = 0.025$ and $R = 2500$ in our implementation.

##### 3.4 Speed-up and approximation of the modified Li-and-Stephens algorithm

To calculate $d(i,j;\ell)$ defined in Eq. (16), we need to calculate $P_{\mathrm{m}}(D^{(i)}|H_\ell = j)$. We apply Bayes' theorem and obtain

$$P_{\mathrm{m}}(D^{(i)}|H_\ell = j) = \frac{P_{\mathrm{m}}(H_{\ell=j}|D^{(i)})P_{\mathrm{m}}(D^{(i)})}{P_{\mathrm{m}}(H_\ell = j)}. \tag{18}$$

We assume the prior probability of copying from any haplotype to be identical, such that $P_{\mathrm{m}}(H_\ell = j) = 1/(N-1)$, and calculate $P_{\mathrm{m}}(H_\ell = j|D^{(i)})$ using a forward-backward algorithm. The forward algorithm calculates $\alpha_j(\ell) = P_{\mathrm{m}}(H_\ell = j, D^{(i)}_{1:\ell})$ and the backward algorithm calculates $\beta_j(\ell) = P(D^{(i)}_{(\ell+1):L}|H_\ell = j)$ such that

$$P_{\mathrm{m}}(H_\ell = j|D^{(i)}) = \frac{\alpha_j(\ell)\beta_j(\ell)}{P_{\mathrm{m}}(D^{(i)})}. \tag{19}$$

By substituting Eq. (19) in Eq. (18) and taking logarithms, we obtain

$$\log P_{\mathrm{m}}(D^{(i)}|H_\ell = j) = \log\left(\alpha_j(\ell)\beta_j(\ell)\right) + \log(N-1). \tag{20}$$

By substituting Eq. (20) in Eq. (16), we obtain

$$d(i,j;\ell) = \frac{\log\left(\alpha_j(\ell)\beta_j(\ell)\right) + \log(N-1) - L\log(1-p)}{\log\left(\frac{p}{1-p}\right)}. \tag{21}$$

To speed-up the calculation of $\alpha_j(\ell)$ and $\beta_j(\ell)$ in Eq. (21), we calculate $P_{\mathrm{m}}(H_\ell = j|D^{(i)}) \propto \alpha_j(\ell)\beta_j(\ell)$ only for SNPs $\ell$ at which haplotype $i$ is a carrier of the derived allele, i.e. $D^{(i)}_\ell = 1$. This implies that the forward-backward algorithm is applied to a different set of SNPs depending on the haplotype $i$.

The calculated $P_{\mathrm{m}}(H_\ell = j|D^{(i)})$ at SNPs $\ell$ with $D^{(i)}_\ell = 1$ are exact up to a multiplicative constant $(1-p)^{L-L^{(i)}}$, where $L^{(i)} = \sum_{\ell=1}^{L} D^{(i)}_\ell$ is the number of derived mutations carried by $i$. We therefore calculate Eq. (21) as

$$d(i,j;\ell) = \frac{\log\left(\alpha^{\mathrm{d}}_j(\ell)\beta^{\mathrm{d}}_j(\ell)\right) + \log(N-1) - L^{(i)}\log(1-p)}{\log\left(\frac{p}{1-p}\right)}, \tag{22}$$

where $\alpha^{\mathrm{d}}_j(\ell)$ and $\beta^{\mathrm{d}}_j(\ell)$ are the forward and backward probabilities calculated for SNPs $\ell$ at which $D^{(i)}_\ell = 1$.

For a site $\ell$ at which haplotype $i$ is not a carrier of the derived allele, we approximate $P_{\mathrm{m}}(H_\ell = j|D^{(i)}) \propto \alpha_j(\ell)\beta_j(\ell)$ using $P_{\mathrm{m}}(H_{\ell_{\mathrm{left}}} = j|D^{(i)})$ and $P_{\mathrm{m}}(H_{\ell_{\mathrm{right}}} = j|D^{(i)})$, where $\ell_{\mathrm{left}}$, $\ell_{\mathrm{right}}$ are the nearest derived sites to either side of $\ell$, i.e., $D^{(i)}_{\ell_{\mathrm{left}}} = D^{(i)}_{\ell_{\mathrm{right}}} = 1$ and $D^{(i)}_\ell = 0$ for $\ell_{\mathrm{left}} < \ell < \ell_{\mathrm{right}}$. For such $\ell$, we approximate $P_{\mathrm{m}}(H_\ell = j|D^{(i)})$ as a weighted average of $P_{\mathrm{m}}(H_{\ell_{\mathrm{left}}} = j|D^{(i)})$ and $P_{\mathrm{m}}(H_{\ell_{\mathrm{right}}} = j|D^{(i)})$. This approximation is valid if the probability for two or more recombinations between $\ell_{\mathrm{left}}$ and $\ell_{\mathrm{right}}$ is sufficiently small. We expand

$$\begin{aligned} P_{\mathrm{m}}(H_\ell = j|D^{(i)}) = {} & P_{\mathrm{m}}(H_\ell = j,\ 0 \text{ recombinations in } [\ell_{\mathrm{left}}, \ell_{\mathrm{right}})|D^{(i)}) \\ & + P_{\mathrm{m}}(H_\ell = j,\ 1 \text{ rec. in } [\ell_{\mathrm{left}}, \ell_{\mathrm{right}})|D^{(i)}) \\ & + P_{\mathrm{m}}(H_\ell = j,\ \text{more than 1 rec. in } [\ell_{\mathrm{left}}, \ell_{\mathrm{right}})|D^{(i)}). \end{aligned} \tag{23}$$

For the third term on the right hand side of Eq. (23), we obtain

$$P_{\mathrm{m}}(H_\ell = j,\ \text{more than 1 rec. in } [\ell_{\mathrm{left}}, \ell_{\mathrm{right}})|D^{(i)}) = O((r_{\mathrm{left}} + r_{\mathrm{right}})^2). \tag{24}$$

For the second term on the right hand side of Eq. (23), we obtain

$$\begin{aligned}
&P_{\mathrm{m}}(H_\ell = j,\ 1 \text{ rec. in } [\ell_{\mathrm{left}}, \ell_{\mathrm{right}})|D^{(i)}) \\
&= P_{\mathrm{m}}(H_{\ell_{\mathrm{right}}} = j,\ 1 \text{ rec. in } [\ell_{\mathrm{left}}, \ell),\ 0 \text{ rec. in } [\ell, \ell_{\mathrm{right}})|D^{(i)}) + \\
&\quad P_{\mathrm{m}}(H_{\ell_{\mathrm{left}}} = j,\ 0 \text{ rec. in } [\ell_{\mathrm{left}}, \ell),\ 1 \text{ rec. in } [\ell, \ell_{\mathrm{right}})|D^{(i)}).
\end{aligned} \tag{25}$$

We notice that the emission probability at any site $\ell_{\mathrm{left}} < \ell < \ell_{\mathrm{right}}$ equals $1-p$ regardless of the hidden state $H_\ell$, because $D_\ell^{(i)} = 0$. Therefore, we find that the probability of a recombination (i.e., switch of hidden state), given the data $D^{(i)}$, is the same at any position between $\ell_{\mathrm{left}}$ and $\ell_{\mathrm{right}}$. We denote the recombination distance from $\ell_{\mathrm{left}}$ to $\ell$ by $r_{\mathrm{left}}$ and the recombination distance from $\ell$ to $\ell_{\mathrm{right}}$ by $r_{\mathrm{right}}$. These recombination distances correspond to the probability of a switch of hidden states between the two SNPs. We therefore obtain

$$\begin{aligned}
&P_{\mathrm{m}}(H_{\ell_{\mathrm{right}}} = j,\ 1 \text{ rec. in } [\ell_{\mathrm{left}}, \ell),\ 0 \text{ rec. in } [\ell, \ell_{\mathrm{right}})|D^{(i)}) \\
&= P_{\mathrm{m}}(\ 1 \text{ rec. in } [\ell_{\mathrm{left}}, \ell)|\ 1 \text{ rec. in } [\ell_{\mathrm{left}}, \ell_{\mathrm{right}}))P_{\mathrm{m}}(H_{\ell_{\mathrm{right}}} = j,\ 1 \text{ rec. in } [\ell_{\mathrm{left}}, \ell_{\mathrm{right}})|D^{(i)}) \\
&= \frac{r_{\mathrm{left}}}{r_{\mathrm{left}} + r_{\mathrm{right}}} P_{\mathrm{m}}(H_{\ell_{\mathrm{right}}} = j,\ 1 \text{ rec. in } [\ell_{\mathrm{left}}, \ell_{\mathrm{right}})|D^{(i)}).
\end{aligned} \tag{26}$$

Analogously, we obtain

$$\begin{aligned}
&P_{\mathrm{m}}(H_{\ell_{\mathrm{left}}} = j,\ 0 \text{ rec. in } [\ell_{\mathrm{left}}, \ell),\ 1 \text{ rec. in } [\ell, \ell_{\mathrm{right}})|D^{(i)}) \\
&= \frac{r_{\mathrm{right}}}{r_{\mathrm{left}} + r_{\mathrm{right}}} P_{\mathrm{m}}(H_{\ell_{\mathrm{left}}} = j,\ 1 \text{ rec. in } [\ell_{\mathrm{left}}, \ell_{\mathrm{right}})|D^{(i)}).
\end{aligned} \tag{27}$$

We substitute Eqs. (26) and (27) in Eq. (25) and obtain

$$\begin{aligned}
&P_{\mathrm{m}}(H_\ell = j,\ 1 \text{ rec. in } [\ell_{\mathrm{left}}, \ell_{\mathrm{right}})|D^{(i)}) \\
&= \frac{r_{\mathrm{left}}}{r_{\mathrm{left}} + r_{\mathrm{right}}} P_{\mathrm{m}}(H_{\ell_{\mathrm{right}}} = j,\ 1 \text{ rec. in } [\ell_{\mathrm{left}}, \ell_{\mathrm{right}})|D^{(i)}) \\
&\quad + \frac{r_{\mathrm{right}}}{r_{\mathrm{left}} + r_{\mathrm{right}}} P_{\mathrm{m}}(H_{\ell_{\mathrm{left}}} = j,\ 1 \text{ rec. in } [\ell_{\mathrm{left}}, \ell_{\mathrm{right}})|D^{(i)}).
\end{aligned} \tag{28}$$

For the first term on the right hand side of Eq. (23), we use that

$$\frac{r_{\mathrm{left}}}{r_{\mathrm{left}} + r_{\mathrm{right}}} + \frac{r_{\mathrm{right}}}{r_{\mathrm{left}} + r_{\mathrm{right}}} = 1, \tag{29}$$

to obtain

$$\begin{aligned}
&P_{\mathrm{m}}(H_\ell = j,\ 0 \text{ rec. in } [\ell_{\mathrm{left}}, \ell_{\mathrm{right}})|D^{(i)}) \\
&= \frac{r_{\mathrm{left}}}{r_{\mathrm{left}} + r_{\mathrm{right}}} P_{\mathrm{m}}(H_{\ell_{\mathrm{right}}} = j,\ 0 \text{ rec. in } [\ell_{\mathrm{left}}, \ell_{\mathrm{right}})|D^{(i)}) \\
&\quad + \frac{r_{\mathrm{right}}}{r_{\mathrm{left}} + r_{\mathrm{right}}} P_{\mathrm{m}}(H_{\ell_{\mathrm{left}}} = j,\ 0 \text{ rec. in } [\ell_{\mathrm{left}}, \ell_{\mathrm{right}})|D^{(i)}).
\end{aligned} \tag{30}$$

We substitute Eqs. (24), (28), and (30) in Eq. (23) and obtain

$$\begin{aligned}
P_{\mathrm{m}}(H_\ell = j|D^{(i)}) &= \frac{r_{\mathrm{left}}}{r_{\mathrm{left}} + r_{\mathrm{right}}} P_{\mathrm{m}}(H_{\ell_{\mathrm{right}}} = j|D^{(i)}) \\
&\quad + \frac{r_{\mathrm{right}}}{r_{\mathrm{left}} + r_{\mathrm{right}}} P_{\mathrm{m}}(H_{\ell_{\mathrm{left}}} = j|D^{(i)}) + O((r_{\mathrm{left}} + r_{\mathrm{right}})^2).
\end{aligned} \tag{31}$$

By multiplying both sides of Eq. (31) by $P_{\mathrm{m}}(D^{(i)})$, we obtain

$$\begin{aligned}
\alpha_j^{\mathrm{d}}(\ell)\beta_j^{\mathrm{d}}(\ell) &= \frac{r_{\mathrm{left}}}{r_{\mathrm{left}} + r_{\mathrm{right}}} \alpha_j^{\mathrm{d}}(\ell_{\mathrm{left}})\beta_j^{\mathrm{d}}(\ell_{\mathrm{left}}) \\
&\quad + \frac{r_{\mathrm{right}}}{r_{\mathrm{left}} + r_{\mathrm{right}}} \alpha_j^{\mathrm{d}}(\ell_{\mathrm{right}})\beta_j^{\mathrm{d}}(\ell_{\mathrm{right}}) + O((r_{\mathrm{left}} + r_{\mathrm{right}})^2).
\end{aligned} \tag{32}$$

By substituting Eq. (32) in Eq. (22) and dropping terms $O((r_{\text{left}}+r_{\text{right}})^2)$, we obtain an approximation for calculating $d(i, j; \ell)$.

The complexity of the modified Li-and-Stephens algorithm is reduced from $N^2L$ to $N^2L^{(i)}$. We expect $L^{(i)}$ to be independent of the sample size $N$ for large $N$. This is because in the standard coalescent, the expected TMRCA is, in the limit $N \to \infty$, given by $4N_e$, such that the expected number of derived mutations carried by haplotype $i$ since the population's TMRCA is $E[L^{(i)}] = 4N_e\mu$. In practice, this can make the application of the algorithm $> 10$ times faster even for modest samples of a few thousand individuals.

### 4 Estimating branch lengths

We have now estimated the genetic ancestry of each SNP in form of a rooted binary tree. To estimate the branch lengths $t_b$ ($b = 0, \ldots, 2N - 2$) of these trees, we use a Metropolis-Hastings type MCMC algorithm.

Before we can apply the MCMC algorithm, we notice that a recombination event changes only a few branches in adjacent trees along the genome. Some branches can persist over multiple trees. We therefore identify equivalent branches in adjacent trees along the genome (see Section 4.1). We then count the number of mutations across equivalent branches and calculate a cumulative mutation rate for each branch. This information is fed into the MCMC algorithm (see Section 4.2). We assume a coalescent prior given by the standard coalescent [8]. The effective population size is predetermined and assumed to be constant. We initialise the order of coalescence events to a random order, but such that no topological constraints are violated (see Section 4.3). We initialise the times while $k$ ancestors remain using an Expectation-maximization (EM) algorithm that calculates the maximum-likelihood estimates (MLEs) of the times while $k$ ancestors remain, where the order of coalescence events is kept fixed (see Section 4.4).

Once the constant population size model has been fitted, we can jointly infer piecewise-constant historical population sizes and branch lengths under a coalescent prior with variable historical population sizes (see Section 5).

#### 4.1 Identifying equivalent branches in neighbouring trees

Let us take a branch $b_1 = (c_1, d_1)$ from one one tree and a branch $b_2 = (c_2, d_2)$ from another tree, where $c_i$ and $d_i$ ($i = 1, 2$) denote coalescence events. We then say that branches $b_1$ and $b_2$ are equivalent if the descendants of $c_1$ coincide with those of $c_2$ and the descendants of $d_1$ coincide with those of $d_2$. To be robust to errors, we slightly relax this requirement as follows.

For every coalescence event, we store a vector of length $N$ containing its present-day descendants, where the $m$'th entry of the vector equals 1 if haplotype $m$ is below the coalescence event, and 0 otherwise. Then, for every pair of branches, with branches coming from different trees, we calculate the correlation coefficient of these vectors for the coalescence events at the lower ends of the branches and the upper ends of the branches.

Two branches are exactly equivalent if the correlation coefficient on both ends of the branches equal one. We first identify exactly equivalent pairs of branches. For the remaining branches, we require that a pair of branches is equivalent if the correlation coefficients on both ends are greater than 0.9. Notice that a branch can satisfy this condition for more than one branch in the other tree. To guarantee that each branch is associated with at most one other branch, we sort pairs of candidate branches in descending order of the correlation coefficient at the lower end of the branches. We then associate branches in the order at which they appear in this list, where we delete any entry for which one of the branches has already been associated with a different branch.

Once we identified equivalent branches, we calculate the number of mutations $m_b$ on a branch $b$ by adding the number of mutations across all equivalent branches. Next, we calculate the cumulative mutation rate for a branch. We assume that mutations occur at a constant rate of $\theta/2$ per base in coalescence time, where $\theta = 4\mu N_\mathrm{e}$ and $\mu$ is the per-generation mutation rate. For each branch $b$, we denote the mutation rate summed over bases for which $b$ persists by $\theta_b/2$.

#### 4.2 Metropolis-Hastings type MCMC to estimate branch lengths in a constant population size

We use a Metropolis-Hastings type MCMC algorithm to estimate branch lengths. The likelihood of observing branch lengths $\boldsymbol{t} = \{t_b\}_{b=0,\ldots,2N-2}$, conditional on the number of mutations on branches $\boldsymbol{m} = \{m_b\}_{b=0,\ldots,2N-2}$, is given by

$$P(\boldsymbol{t}|\boldsymbol{m}) \propto P(\boldsymbol{t})P(\boldsymbol{m}|\boldsymbol{t}) = P(\boldsymbol{t}) \prod_{b=0}^{2N-2} P(m_b|t_b), \tag{33}$$

where $P(m_b|t_b)$ is Poisson distributed with mean $\theta_b t_b/2$ and the prior $P(\boldsymbol{t})$ is given by the standard coalescent.

We can now define a reversible Markov-Chain with a unique stationary distribution which is the target distribution $P(\boldsymbol{t}|\boldsymbol{m})$. For this, we assign a label $\{1, \ldots, N-1\}$ to every coalescence event. The coalescence event that decreases the number of lineages from $k$ to $k-1$ is stored in variable $n_k$ ($k = 2, \ldots, N$). We denote the time while $k$ ancestors exist by $\tau_k$ ($k = 2, \ldots, N$). Notice that the branch lengths are uniquely determined by $\boldsymbol{n} = \{n_k\}_{k=2,\ldots,N}$ and $\boldsymbol{\tau} = \{\tau_k\}_{k=2,\ldots,N}$. In particular, the coalescent prior is given by

$$P(\boldsymbol{t}) = P(\boldsymbol{n})P(\boldsymbol{\tau}) = P(\boldsymbol{n}) \prod_{k=2}^{N} P(\tau_k), \tag{34}$$

where $\boldsymbol{n}$ is uniformly distributed over all possible orders of coalescence events and $\tau_k$ is exponentially distributed with rate $\binom{k}{2}$. We initialise $\boldsymbol{n}$ and $\boldsymbol{\tau}$ as described in Sections 4.3 and 4.4. In every step of the Metropolis-Hastings algorithm, we propose a change in $\boldsymbol{n}$ with probability $q$ and a change in $\boldsymbol{\tau}$ with probability $1-q$. In our implementation, we have chosen $q = 0.8$.

For a change in $\boldsymbol{n}$, we first choose one coalescence event $e_1 = n_k$ uniformly at random. This is the event that decreases the number of lineages from $k$ to $k-1$. We then propose to swap $e_1$ with another coalescence event $e_2 = n_h$ chosen uniformly at random from any event with age between $e_1$'s parent event and $e_1$'s daughter events. If the proposal is accepted, event $e_1$ now decreases the number of lineages from $h$ to $h-1$ and $e_2$ decreases the number of lineages from $k$ to $k-1$. We discard any proposal that would violate topological constraints of the tree if all other events retain their times. In a swap of the order of coalescence events, we keep $\boldsymbol{\tau}$ fixed. Such a swap therefore changes the lengths of six branches, but keeps all other branch lengths fixed. Denote these six branches by $b_1, \ldots, b_6$. By using Eqs. (33) and (34) and noticing that the proposal distribution is symmetric, the acceptance probability is given by

$$\min\left(1, \frac{P(\tilde{\boldsymbol{t}}|\boldsymbol{m})}{P(\boldsymbol{t}|\boldsymbol{m})}\right) = \min\left(1, \prod_{\ell=1}^{6} \frac{P(m_{b_\ell}|\tilde{t}_{b_\ell})}{P(m_{b_\ell}|t_{b_\ell})}\right), \tag{35}$$

where $\tilde{\boldsymbol{t}} = \{\tilde{t}_b\}_{b=0,\ldots.2N-2}$ are the proposed branch lengths.

For a change in $\boldsymbol{\tau}$, we choose one $\tau_k$ uniformly at random. We propose $\tilde{\tau}_k$ from an exponential distribution with expectation $\tau_k$. A change in $\tau_k$ changes the length of $k$ branches that are present at the time while $k$ ancestors remain. Denote these $k$ branches by $b_1, \ldots, b_k$. The remaining branch lengths remain unchanged. By using Eqs. (33) and (34), the acceptance probability is given by

$$\min\left(1, \frac{\tilde{\tau}_k}{\tau_k} \exp\left[-\frac{\tilde{\tau}_k}{\tau_k} + \frac{\tau_k}{\tilde{\tau}_k} - \binom{k}{2}(\tilde{\tau}_k - \tau_k)\right] \prod_{\ell=1}^{k} \frac{P(m_{b_\ell}|\tilde{t}_{b_\ell})}{P(m_{b_\ell}|t_{b_\ell})}\right). \tag{36}$$

Using this MCMC algorithm, we calculate the mean age of every coalescence event. We then calculate the branch lengths as the differences in the mean ages of coalescence events. This guarantees that the time from any tip to the root is equal, a property also known as ultrametric.

In our implementation, we initially perform $\max\{10N, 1000\}$ burn-in iterations. We then apply the algorithm until every $\tau_k$ $(k = 2, \ldots, N)$ has had at least 20 proposals and then terminate the MCMC algorithm, conditional on all branch lengths being positive, and continue until the latter condition is satisfied otherwise.

#### 4.3 Initialising the order of coalescence events

We initialise the order of coalescence events $\boldsymbol{n}$ by applying a simple MCMC algorithm. In the standard coalescent, any order of coalescence events is equally likely, provided that the order does not contradict the topological constraints of the tree. We therefore propose the following swap moves. We choose a coalescence event $n$ uniformly at random and propose a swap with another coalescence event $n'$. To ensure that we do not contradict tree topology after swapping the two events, we choose $n'$ from coalescence events that are between the parent and the children of $n$. We then assert whether $n$ is between the parent and children of $n'$. If this is the case, we accept the proposal. We accept a proposal with probability 1 because the transition probabilities are symmetric and any order of coalescence events is equally likely. We initialise the order of coalescence events by the order obtained after proposing $N^2$ swap moves.

#### 4.4 Initialising the time while $k$ ancestors remain

After initialising $\boldsymbol{n}$,we initialise the time $\tau_k$ while $k$ ancestors remain using the MLE of $\tau_k$ conditional on a fixed order of coalescence events. We denote by $\tilde{m}_b$ the number of mutations that are on branch $b$ in the interval while there are $k$ lineages remaining. We set up an EM algorithm to estimate the MLE of the parameter $\boldsymbol{\tau}$, given the data $\boldsymbol{m}$ and the unobserved variables $\tilde{\boldsymbol{m}} = \{\tilde{m}_b\}_{b=1,\ldots,2N-1}$. We denote by $\hat{\boldsymbol{\tau}}^{(s)} = \{\hat{\tau}_k^{(s)}\}_{k=N,\ldots,2}$ the estimate of the MLE of $\tau_k$ after $s$ iterations. We initialise $\hat{\tau}_k^{(0)} = \binom{k}{2}^{-1}$.

We fix $k$. Let $b_1, \ldots, b_k$ be the $k$ branches while there are $k$ lineages remaining in the tree and define $\tilde{\boldsymbol{m}}_k = \{\tilde{m}_{b_1}, \ldots, \tilde{m}_{b_k}\}$. The update rule for the EM algorithm is given by the expectation of the log likelihood function $\log P(\tau_k|\tilde{\boldsymbol{m}}_k)$, where the expectation is taken conditional on the data and parameters from the previous iteration. We obtain,

$$\hat{\tau}_k^{(s+1)} = \arg\max_{\tau_k} E\left[\log P(\tau_k|\tilde{\boldsymbol{m}}_k) \,\middle|\, \boldsymbol{m}, \hat{\boldsymbol{\tau}}^{(s)}\right]. \tag{37}$$

We first calculate $\log P(\tau_k|\tilde{\boldsymbol{m}}_k)$. By using Bayes' theorem, we obtain

$$\begin{aligned} P(\tau_k|\tilde{\boldsymbol{m}}_k) &\propto P(\tilde{\boldsymbol{m}}_k|\tau_k)P(\tau_k) \\ &= P(\tau_k)\prod_{\ell=1}^{k} P(\tilde{m}_{b_\ell}|\tau_k), \end{aligned} \tag{38}$$

where $P(\tau_k)$ is an exponential distribution with rate $\binom{k}{2}$ and $P(\tilde{m}_{b_\ell}|\tau_k)$ is a Poisson distribution with mean $\theta_{b_\ell}\tau_k/2$. By using this together with Eq.(38), we obtain

$$P(\tau_k|\tilde{\boldsymbol{m}}_k) \propto \tau_k^{\sum_{\ell=1}^{k}\tilde{m}_{b_\ell}} \exp\left[-\left(\sum_{\ell=1}^{k}\frac{\theta_{b_\ell}}{2} + \binom{k}{2}\right)\tau_k\right]. \tag{39}$$

By taking logarithms on both sides of Eq. (39), we obtain

$$\log P(\tau_k|\tilde{\boldsymbol{m}}_k) = \left(\sum_{\ell=1}^{k}\tilde{m}_{b_\ell}\right)\log(\tau_k) - \left(\sum_{\ell=1}^{k}\frac{\theta_{b_\ell}}{2} + \binom{k}{2}\right)\tau_k + \text{const.} \tag{40}$$

We substitute Eq. (40) into Eq. (37) and obtain

$$\hat{\tau}_k^{(s+1)} = \arg\max_{\tau_k}\sum_{\ell=1}^{k} E\left[\tilde{m}_{b_\ell}\,\middle|\,\boldsymbol{m}, \hat{\boldsymbol{\tau}}^{(s)}\right]\log(\tau_k) - \left(\sum_{\ell=1}^{k}\frac{\theta_{b_\ell}}{2} + \binom{k}{2}\right)\tau_k. \tag{41}$$

By taking the derivative with respect to $\tau_k$, we obtain the condition

$$\sum_{\ell=1}^{k} E\left[\tilde{m}_{b_\ell}\middle|\boldsymbol{m},\hat{\boldsymbol{\tau}}^{(s)}\right]\frac{1}{\hat{\tau}_k^{(s+1)}}-\left(\sum_{\ell=1}^{k}\frac{\theta_{b_\ell}}{2}+\binom{k}{2}\right)=0. \tag{42}$$

After reorganizing the terms, we obtain

$$\hat{\tau}_k^{(s+1)}=\frac{\sum_{\ell=1}^{k}E\left[\tilde{m}_{b_\ell}\middle|\boldsymbol{m},\hat{\boldsymbol{\tau}}^{(s)}\right]}{\sum_{\ell=1}^{k}\frac{\theta_{b_\ell}}{2}+\binom{k}{2}}. \tag{43}$$

It therefore remains to calculate $E\left[\tilde{m}_{b_\ell}|\boldsymbol{m},\hat{\boldsymbol{\tau}}^{(s)}\right]$ for $\ell=1,\ldots,k$. In the standard coalescent, mutation events are uniformly distributed on each branch. Therefore, the likelihood that an event on branch $b$ falls within the interval while $k$ lineages are remaining is given by $\hat{\tau}_k^{(s)}/t_b^{(s)}$. Therefore, the likelihood that $\tilde{m}_b$ mutations fall into that interval is given by the binomial distribution

$$P(\tilde{m}_b|m_b,\tau_k^{(s)})=\binom{m_b}{\tilde{m}_b}\left(\frac{\tau_k^{(s)}}{t_b^{(s)}}\right)^{\tilde{m}_b}\left(1-\frac{\tau_k^{(s)}}{t_b^{(s)}}\right)^{m_b-\tilde{m}_b}. \tag{44}$$

It follows that

$$E\left[\sum_{\ell=1}^{k}\tilde{m}_{b_\ell}\middle|\boldsymbol{m},\hat{\boldsymbol{\tau}}^{(s)}\right]=\sum_{\ell=1}^{k}\frac{m_{b_\ell}\tau_k^{(s)}}{t_{b_\ell}^{(s)}}. \tag{45}$$

By substituting Eq. (45) into Eq. (43), we obtain

$$\hat{\tau}_k^{(s+1)}=\frac{\sum_{\ell=1}^{k}m_{b_\ell}\frac{\tau_k^{(s)}}{t_{b_\ell}^{(s)}}}{\sum_{\ell=1}^{k}\frac{\theta_{b_\ell}}{2}+\binom{k}{2}}. \tag{46}$$

We iterate $\boldsymbol{\tau}^{(s)}$ until convergence using Eq. (46).

### 5 Estimating coalescence and mutation rates through time

We have developed a method for jointly estimating (cross-)coalescence rates and mutation rates through time, as well as branch lengths reflecting these estimated coalescence and mutation rates. This yields a self-contained method for inferring the demographic history and branch lengths, and should, for instance, improve age estimates of mutations. It can also be used to infer separation histories between diverged populations or track fine-scale population structure through time (see Section 5.1).

When estimating genealogies, we fit a constant population size model as described in Sections 4.2 to 4.4. In practice, we use $2N_\mathrm{e}=30,000$ for humans. We then use these branch lengths as the initial state for an iterative algorithm in which we repeatedly estimate coalescence rates and update branch lengths. This iterative algorithm proceeds as follows.

We first estimate a population-wide coalescence rate (see Section 5.1). Second, we estimate a population-wide mutation rate over time, where we divide time into epochs and calculate the quotient of the number of mutations that occurred in an epoch and the total branch length in that epoch. We use this estimated mutation rate in a heuristic step intended to speed-up convergence. In this step, we multiply the population-wide coalescence rate by the quotient calculated by dividing a predetermined constant mutation rate $\mu$ ($\mu=1.25\times10^{-8}$ for humans) by the estimated mutation rate over time, reflecting the assumption of a constant population-wide mutation rate through time. Finally, using this rescaled coalescence rate estimate as input, we reestimate branch lengths, now under a variable population size model (see Section 5.2). We then return to the first step.

We terminate this algorithm after five iterations. To speed up convergence and computation time, we apply this algorithm only to trees with sufficiently many mutations, which in our implementation is $N$ mutations per tree, where $N$ is the number of haplotypes. Using the output coalescence rates, we then calculate a final estimate of the branch lengths by reestimating branch lengths for all trees for a final time.

#### 5.1 Estimating the coalescence rate for a pair of haplotypes

Here, we derive an MLE for the historical coalescence rates for a pair of haplotypes, given a genealogy with estimated branch lengths. To obtain a population-wide estimate of the coalescence rate, we take the mean over all pairs of haplotypes in a population. We note that this is not the MLE for the population-wide coalescence rate assuming a panmictic population. In practice, this approach, though heuristic, allows us to avoid having to assume panmixia in this step of the algorithm. It also enables us to calculate coalescence rates for any subset of haplotypes, cross-coalescence rates between subpopulations, or track fine-scale population structure through time.

To estimate pairwise coalescence rates, we divide time into epochs. Within an epoch, we assume that the coalescence rate remains constant. For every pair of haplotypes, we estimate these piecewise constant coalescence rates using an MLE. Denote epochs by $e = 0, \dots, E$, where epoch $e$ begins at time $T_e$ and ends at time $T_{e+1}$. We denote the coalescence rate in epoch $e$ by $\gamma(e)$.

We denote the time at which the two haplotypes coalesce in tree $z$ by $t_z$. Also, we denote by $e_z$ the index of the epoch in which the haplotypes coalesce. We therefore have

$$T_{e_z} \leq t_z < T_{e_z+1}. \tag{47}$$

Conditioning on tree topology and branch lengths, the likelihood that the two haplotypes coalesce at time $t_z$ is given by

$$P(t_z) = \gamma(e_z) \exp\left[-\gamma(e_z)(t_z - T_{e_z})\right] \left[\prod_{e=1}^{e_z} \exp\left[-\gamma(e-1)(T_e - T_{e-1})\right]\right]. \tag{48}$$

By taking logarithms on both sides of Eq. (48), we obtain

$$\log P(t_z) = \log \gamma(e_z) - \gamma(e_z)(t_z - T_{e_z}) - \sum_{e=1}^{e_z} \gamma(e-1)(T_e - T_{e-1}). \tag{49}$$

Assuming independence across trees, the log-likelihood for the whole genome is given by $\sum_{z=0}^{M} \log P(t_z)$, where $M$ is the number of trees built. By differentiating with respect to $\gamma(e)$, we obtain an MLE given by

$$\hat{\gamma}(e) = \frac{n_e}{\sum_{z:e=e_z}(t_z - T_e) + \sum_{z:e<e_z}(T_e - T_{e-1})}, \tag{50}$$

where $n_e$ denotes the number of trees for which the two haplotypes coalesce in epoch $e$.

#### 5.2 Reestimating branch lengths using a coalescent prior with variable population sizes

We reestimate branch lengths using an MCMC sampling identical to that described in Section 4.2, but with a modified Eq. (36) as shown below, reflecting a coalescent prior that incorporates piecewise-constant coalescence rates. The MCMC sampler is initialised using the branch lengths of the input genealogies.

Whenever $\tau_k$ is updated in an MCMC iteration by a proposed value $\tilde{\tau}_k$, all coalescence events older than this event are updated by $\Delta\tau = \tilde{\tau}_k - \tau_k$. Therefore, older events may now coalesce in a different time epoch to before, which we need to reflect in the acceptance probability of $\tilde{\tau}_k$. We modify Eq. (36), which states the acceptance probability of a proposed update $\tilde{\tau}_k$ of $\tau_k$, to reflect a variable population size and obtain

$$\min\left(1, \frac{\tilde{\tau}_k}{\tau_k} \exp\left[-\frac{\tilde{\tau}_k}{\tau_k} + \frac{\tau_k}{\tilde{\tau}_k}\right] \prod_{\ell=1}^{k} \frac{P(m_{b_\ell}|\tilde{t}_{b_\ell})}{P(m_{b_\ell}|t_{b_\ell})} \prod_{m=2}^{k} c_m\right), \tag{51}$$

for $c_m$ ($m = 2, \ldots, k$) which we will derive below.

Let us define a function $\eta(t) \in \{0, \ldots, E\}$ mapping time $t$ to its corresponding epoch. For a piecewise constant coalescence rate as defined in Section 5.1, the time while $\tau_k$ ancestors remain, conditional on $(\tau_\ell)_{\ell=k+1,\ldots,N}$ has density

$$f_{\tau_k|\tau_{k+1},\ldots,\tau_N} = \binom{k}{2} \gamma\left(\eta\left(\sum_{\ell=k}^{N} \tau_\ell\right)\right) \exp\left[-\binom{k}{2} \int_{\sum_{\ell=k+1}^{N} \tau_\ell}^{\sum_{\ell=k}^{N} \tau_\ell} \gamma(\eta(\tau)) d\tau\right]. \tag{52}$$

It follows that the ratio of the prior probabilities of $\tilde{\tau}_k$ and $\tau_k$, conditional on $\tau_{k+1}, \ldots, \tau_N$, is given by

$$c_k = \frac{\gamma\left(\eta\left(\Delta\tau + \sum_{\ell=k}^{N} \tau_\ell\right)\right)}{\gamma\left(\eta\left(\sum_{\ell=k}^{N} \tau_\ell\right)\right)} \exp\left[-\binom{k}{2} \int_{\sum_{\ell=k}^{N} \tau_\ell}^{\Delta\tau + \sum_{\ell=k}^{N} \tau_\ell} \gamma(\eta(\tau)) d\tau\right]. \tag{53}$$

Because we also update the times of all events older than event $k$, we need to calculate the ratio of prior probabilities for these events, and we obtain for $m < k$,

$$c_m = \frac{\gamma\left(\eta\left(\Delta\tau + \sum_{\ell=m}^{N} \tau_\ell\right)\right)}{\gamma\left(\eta\left(\sum_{\ell=m}^{N} \tau_\ell\right)\right)} \exp\left[-\binom{m}{2} \int_{\sum_{\ell=m}^{N} \tau_\ell}^{\sum_{\ell=m}^{N} \tau_\ell} \left[\gamma(\eta(\tau + \Delta\tau)) - \gamma(\eta(\tau))\right] d\tau\right]. \tag{54}$$

Substituting Eqs. (53) and (54) in Eq. (52), we obtain the acceptance probability of a proposed change $\Delta\tau$ to the time while $k$ ancestors remain. In particular, we note that if $\gamma(e) \equiv 1$ for all epochs $e$, we obtain $c_k = \exp\left[-\binom{k}{2}\Delta\tau\right]$ and $c_m = 1$ ($m < k$), reducing Eq. (52) to Eq. (36).

### 6 A tree-based statistic for detecting positive selection

We define a tree-based statistic for detecting positive selection. Let $f_N$ be the number of carriers of a mutation today, and $k$ be the number of lineages when the mutation increased from frequency 1 to frequency 2. Then, under the standard coalescent model, the probability that a mutation spreads to $f_N$ haplotypes is given by (see e.g., Ref. [9])

$$P(f_N) = \frac{(f_N - 1)\binom{N - f_N - 1}{k-3}}{\binom{N-1}{k-1}}. \tag{55}$$

It follows that the probability that the mutation spreads to *at least* $f_N$ haplotypes is given by

$$p_{\mathrm{R}} = \sum_{f=f_N}^{N-k+2} P(f) = \sum_{f=f_N}^{N-k+2} \frac{(f-1)\binom{N-f-1}{k-3}}{\binom{N-1}{k-1}}. \tag{56}$$

We reject the null hypothesis that the frequency change happened under random drift (and hence no selective pressures) if this p-value is sufficiently small. We note that this test statistic does not use branch length estimates, and relies only on the order of coalescences. For this reason, we expect this test to be robust to misspecification of population size histories.

### Supplementary Note: Simulations

We test Relate on data simulated using msprime [10] which simulates the standard coalescent with recombination [11]. In the main text, we compared accuracy attained by Relate to that of ARGweaver [12]. Here, we additionally compare to accuracy of RENT+ [13] which is a recent non-parametric method that estimates genealogies more efficiently than ARGweaver. For all methods, we use default parameters and the true mutation rates, as well as effective population size. We provide the true recombination rates for ARGweaver, which are not required for RENT+.

Whenever comparisons across methods are made, we use small sample sizes ($N = 50$, $N = 200$). We note that ARGweaver is computationally infeasible for larger data sets. RENT+ is faster than ARGweaver but about 30 times slower than Relate on small data sets. We tested RENT+ on a data set of $N = 1000$ and 2.5Mb, where it ran out of memory after approximately 400 CPU hours on a computing server with 100GB of RAM.

#### 1 Accuracy of TMRCAs and mutation ages

We compare the TMRCA of pairs of haplotypes to the truth (Supplementary Fig. 3**a**). We find that estimates using Relate appear to be unbiased, whereas ARGweaver underestimates the TMRCA for recent coalescences and RENT+ exhibits a non-linear relationship to the true TMRCA. Next, we map mutations to trees to estimate mutation ages (Supplementary Fig. 3 **b**). We applied our code to do the same for ARGweaver and RENT+ because the output files of these methods do not specify on which branches the mutations occurred. We find that across all methods, age estimates of mutations appear to have a smaller deviation from the truth than TMRCAs. This may indicate that branches with at least one mutation have less uncertainty in their age estimates. In terms of accuracy, we observe similar trends as for TMRCAs, where our method appears to have the least bias in terms of dating mutations.

Additionally, we test Relate on a simulated data set with a variable population size history ($N = 200$). We simulate 200Mb using the population size inferred for GBR using Relate. In Supplementary Fig. 3 **c**, we estimate branch lengths using a constant population size of $2N_e = 30,000$. Here, we deliberately misspecified the population size history and we observe a clear bias in the estimated TMRCAs, e.g., a coalescence event dated at approximately $10^5$ years before present may have occurred between $10^4$ to $2 \times 10^6$ years before present in the true trees. In Supplementary Fig. 3**d**, we jointly infer branch lengths and population sizes for the same data set. We observe that this corrects for most of this bias, highlighting the importance of accounting for the demographic history of the sample.

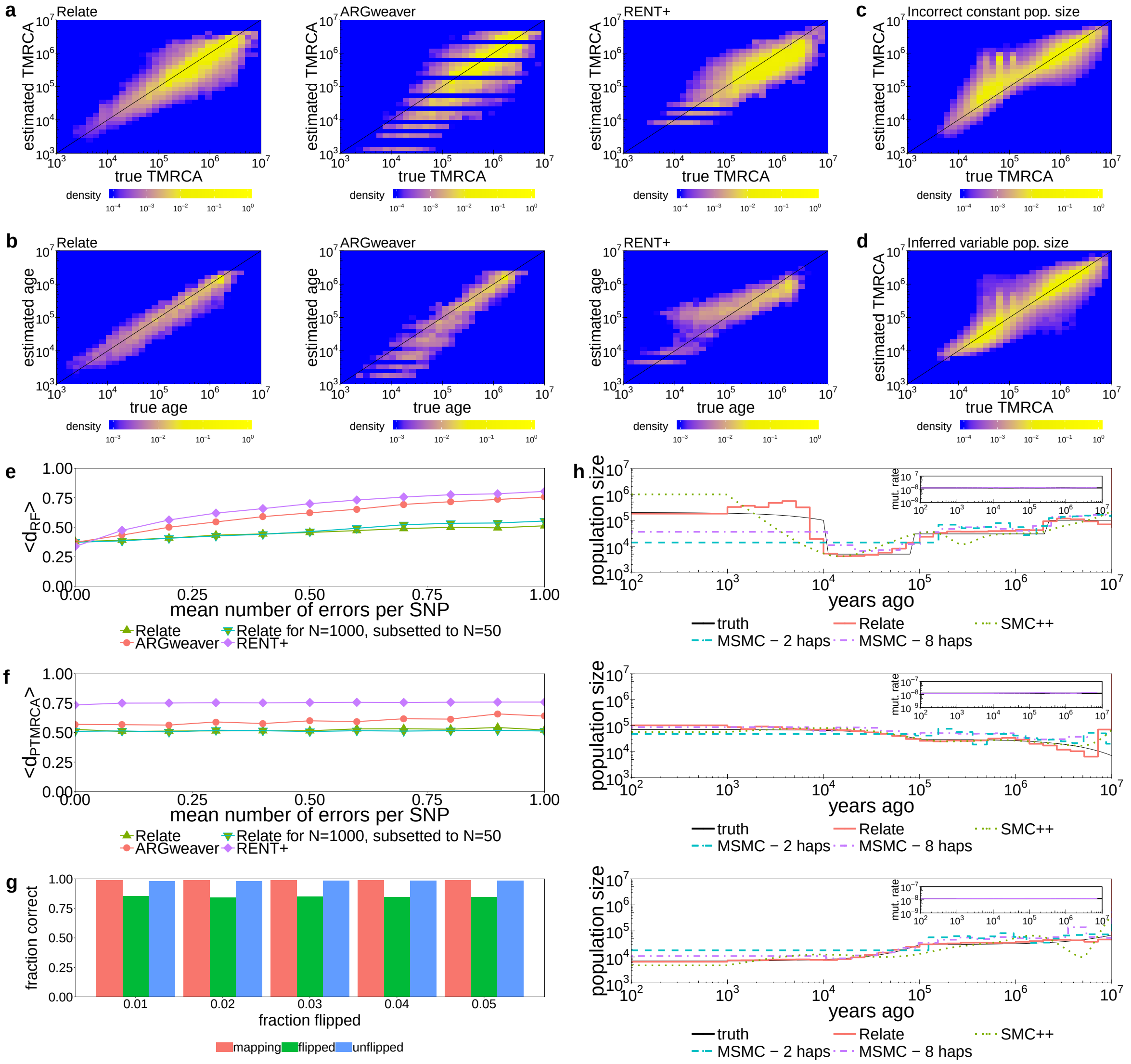

**Supplementary Figure 3: Performance of Relate on simulated data.**
**a**, Estimated times to most recent common ancestors (TMRCAs) between pairs of haplotypes compared to the truth for Relate, ARGweaver, and Rent+. **b**, Estimated ages of mutations plotted against the true age for Relate, ARGweaver, and Rent+. We determine the age of a mutation by placing it at the midpoint of the branch onto which it maps. In **a** and **b**, we simulate $N = 200$ haplotypes with $2N_e = 40,000$. **c**, TMRCAs between pairs of haplotypes compared to the truth for a simulated data set with $N = 200$ haplotypes and a population bottleneck resembling that of Europeans, where branch lengths are estimated using a constant population size of $2N_e = 30,000$. **d**, Estimated TMRCAs compared to the truth for the same example as in **c**, where branch lengths and population size history are jointly inferred. **e**, Robinson-Foulds distance and **f**, pairwise TMRCA distance averaged over 2.4Mb for Relate, ARGweaver, and RENT+. We estimate genealogies for $N = 50$ haplotypes at different number of errors. In addition, we show the accuracy of the genealogy corresponding to $N = 50$ haplotypes, embedded in an estimated genealogy for $N = 1000$ haplotypes (see Supplementary Note: Simulations, Section 2.1 for details). **g**, Robustness of Relate with respect to randomly introduced flipped mutations. We show the fraction of SNPs mapping to a unique branch, fraction of correctly flipped SNPs, and fraction of correctly unflipped SNPs for Relate. We exclude SNPs at frequency 1, which always map to the tree. We simulate 2.5Mb for $N = 200$ haplotypes with $2N_e = 30,000$. **h**, Population size estimates for simulations with a discrete bottleneck, an increasing trend, and a decreasing trend in populations size. Estimates using Relate are shown by the blue solid line. We apply SMC++ to the same data set and we also apply MSMC2 with 2 and 8 haplotypes. In the inset, we show the mutation rate over time estimated by Relate. For each scenario, we simulate 200Mb for $N = 200$ haplotypes. In all simulations, the mutation rate is set to $1.25 \times 10^{-8}$ and recombination rates are taken from the 1000 Genomes Project map for chromosome 1.

### 2 Accuracy measured using the Robinson-Foulds metric and a TMRCA metric

In addition to measuring accuracy using pairwise TMRCAs, we use two distance metrics between trees. For each distance metric, we calculate a genome-wide mean score between an estimate of the genealogy and the truth.

We compare tree topologies using the Robinson-Foulds metric $d_{\mathrm{RF}}$ adapted for rooted binary trees [14]. For each coalescence event, we find the set of present-day descendants, which we call a clade. We count the number of clades that exist in one tree but not the other. We then divide this number by $4N - 2$ such that $d_{\mathrm{RF}} = 1$ if trees are entirely different and $d_{\mathrm{RF}} = 0$ if they are exactly the same. Notice that this metric is independent of branch lengths. For two uncorrelated trees drawn from the standard coalescent, the Robinson-Foulds metric equals 1 with probability 1 as $N \to \infty$.

To also compare the accuracy of estimated branch lengths, we define a second metric $d_{\mathrm{PTMRCA}}$, in which we compare the time to the MRCA (TMRCA) of every pair of haplotypes. For each tree, we calculate a vector of lengths $\binom{N}{2}$ containing the TMRCAs between every pair of tips. We then calculate, at every SNP, the mean squared difference between the vectors corresponding to the estimated and true trees and divide the result by the diploid effective population size $N_{\mathrm{e}}$. We note that $d_{\mathrm{PTMRCA}}$ inherits its metric properties from the mean square difference together with the fact that for trees $T_1$ and $T_2$, we have $d_{\mathrm{PTMRCA}}(T_1, T_2) = 0$ if and only if $T_1 = T_2$. In this metric, the expected score for two uncorrelated trees drawn at random from the standard coalescent model equals 1.

#### 2.1 Impact of errors in the data set

We evaluate Relate at varying levels of errors introduced to the data. We simulate 2.5Mb and $N = 1000$ haplotypes with $2N_{\mathrm{e}} = 30,000$, $\mu = 1.25 \times 10^{-8}$, and recombination rates taken from the 1000 Genomes Project map for chromosome 1. We then subset this dataset to $N = 50$ haplotypes. We estimate genealogies for the same 50 haplotypes using Relate, RENT+, and ARGweaver. In addition, we estimate the genealogy for 1000 haplotypes using Relate, and extract the embedded genealogy corresponding to the same 50 haplotypes.

We introduce errors to the 50 haplotypes by first choosing a SNP and a haplotype uniformly at random and then, changing this haplotype. We find that in the absence of errors, accuracy across the three methods is similar by the $d_{\mathrm{RF}}$ metric, while Relate offers improvements for the $d_{\mathrm{PTMRCA}}$ metric. However when we introduce errors, Relate outperforms ARGweaver and RENT+ in both the Robinson-Foulds and PTMRCA metrics (Supplementary Figs. 3**e** and **f**).

#### 2.2 Impact of incorrect ancestral allele estimates

We evaluate the robustness of Relate with respect to incorrect identification of ancestral and alternative alleles. We simulate $N = 200$ haplotypes, with other parameters identical to before. We find that over 99% of SNPs can be mapped to a unique branch (Supplementary Fig. 3**g**). The fraction of correctly unflipped SNPs remains above 98% regardless of the fraction of flips introduced to the data. The fraction of correctly flipped SNPs decreases slightly but stays at around 85%. We note that for SNPs that map to a branch connected to the root of the tree, we cannot identify whether it is flipped or unflipped. We therefore excluded such SNPs from this analysis. In addition, we also excluded singleton mutations because these can always be mapped to a unique branch.

### 3 Estimating coalescence and mutation rates

We compare our algorithm for estimating historical coalescence and mutation rates to MSMC [15] and SMC++ [16] across different simulated population size histories. We simulate 200Mb for $N = 200$

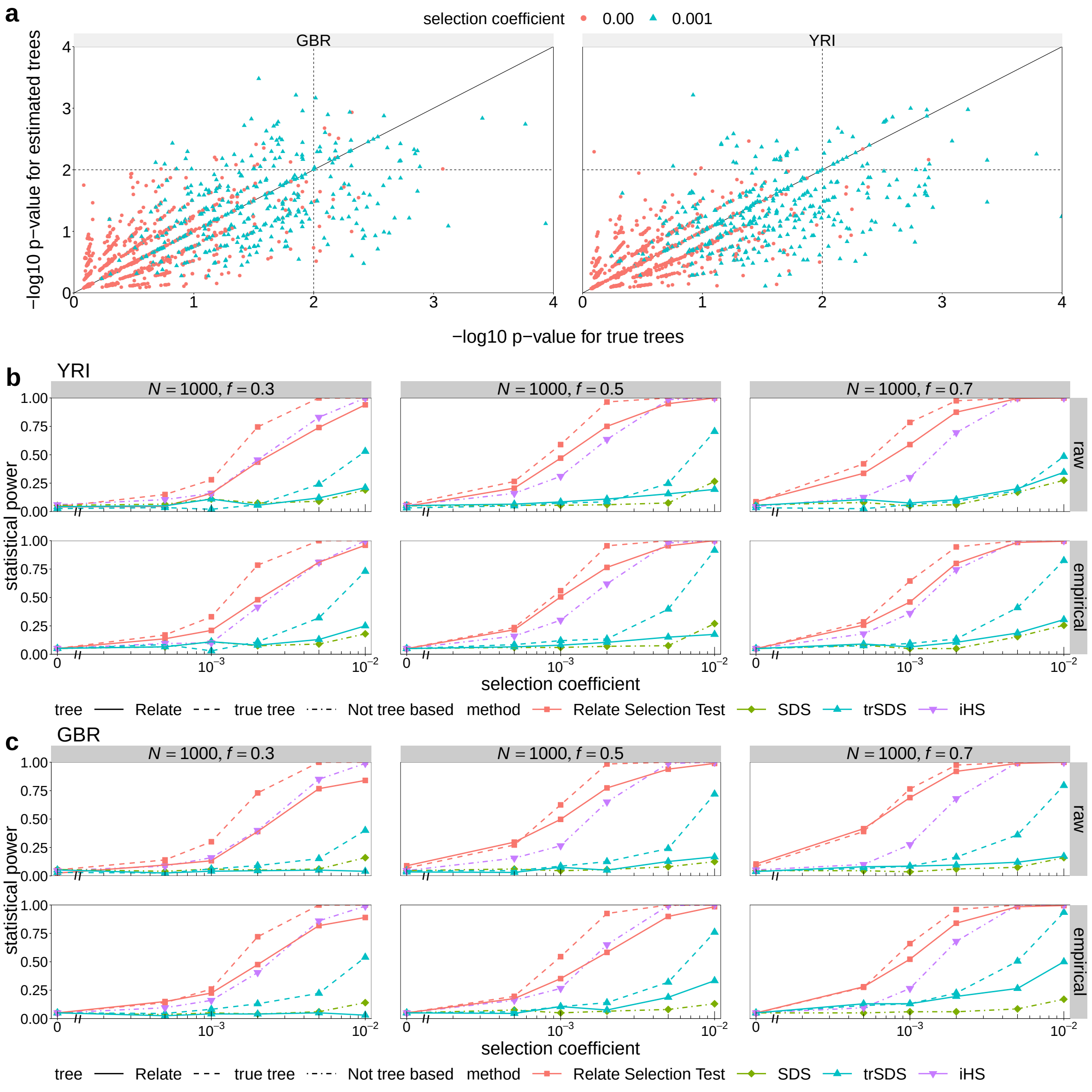

**Supplementary Figure 4: Power simulations for selection test.**
**a**, P-values for selection evidence in simulations calculated using true trees (horizontal axis) and estimated trees (vertical axis) with population size histories estimated for GBR and YRI and a present-day derived allele frequency of 0.5. We plot p-values for loci under no selection (circles) and loci under weak selection (triangles). **b**, **c**, Power simulations with $N = 1000$ haplotypes and present-day derived allele frequencies of 0.3, 0.5, and 0.7. We assume a population size history estimated for YRI (**c**), and GBR (**c**), respectively. The significance threshold is 0.05. We show power estimates using the p-values for trees estimated by Relate, as well as those for the true trees. In both cases, we estimate power using raw p-values of our test statistic (top row) and empirical p-values given the distribution of raw p-values in the neutral case (bottom row). For iHS, SDS, and trSDS, power is estimated by standardising raw scores by the frequency specific mean and standard deviation under the null. In the top row, we assume a standard normal distribution of the standardised score and in the bottom row, we calculate empirical p-values by determining a critical score corresponding to the 0.05 significance level in the neutral case.

haplotypes and a constant mutation rate of $1.25 \times 10^{-8}$. The recombination rates are taken from the 1000 Genomes Project map for chromosome 1.

The command line used for MSMC is

```
msmc_2.0.0 \
     -t 6 \
     -p 1*2+15*1+1*2 \
     -o msprime.msmc2 \
     msprime.multihetsep.txt
```

where "msprime.multihetsep.txt" denotes the input filename and "msprime.msmc2" denotes the output filename. The command line used for SMC++ is

```
smc++ estimate \
     --regularization_penalty 5.0 \
     --knots 16 \
     --timepoints 35,100000 \
     1.25e-8 \
     -o analysis/ \
     out/msprime.smc.gz
```

where "analysis/" denotes the directory into which the output of SMC++ is saved and "out/msprime.smc.gz" denotes the input filename. We noticed that the accuracy of SMC++ is sensitive to the choice of parameters "regularization_penalty" and "knots". Our choice of these parameters is based on advice (personal communication) from the authors of Ref. [16].

We simulate three scenarios: a discrete bottleneck, with a ten-fold change in population size ranging from $2N_{\mathrm{e}} = 7,000$ to $2N_{\mathrm{e}} = 70,000$, an increasing trend, and a decreasing trend (Supplementary Fig. 3**h**). We find that Relate estimates the population size with high accuracy in all three scenarios. The accuracy of MSMC depends on the number of haplotypes used, where its population size estimate is accurate up until 100,000 years before present with 2 haplotypes and 10,000 with 8 haplotypes. SMC++ has a comparable accuracy to MSMC, but detects trends in the more recent past as well.

A consequence of correctly estimated population sizes is a mutation rate that remains constant and close to $\mu = 1.25 \times 10^{-8}$ through time. The insets of Supplementary Fig. 3**h** show that indeed the mutation rate stays mostly constant and close to the truth.

### 4 Positive natural selection

In Supplementary Fig. 4**a**, we plot p-values calculated with the true trees against p-values calculated using estimated trees for selection coefficients equalling 0.00 and 0.001. We observe a high correlation and a clear shift in p-values when selection is turned on.

In Supplementary Fig. 4**b** and **c**, we estimate the power of our selection test for varying present-day derived allele frequencies. We compare our method to the integrated haplotype score (iHS) [17] and singleton density score (SDS) [18]. We use selscan [19] and hapbin [20] to calculate iHS scores. In addition, we define a tree-based SDS score (trSDS) as proposed in Ref. [21]. For iHS, SDS, and trSDS, we standardise the raw scores using the frequency specific empirical mean and standard deviation for the neutral case, which is an idealised setting that should favour power estimates of both methods. We find that the statistical power of our test outperforms iHS, SDS, and trSDS when using the true trees in all considered scenarios. Using trees estimated by Relate, we attain a higher power than iHS for weak selection and a similar or slightly lower power than iHS for strong selection. While iHS is similarly powered for all present-day derived allele frequencies, the power of our test statistic increases with larger derived allele frequency. Both methods outperform SDS and trSDS, which have been designed to perform better for very recent selection on standing variation. In most cases, trSDS outperforms SDS. This is an encouraging result, suggesting that adapting existing ideas for trees can improve power. We postpone a more thorough study in this direction, including simulating scenarios that are more suited to SDS and trSDS, to future work.

### Supplementary Note: 1000 Genomes Project data set

#### 1 Runtime

Relate terminated after less than 2 CPU years, or 4 days when run on a high-performance cluster, using up to 300 cores (see Supplementary Table 1). Each core had a maximum memory allowance of 16GB and was equipped with an Intel Ivybridge 2.4 GHz or Intel Haswell 2.6 GHz processor.

#### 2 Number of trees built

As a first indication of good accuracy of the inferred genealogy, we find a high correlation between the number of trees built and the recombination distance in bins of $10^5$ SNPs (Supplementary Fig. 5**a**). In a low recombination rate region, we estimate that we construct approximately one tree every 125 SNPs. The number of trees we build is slightly inflated by the fact that in our implementation of Relate, we rebuild a tree after 200 - 1000 SNPs for computational reasons. Consequently, not every new tree represents a recombination event. We also find that the mean number of SNPs mapping onto a tree increases with reducing local recombination distance (Supplementary Fig. 5**b**).

#### 3 CpG mutations map less frequently than other mutations

We next investigate SNPs that could not be mapped to a unique branch. Singleton mutations always map to a unique branch, such that we exclude singletons from the following analysis. For any non-singleton, 14.3 % of SNPs do not map to a unique branch (Supplementary Fig. 5**c**). SNPs with a small derived allele frequency are particularly susceptible to errors in the data. How the derived allele frequency of a mutation affects the fraction of non-mapping SNPs is shown in Supplementary Fig. 5**f**. We can see that the fraction of non-mapping SNPs is relatively high for rare SNPs and rapidly decreases to around 5% for more frequent SNPs. If we exclude any SNP with a derived allele frequency of less or equal to 10, only 4 % of SNPs do not map to a unique branch (Supplementary Fig. 5**c**).

We further notice that CpG sites, which are sites at which a C nucleotide is followed by a G nucleotide, are known to have a significantly higher rate of mutations $CG \to TG$ [22]. We should therefore expect a higher probability of observing two or more mutations at the same genomic position. Such SNPs usually cannot be mapped to a unique branch. Indeed, we observe that the fraction of non-mapping CpG transitions is 24.3 % which is two-fold higher than the overall average.

To further study the effect of adjacent nucleotides, we group mutations by their two neighbouring nucleotides in sequence. This yields 96 categories after accounting for equivalent mutations due to symmetry on the complementary strands. We observe that the fraction of non-mapping SNPs is highly variable with respect to the mutation category (Supplementary Fig. 5**e**). In some categories, we observe a higher fraction of non-mapping SNPs resembling those of CpG transitions. We have identified that the spike in TTA $\to$ TAA is likely caused by alignment errors, where SNPs are mistakenly identified

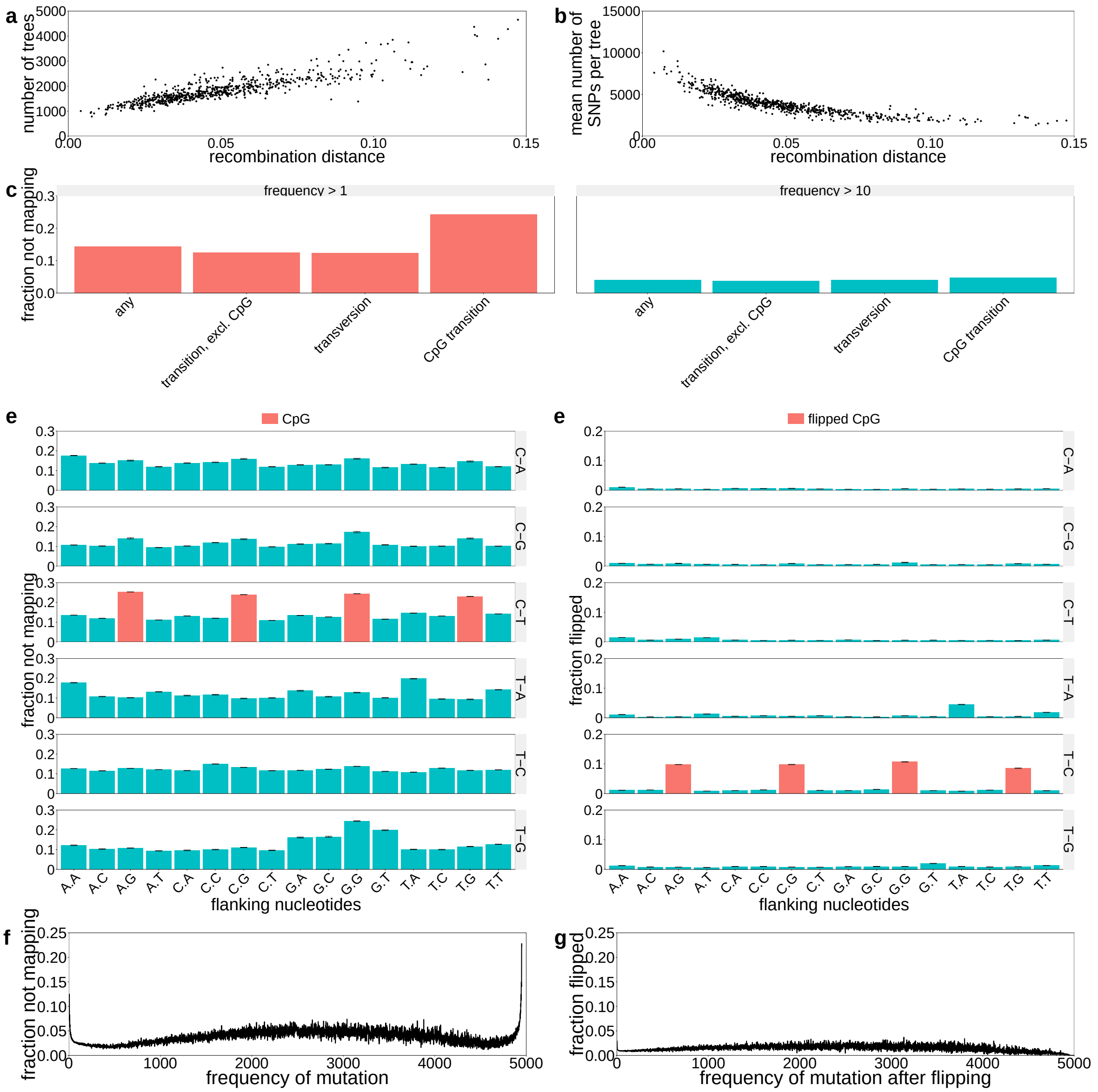

**Supplementary Figure 5: Properties of the genealogy constructed for the 1000 Genomes Project data set.**

**a**, Number of trees built versus the recombination distance for all 22 chromosomes. **b**, Mean number of SNPs that map to a unique branch versus the recombination distance in that bin. Every point represents a subregion of $10^5$ SNPs. **c**, Fraction of SNPs that could not be mapped to a unique branch for SNPs excluding singletons (left) and SNPs with derived allele frequencies larger than 10 (right). **d**, Fraction of SNPs that could not be mapped to a unique branch for all 96 possible triplet mutations, excluding singletons. **e**, Fraction of SNPs that were flipped for all 96 possible triplet mutations, excluding singletons. In **d** and **e**, CpG transitions are indicated in red. The 95% condidence intervals are indicated by black brackets. **f**, Fraction of non-mapping SNPs by derived allele frequency of the mutation in the sample. For each frequency, we divide the number of non-mapping mutations of that frequency by the number of mutations of that frequency. **g**, Fraction of flipped SNPs by derived allele frequency of the mutation after flipping. For each frequency, we divide the number of flipped SNPs of that frequency (after flipping) by the number of SNPs of that frequency.

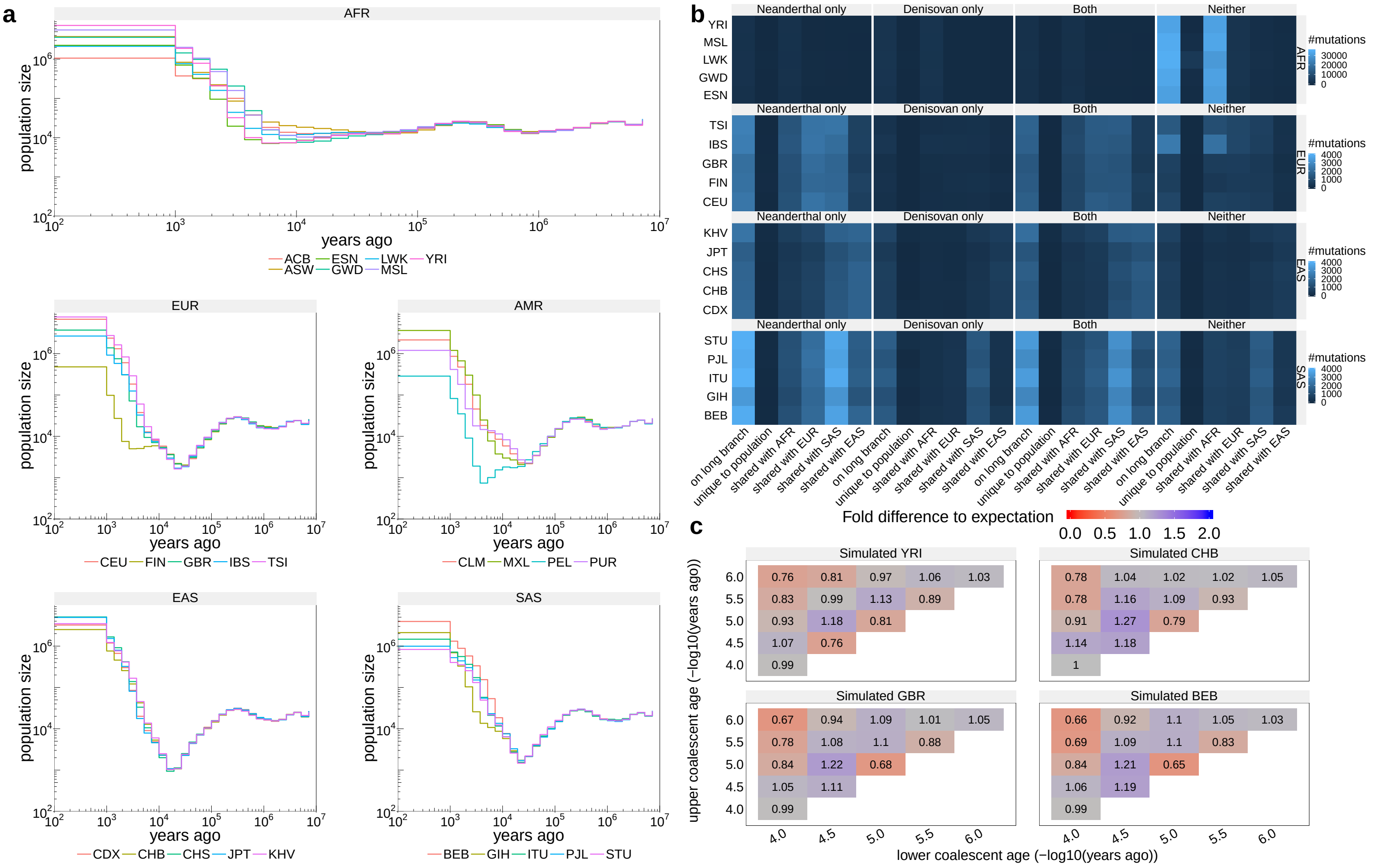

**Supplementary Figure 6: Historical effective population sizes and evidence of introgression.**
**a**, Historical effective population sizes for all 26 populations in the 1000 Genomes Project dataset. For each population, we first extract the genealogy corresponding to that population. We then estimate the population size using this genealogy. **b**, Number of mutations on branches with an upper end older than 1M YBP and lower end younger than 30,000 YBP, categorised by whether the mutation is additionally found only in Neanderthals, only in Denisovans, both, or neither. For each category, we also distinguish whether the mutation is unique to the population of interest or shared with other populations in AFR, EUR, SAS, or EAS. **c**, Number of mutations binned by age of upper and lower coalescence event, relative to the expected number of mutations when randomising topology while fixing ages of coalescence events for four simulated data sets (**Methods**). We simulated $3 \times 10^9$ bases with population size histories of YRI, CHB, GBR, and BEB.

in regions of repeated T alleles followed by repeated A alleles. Currently, we cannot explain spikes in other categories, such as ATA → AAA, or GT → GG.

Mutations that occur at the same genomic position both in humans and other primates can cause confusion of the ancestral and derived alleles which we can detect as flipped SNPs. The fraction of flipped SNPs appears to be less dependent on the mutation category (Supplementary Fig. 5**e**). However, we again find a clear signal for CpG transitions $CG \rightarrow TG$. This is likely to be caused by a mutation at the same genomic position in humans and other primates, leading to an error in estimating the ancestral allele. For other categories, the fraction of flipped SNPs is approximately 1%. The fraction of flipped SNPs is moderately dependent on the frequency of the alternative allele in the sample (Supplementary Fig. 5**g**).

### 4 Historical population sizes of 26 populations

We estimate historical population sizes for all 26 populations in the 1000 Genomes Project data set (Supplementary Fig. 6). All non-African groups show a severe bottleneck following their out-of-Africa migration. As already discussed in the main text, we observe a second bottleneck in the Finnish (FIN) population around 3,000 to 9,000 YBP. Indications of a second bottleneck can also be observed in Gujarati individuals (GIH) and a severe second bottleneck can be observed in the Peruvian population (PEL).

### 5 Positive selection

#### 5.1 Genome-wide significant hits for positive selection

Because there is no guarantee that the mutation with the lowest p-value is the causal variant, we determine genomic regions that should contain the causal variant. In Supplementary Table 3, we list all regions in the genome that contain genome-wide significant hits ($p < 5 \times 10^{-8}$) in at least three populations.

To determine genomic regions under selection, we cluster genome-wide significant hits into blocks, such that any two SNPs in different blocks have a Pearson correlation coefficient $r^2 < 0.5$, and for any SNP, there is always another SNP in the same block with $r^2 \geq 0.5$. We then extend these identified genomic regions, by including any SNP within 2Mb that have an $r^2 \geq 0.5$ to any of the genome-wide significant hits in a block.

We annotate each region using genes, eQTLs, GWAS hits, and non-synonymous substitutions at SNPs with highest $r^2$ to the listed mutation. We combined the genome-wide significant GWAS hits ($p < 5 \times 10^{-8}$) of the GWAS catalogue [23] and the UK Biobank project [24]. We used the eQTL annotation provided by GTEx [25] (see URLs).

In Supplementary Table 3, we list all identified regions, where we choose the SNP with the lowest p-value in that genomic region for each population. We observe that all considered geographic regions (AFR, EUR, SAS, EAS) are represented in this table. With the exception of 6 identified genomic regions, populations come from same geographic region in each identified genomic region. To the best of our knowledge, 13 out of 35 regions have been reported previously. We additionally record the statistic that attains a value in the 0.05% tail of its empirical distribution in the 1000 Genomes Project dataset according to the PopHumanScan resource [26] (URLs), whenever the region reported in PopHumanScan overlaps the region listed in this table and is attributed to a population listed in this table. Including such regions, our method detects 18 previously reported regions. Of all unreported regions, 12 are attributed only to African populations, and 5 unreported regions are attributed to non-Africans.

In 9 regions, we find an eQTL and in 8 regions, we find a GWAS hit within $r^2 \geq 0.8$ of the SNP with the lowest p-value for selection evidence. As discussed in the main text, two well known regions of positive selection, EDAR and LCT, each fall into one identified genomic regions. While the causal

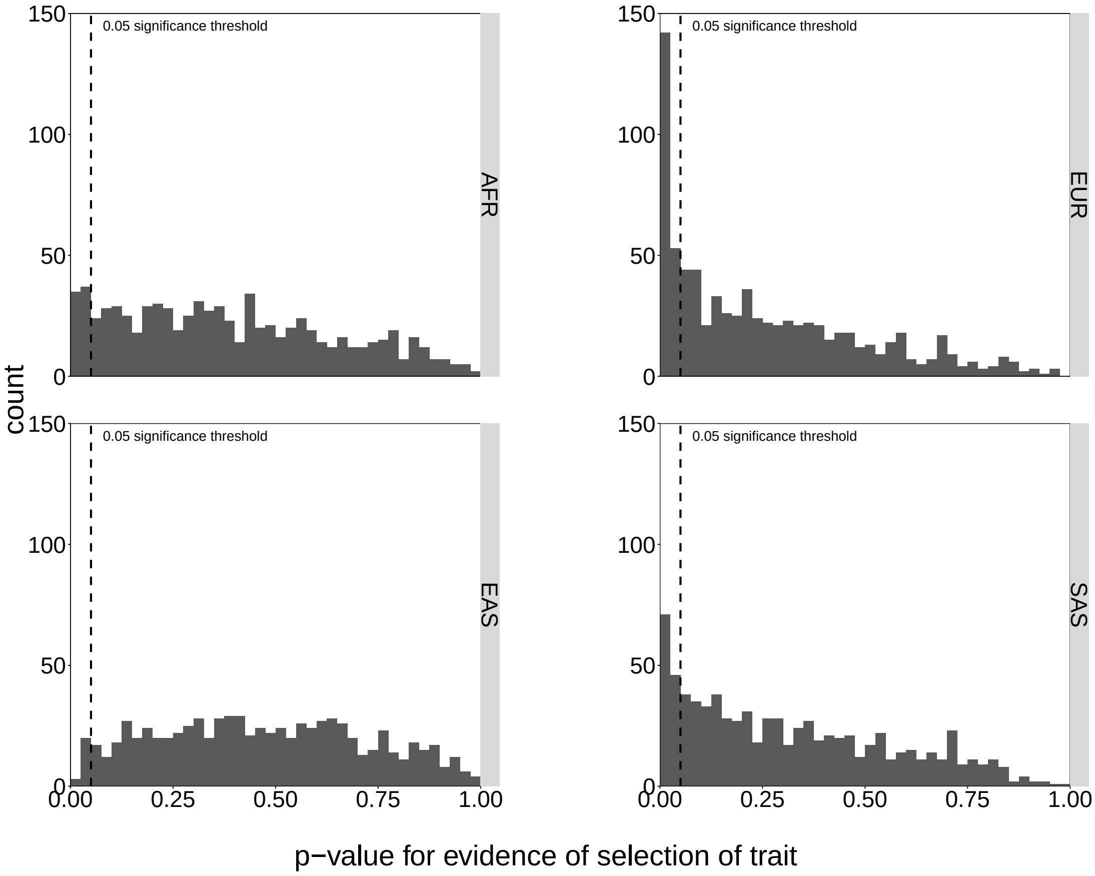

**Supplementary Figure 7: Histograms of p-values for evidence of selection on complex traits.** We aggregated both effect directions of 84 considered traits, as well as populations in each of the four considered geographic regions (AFR, EAS, EUR, SAS).

variants are not those with the lowerst p-value for selection evidence, they are in $r^2 \geq 0.8$ in both cases. Moreover, we identify a previously unreported hit in the EDARADD gene, which is known to directly interact with the EDAR protein [27], to be selected in all Southern Asian populations, as well as the Finnish population. This mutation achieves a selection p-value of less than $10^{-6}$ in all European populations.

**Supplementary Table 1: Runtime of Relate on the 1000 Genomes Project data set.** CPU time spent in days (second column) and maximum memory usage in Mb (third column) in each stage of the algorithm applied to 4956 haplotypes of the 1000 Genomes Project dataset.

| | CPU days | Max. memory usage (Mb) |
|---|---|---|
| Modified Li-and-Stephens algorithm | 6.9 | 2,802 |
| Tree building | 132.2 | 14,632 |
| Finding Equivalent branches | 1.0 | 7,435 |
| Estimating branch lengths | 482.2 | 83 |
| Other | 0.2 | 6310 |
| Total | 622.4 | |

**Supplementary Table 2: Number of 1000 Genomes Project samples used in our analysis by population label.**

| | | | | | | | | | | | | |
|---|---|---|---|---|---|---|---|---|---|---|---|---|
| ACB | ASW | BEB | CDX | CEU | CHB | CHS | CLM | ESN | FIN | GBR | GIH | GWD |
| 95 | 60 | 85 | 92 | 98 | 102 | 104 | 93 | 98 | 98 | 90 | 102 | 112 |
| IBS | ITU | JPT | KHV | LWK | MSL | MXL | PEL | PJL | PUR | STU | TSI | YRI |
| 106 | 101 | 103 | 98 | 98 | 84 | 63 | 84 | 95 | 103 | 101 | 106 | 107 |

| AMR | (Americas) |
|---|---|
| CLM | Colombian in Medellin, Colombia |
| MXL | Mexican Ancestry in Los Angeles, CA, USA |
| PEL | Peruvian in Lima, Peru |
| PUR | Puerto Rican in Puerto Rico |

| SAS | (Southern Asians) |
|---|---|
| BEB | Bengali in Bangladesh |
| GIH | Gujarati Indian in Houston, TX, USA |
| ITU | Indian Telugu in the UK |
| PJL | Punjabi in Lahore, Pakistan |
| STU | Sri Lankan Tamil in the UK |

| EAS | (East Asians) |
|---|---|
| CDX | Chinese Dai in Xishuangbanna, China |
| CHB | Han Chinese in Bejing, China |
| CHS | Southern Han Chinese, China |
| JPT | Japanese in Tokyo, Japan |
| KHV | Kinh in Ho Chi Minh City, Vietnam |

| EUR | (Europeans) |
|---|---|
| CEU | Utah residents with Northern and Western European ancestry |
| IBS | Iberian populations in Spain |
| FIN | Finnish in Finland |
| GBR | British in England and Scotland |
| TSI | Toscani in Italy |

| AFR | (Africans) |
|---|---|
| ACB | African Caribbean in Barbados |
| ASW | African Ancestry in Southwest US |
| ESN | Esan in Nigeria |
| GWD | Gambian in Western Division, The Gambia |
| LWK | Luhya in Webuye, Kenya |
| MSL | Mende in Sierra Leone |
| YRI | Yoruba in Ibadan, Nigeria |

**Supplementary Table 3: Genome-wide significant hits for positive selection.**
Regions containing a SNP with p-value for selection evidence of less than $5 \times 10^{-8}$ in at least three populations. We list genes, eQTLs, and GWAS at mutations with $r^2 \geq 0.5$, where a bold font correposonds to $r^2 = 1.0$, a plain font corresponds to $r^2 \geq 0.8$, and brackets correspond to $r^2 \geq 0.5$. If a non-synonymous mutation falls within $r^2 \geq 0.5$, we indicate this similarly in the NSM column. BP denotes the base-pair position of the SNP (GRCh37). In the notes column, we indicate whether this region has been highlighted in a previous study. Additionally, we record the statistic that attains a significant value according to the PopHumanScan resource, whenever the region reported in PopHumanScan overlaps the region listed in this table and is attributed to a population listed in this table. See Supplementary Note: 1000 Genomes Project data set, Section 5.1 for details

**GWAS catalogue phenotypes**:

a Helix rolling

b Cholesterol, total+;Blood metabolite levels-

c Nonsyndromic cleft lip with cleft palate

d Nonsyndromic cleft lip with cleft palate

e Glaucoma (primary open-angle)

**UK BIOBANK phenotypes**: **BIOBANK phenotypes**:

1. Forced vital capacity (FVC)-;Monocyte percentage+;Place of birth in UK - east co-ordinate-;Place of birth in UK - north co-ordinate+

2. Arm fat mass (left)+;Arm fat mass (right)+;Body fat percentage+;Forced vital capacity (FVC)-;Leg fat mass (left)+;Leg fat mass (right)+;Monocyte percentage+;Place of birth in UK - east co-ordinate-;Place of birth in UK - north co-ordinate+;Trunk fat mass+;Trunk fat percentage+;Whole body fat mass+

3. General happiness with own health+;High light scatter reticulocyte count-;High light scatter reticulocyte percentage-;Immature reticulocyte fraction-;Impedance of arm (right)+;Impedance of leg (left)+;Impedance of leg (right)+;Impedance of whole body+;Reticulocyte count-;Reticulocyte percentage-;Standing height+

4. Standing height-

5. White blood cell (leukocyte) count-

6. Impedance of leg (right)-;Impedance of whole body-;Red blood cell (erythrocyte) distribution width+

7. Leg fat-free mass (right)+;Leg predicted mass (right)+;Red blood cell (erythrocyte) distribution width+;Trunk fat-free mass+;Trunk predicted mass+;Whole body fat-free mass+;Whole body water mass+

8. Birth weight-;High light scatter reticulocyte count+;High light scatter reticulocyte percentage+;Nervous feelings+;Reticulocyte count+;Reticulocyte percentage+;Systolic blood pressure, automated reading+

9. Hair colour (natural, before greying): Blonde-;Hair colour (natural, before greying): Dark brown+

10. Hair colour (natural, before greying): Blonde-;Hair colour (natural, before greying): Dark brown+

11. 3mm strong meridian (left)+;3mm weak meridian (left)+;6mm strong meridian (left)+;6mm weak meridian (left)+;Age high blood pressure diagnosed-;Arm fat-free mass (left)-;Arm fat-free mass (right)-;Arm predicted mass (left)-;Arm predicted mass (right)-;Basal metabolic rate-;Basophill count+;Birth weight of first child-;Birth weight-;Blood clot, DVT, bronchitis, emphysema, asthma, rhinitis, eczema, allergy diagnosed by doctor: Hayfever, allergic rhinitis or eczema-;Blood clot, DVT, bronchitis, emphysema, asthma, rhinitis, eczema, allergy diagnosed by doctor: None of the above+;Comparative height size at age 10-;Coronary atherosclerosis+;Diagnoses - main ICD10: I21 Acute myocardial infarction+;Diastolic blood pressure, automated reading+;Diseases of the circulatory system+;Duration to first press of snap-button in each round+;Eosinophill count+;Eosinophill percentage+;Ever smoked+;Haematocrit percentage+;Haemoglobin concentration+;High light scatter reticulocyte count+;High light scatter reticulocyte percentage+;Hip circumference-;Illnesses of father: Heart disease+;Illnesses of siblings: High blood pressure+;Illnesses of siblings: None of the above (group 1)-;Immature reticulocyte fraction+;Impedance of arm (left)+;Impedance of arm (right)+;Impedance of leg (right)+;Impedance of whole body+;Ischaemic heart disease, wide definition+;Leg fat-free mass (left)-;Leg fat-free mass (right)-;Leg predicted mass (left)-;Leg predicted mass (right)-;Long-standing illness, disability or infirmity+;Lymphocyte count+;Lymphocyte percentage+;Major coronary heart disease event excluding revascularizations+;Major coronary heart disease event+;Mean corpuscular haemoglobin+;Mean corpuscular volume+;Mean sphered cell volume-;Mean time to correctly identify matches+;Medication for cholesterol, blood pressure, diabetes, or take exogenous hormones: Blood pressure medication+;Medication for cholesterol, blood pressure, diabetes, or take exogenous hormones: None of the above-;Medication for cholesterol, blood pressure or diabetes: Blood pressure medication+;Medication for cholesterol, blood pressure or diabetes: None of the above-;Monocyte count+;Myocardial infarction+;Myocardial infarction, strict+;Neutrophill count+;Neutrophill percentage-;Non-cancer illness code, self-reported: hypertension+;Non-cancer illness code, self-reported: psoriasis+;Number of self-reported non-cancer illnesses+;Past tobacco smoking-;Platelet count+;Platelet crit+;Platelet distribution width+;Red blood cell (erythrocyte) count+;Reticulocyte count+;Reticulocyte percentage+;Smoking status: Never-;Smoking status: Previous+;Standing height-;Systolic blood pressure, automated reading+;Taking other prescription medications+;Trunk fat-free mass-;Trunk predicted mass-;Weight-;White blood cell (leukocyte) count+;Whole body fat-free mass-;Whole body water mass-

12. Comparative body size at age 10-;Impedance of leg (left)+;Impedance of leg (right)+;Lymphocyte percentage-;Monocyte count+;Monocyte percentage+;Neutrophill count+;White blood cell (leukocyte) count+

13. Arm fat mass (left)-;Arm fat mass (right)-;Arm fat percentage (right)-;Basal metabolic rate-;Body mass index (BMI)-;Comparative body size at age 10-;Duration to first press of snap-button in each round+;Eosinophill percentage+;Haematocrit percentage-;Haemoglobin concentration-;High light scatter reticulocyte count-;High light scatter reticulocyte percentage-;Impedance of leg (left)+;Leg fat-free mass (left)-;Leg fat-free mass (right)-;Leg fat mass (left)-;Leg fat mass (right)-;Leg predicted mass (left)-;Leg predicted mass (right)-;Lymphocyte percentage+;Mean corpuscular haemoglobin+;Mean corpuscular volume+;Mean platelet (thrombocyte) volume+;Mean reticulocyte volume+;Mean sphered cell volume+;Mean time to correctly identify matches+;Nap during day+;Neutrophill count-;Neutrophill percentage-;Platelet count-;Red blood cell (erythrocyte) count-;Red blood cell (erythrocyte) distribution width-;Reticulocyte count-;Waist circumference-;Weight-;Whole body fat mass-

14. Arm fat mass (right)-;Arm fat percentage (right)-;Body mass index (BMI)-;Comparative body size at age 10-;Duration to first press of snap-button in each round+;Eosinophill percentage+;Haematocrit percentage-;Haemoglobin concentration-;High light scatter reticulocyte count-;High light scatter reticulocyte percentage-;Lymphocyte percentage+;Mean corpuscular haemoglobin+;Mean corpuscular volume+;Mean platelet (thrombocyte) volume+;Mean reticulocyte volume+;Mean sphered cell volume+;Mean time to correctly identify matches+;Nap during day+;Neutrophill count-;Neutrophill percentage-;Platelet count-;Red blood cell (erythrocyte) count-;Red blood cell (erythrocyte) distribution width-;Reticulocyte count-;Reticulocyte percentage-;Sitting height+;Waist circumference-;White blood cell (leukocyte) count-

15. Mean platelet (thrombocyte) volume-;Platelet count+;Platelet distribution width-

| CHR:REGION | ID/BP | population | logP | gene by r2 | eQTL | GWAS | NSM | AFR | EUR | SAS | EAS | notes |
|---|---|---|---|---|---|---|---|---|---|---|---|---|
| 1:76.1-76.4 | BP76111796 | KHV (EAS)<br>ITU (SAS)<br>STU (SAS) | $6.8 \times 10^{-10}$<br>$1.2 \times 10^{-8}$<br>$3.5 \times 10^{-9}$ | SLC44A5 | (MSH4) | | | 0.03 | 0.21 | 0.39 | 0.63 | [17]; Fay & Wu's H |
| 1:236.5-236.6 | rs76420343 | FIN (EUR)<br>BEB (SAS)<br>GIH (SAS)<br>ITU (SAS)<br>PJL (SAS) | $1 \times 10^{-9}$<br>$1.4 \times 10^{-8}$<br>$2.6 \times 10^{-9}$<br>$4.2 \times 10^{-8}$<br>$4.8 \times 10^{-8}$ | **EDARADD** | EDARADD | | | 0.08 | 0.52 | 0.38 | 0.22 | The EDARADD protein is known to directly interact with the EDAR protein [27]. |
| | rs79881482 | STU (SAS) | $3 \times 10^{-9}$ | **EDARADD** | EDARADD | | | 0.06 | 0.52 | 0.38 | 0.22 | |
| 2:108.9-109.6 | rs11123695 | CDX (EAS)<br>CHB (EAS)<br>CHS (EAS)<br>KHV (EAS) | $7.4 \times 10^{-10}$<br>$1.2 \times 10^{-12}$<br>$5.7 \times 10^{-12}$<br>$8 \times 10^{-12}$ | **GCC2** | (GCC2) | a | EDAR | 0 | 0.01 | 0.01 | 0.85 | [17, 28]; iHS, XPEHH |
| 2:135.6-136.9 | rs6730157 | FIN (EUR) | $1.4 \times 10^{-8}$ | **RAB3GAP1** | **MCM6** | **1**,b | (RAB3GAP1) | 0 | 0.49 | 0.12 | 0 | [28, 17, 18, 29, 30, 31] |
| | rs1375131 | GBR (EUR) | $4.3 \times 10^{-10}$ | **ZRANB3** | **MCM6** | **1**,b | (RAB3GAP1) | 0 | 0.49 | 0.12 | 0 | iHS, XPEHH |
| | rs56369224 | CEU (EUR) | $3.3 \times 10^{-9}$ | **R3HDM1** | MCM6 | **2**,b | (RAB3GAP1) | 0 | 0.49 | 0.12 | 0 | Fu & Li's D, $\alpha$ |
| 2:168.4-168.5 | rs150960584 | CEU (EUR)<br>GBR (EUR)<br>IBS (EUR)<br>TSI (EUR) | $1.7 \times 10^{-9}$<br>$2.8 \times 10^{-8}$<br>$7.3 \times 10^{-9}$<br>$2.5 \times 10^{-9}$ | **U7** | | | | 0.02 | 0.67 | 0.2 | 0.04 | – |
| 3:48.7-50.5 | rs139083518 | KHV (EAS) | $3.6 \times 10^{-9}$ | **DAG1** | (NAT6) | | | 0 | 0.01 | 0.02 | 0.6 | [31, 17] |
| | rs201632611 | CDX (EAS)<br>CHS (EAS) | $3 \times 10^{-8}$<br>$1.3 \times 10^{-11}$ | **GNAI2** | NAT6 | (3) | (C3orf45) | 0 | 0.01 | 0.01 | 0.56 | |
| 4:107.6-107.8 | rs1364808 | ESN (AFR) | $3.1 \times 10^{-8}$ | **DKK2** | | | | 0.86 | 0.87 | 0.95 | 1 | [17]; |
| | rs817146 | MSL (AFR) | $9.7 \times 10^{-9}$ | **DKK2** | | | | 0.93 | 0.87 | 0.95 | 1 | Fu & Li's D, |
| | rs704049 | LWK (AFR) | $5 \times 10^{-10}$ | **DKK2** | | | | 0.92 | 0.87 | 0.95 | 1 | Fu & Li's F |
| | rs3111706 | GWD (AFR) | $1 \times 10^{-8}$ | **DKK2** | | | | 0.93 | 0.87 | 0.96 | 1 | |

| CHR:REGION | ID/BP | population | logP | gene by r2 | eQTL | GWAS | NSM | AFR | EUR | SAS | EAS | notes |
|---|---|---|---|---|---|---|---|---|---|---|---|---|
| 4:107.8-108 | rs6829139 | YRI (AFR) | $7.8 \times 10^{-10}$ | **DKK2** | | | | 0.9 | 0.97 | 0.99 | 1 | in close physical |
| | rs17037205 | LWK (AFR) | $1.3 \times 10^{-8}$ | **DKK2** | | | | 0.9 | 0.97 | 0.99 | 1 | proximity with |
| | | MSL (AFR) | $8.7 \times 10^{-9}$ | | | | | | | | | previous region |
| 5:65.2-65.3 | rs59755544 | GIH (SAS) | $1.7 \times 10^{-8}$ | **ERBB2IP** | | | | 0.44 | 0.85 | 0.86 | 0.77 | – |
| | | PJL (SAS) | $9.3 \times 10^{-10}$ | | | | | | | | | |
| | | STU (SAS) | $3.5 \times 10^{-8}$ | | | | | | | | | |
| 5:178.2-178.3 | BP178258803 | TSI (EUR) | $2.5 \times 10^{-9}$ | RP11-281O15.3 | **AACSP1** | | | 0.13 | 0.28 | 0.32 | 0.25 | Fu & Li's D |
| | | BEB (SAS) | $1.3 \times 10^{-9}$ | | | | | | | | | |
| | | ITU (SAS) | $1.7 \times 10^{-9}$ | | | | | | | | | |
| 7:19.5-19.6 | rs71530658 | CEU (EUR) | $1.4 \times 10^{-8}$ | **AC007091.1** | | (4) | | 0.03 | 0.42 | 0.15 | 0.01 | $F_{\mathrm{ST}}$ |
| | | GBR (EUR) | $1.3 \times 10^{-10}$ | | | | | | | | | |
| | | IBS (EUR) | $2.8 \times 10^{-8}$ | | | | | | | | | |
| | | TSI (EUR) | $2.8 \times 10^{-9}$ | | | | | | | | | |
| 7:98.9-99.1 | BP98971118 | GWD (AFR) | $1.2 \times 10^{-12}$ | ARPC1A | ARPC1B | (5) | | 0.5 | 0.07 | 0.06 | 0 | – |
| | rs10229886 | ESN (AFR) | $9.2 \times 10^{-9}$ | **ARPC1A** | (ARPC1B) | (5) | | 0.37 | 0.03 | 0.02 | 0 | |
| | BP98971173 | YRI (AFR) | $1.1 \times 10^{-10}$ | ARPC1A | (ARPC1B) | (5) | | 0.37 | 0.03 | 0.02 | 0 | |
| | BP98971260 | LWK (AFR) | $2.1 \times 10^{-13}$ | (ARPC1A) | (GS1-259H13.2) | (5) | | 0.37 | 0.03 | 0.04 | 0 | |
| 8:38-38.3 | rs59911155 | ESN (AFR) | $4.3 \times 10^{-8}$ | **LSM1** | **DDHD2** | **6**,(c) | (DDHD2) | 0.75 | 0.74 | 0.91 | 0.68 | – |
| | rs3739252 | GWD (AFR) | $3.4 \times 10^{-8}$ | **DDHD2** | **DDHD2** | **7**,d | (DDHD2) | 0.78 | 0.74 | 0.91 | 0.68 | |
| | | LWK (AFR) | $4.2 \times 10^{-9}$ | | | | | | | | | |
| 9:99.4-99.5 | BP99411763 | YRI (AFR) | $4.2 \times 10^{-8}$ | | | | | 0.29 | 0 | 0 | 0 | – |
| | BP99411801 | GWD (AFR) | $5.4 \times 10^{-9}$ | | | | | 0.33 | 0 | 0 | 0 | |
| | | MSL (AFR) | $1.2 \times 10^{-9}$ | | | | | | | | | |
| 9:107.3-107.4 | rs112315552 | ESN (AFR) | $3.8 \times 10^{-8}$ | **OR13C5** | NIPSNAP3A | | OR13C2 | 0.6 | 0.16 | 0.38 | 0.55 | – |
| | | GWD (AFR) | $2.4 \times 10^{-10}$ | | | | | | | | | |
| | | MSL (AFR) | $2.1 \times 10^{-9}$ | | | | | | | | | |
| | | YRI (AFR) | $2.3 \times 10^{-9}$ | | | | | | | | | |

| CHR:REGION | ID/BP | population | logP | gene by r2 | eQTL | GWAS | NSM | AFR | EUR | SAS | EAS | notes |
|---|---|---|---|---|---|---|---|---|---|---|---|---|
| 10:0.4-0.6 | rs77347335 | ESN (AFR) | $2.4 \times 10^{-10}$ | **DIP2C** | | | | 0.43 | 0.01 | 0.03 | 0.02 | – |
| | | GWD (AFR) | $2.5 \times 10^{-8}$ | | | | | | | | | |
| | | LWK (AFR) | $1.1 \times 10^{-11}$ | | | | | | | | | |
| | | MSL (AFR) | $4 \times 10^{-8}$ | | | | | | | | | |
| 10:104.6-105 | rs11191469 | LWK (AFR) | $8.4 \times 10^{-9}$ | **CNNM2** | C10orf32 | **8** | | 0.66 | 0.38 | 0.25 | 0.27 | [31] |
| | | MSL (AFR) | $6.8 \times 10^{-10}$ | | | | | | | | | |
| | | YRI (AFR) | $5.6 \times 10^{-9}$ | | | | | | | | | |
| 10:117.2-117.5 | rs150028049 | CHB (EAS) | $4.3 \times 10^{-8}$ | **ATRNL1** | | | | 0.1 | 0.23 | 0.34 | 0.62 | $F_{\mathrm{ST}}$ |
| | | CHS (EAS) | $1.9 \times 10^{-12}$ | | | | | | | | | |
| | | KHV (EAS) | $1.2 \times 10^{-8}$ | | | | | | | | | |
| 10:122.8-123 | rs2246730 | IBS (EUR) | $3.8 \times 10^{-8}$ | **RP11-159H3.2** | | | | 0.39 | 0.97 | 0.96 | 0.65 | Fay & Wu's H |
| | rs1873446 | ITU (SAS) | $1.7 \times 10^{-8}$ | **RP11-159H3.2** | | | | 0.37 | 0.97 | 0.95 | 0.64 | |
| | rs10886862 | STU (SAS) | $4.8 \times 10^{-8}$ | **RP11-159H3.2** | | | | 0.42 | 0.97 | 0.95 | 0.61 | |
| 11:65.1-65.2 | rs188162087 | BEB (SAS) | $1.1 \times 10^{-8}$ | **DPF2** | (AP003068.18) | | | 0 | 0.29 | 0.24 | 0.12 | – |
| | | GIH (SAS) | $1.7 \times 10^{-11}$ | | | | | | | | | |
| | | PJL (SAS) | $1.1 \times 10^{-8}$ | | | | | | | | | |
| 11:91.8-92 | rs11019805 | CHS (EAS) | $3.7 \times 10^{-8}$ | **FAT3** | (FAT3) | | | 0.08 | 0.31 | 0.54 | 0.59 | [30] for GBR |
| | | BEB (SAS) | $6.5 \times 10^{-9}$ | | | | | | | | | |
| | | ITU (SAS) | $4.1 \times 10^{-8}$ | | | | | | | | | |
| 12:79.7-80.2 | rs10778678 | LWK (AFR) | $3.6 \times 10^{-9}$ | **RP11-359M6.1** | **RP11-530C5.2** | | | 0.72 | 0.01 | 0.08 | 0.21 | [28, 17]; iHS |
| | rs12316084 | YRI (AFR) | $7.3 \times 10^{-9}$ | **PAWR** | RP11-530C5.2 | | | 0.54 | 0 | 0 | 0.01 | |
| | rs7306681 | MSL (AFR) | $2.5 \times 10^{-8}$ | **PAWR** | **RP11-530C5.2** | | | 0.59 | 0 | 0 | 0.01 | |
| 12:83-83.1 | rs11115333 | ESN (AFR) | $3.9 \times 10^{-10}$ | **TMTC2** | | | | 0.8 | 0.77 | 0.81 | 0.71 | – |
| | | GWD (AFR) | $9.2 \times 10^{-10}$ | | | | | | | | | |
| | | LWK (AFR) | $6.9 \times 10^{-9}$ | | | | | | | | | |
| 12:87.3-87.4 | rs11104181 | GWD (AFR) | $4 \times 10^{-8}$ | **RP11-202H2.1** | | | | 0.87 | 0.43 | 0.48 | 0.77 | [31] (KITLG for CEU) |
| | rs11503304 | ESN (AFR) | $4.8 \times 10^{-9}$ | **RP11-202H2.1** | | (9) | | 0.82 | 0.66 | 0.54 | 0.78 | |
| | rs2406741 | LWK (AFR) | $1.8 \times 10^{-10}$ | **RP11-202H2.1** | | (9) | | 0.85 | 0.66 | 0.54 | 0.78 | |
| | rs7309012 | MSL (AFR) | $7.4 \times 10^{-9}$ | **RP11-202H2.1** | | **10** | | 0.84 | 0.67 | 0.52 | 0.61 | |

| CHR:REGION | ID/BP | population | logP | gene by r2 | eQTL | GWAS | NSM | AFR | EUR | SAS | EAS | notes |
|---|---|---|---|---|---|---|---|---|---|---|---|---|
| 12:111.7-113 | rs7137828 | GBR (EUR) | $3.3 \times 10^{-11}$ | **ATXN2** | ALDH2 | **11**,e | SH2B3 | 0 | 0.46 | 0.07 | 0 | [29]; iHS |
| | | IBS (EUR) | $1.2 \times 10^{-9}$ | | | | | | | | | |
| | | TSI (EUR) | $9.9 \times 10^{-11}$ | | | | | | | | | |
| 13:28.6-28.7 | rs9554250 | CDX (EAS) | $7.1 \times 10^{-10}$ | **FLT3** | **FLT3** | **12** | | 0.28 | 0.55 | 0.45 | 0.7 | – |
| | | CHB (EAS) | $1.9 \times 10^{-10}$ | | | | | | | | | |
| | | CHS (EAS) | $9.2 \times 10^{-11}$ | | | | | | | | | |
| | | KHV (EAS) | $4.1 \times 10^{-10}$ | | | | | | | | | |
| 14:32.9-33 | rs7153204 | MSL (AFR) | $5.6 \times 10^{-9}$ | **AKAP6** | (AKAP6) | | | 0.88 | 0.08 | 0.12 | 0.29 | – |
| | rs10138310 | LWK (AFR) | $1.5 \times 10^{-8}$ | **AKAP6** | (AKAP6) | | | 0.86 | 0.1 | 0.21 | 0.3 | |
| | rs11628486 | GWD (AFR) | $1.9 \times 10^{-8}$ | **AKAP6** | (AKAP6) | | | 0.86 | 0.09 | 0.2 | 0.26 | |
| | | YRI (AFR) | $1.4 \times 10^{-8}$ | | | | | | | | | |
| 16:22.9-23.1 | rs16974808 | YRI (AFR) | $4.8 \times 10^{-9}$ | **HS3ST2** | | | | 0.79 | 0.06 | 0.11 | 0.01 | [17] |
| | rs1604799 | ESN (AFR) | $3.1 \times 10^{-8}$ | **HS3ST2** | | | | 0.79 | 0.06 | 0.11 | 0.01 | |
| | rs8063811 | MSL (AFR) | $9.5 \times 10^{-9}$ | **RP11-20G6.2** | | | | 0.79 | 0.06 | 0.11 | 0.01 | |
| 16:59.5-59.7 | rs9929021 | ESN (AFR) | $3.1 \times 10^{-8}$ | **U4** | | | | 0.76 | 0.32 | 0.46 | 0.29 | – |
| | rs9937266 | GWD (AFR) | $2.1 \times 10^{-8}$ | **U4** | | | | 0.76 | 0.35 | 0.45 | 0.32 | |
| | | YRI (AFR) | $4.2 \times 10^{-8}$ | | | | | | | | | |
| 16:81.4-81.5 | rs310010 | ESN (AFR) | $2.3 \times 10^{-8}$ | **CMIP** | | | | 0.5 | 0.06 | 0.13 | 0.1 | – |
| | | GWD (AFR) | $3.3 \times 10^{-8}$ | | | | | | | | | |
| | | MSL (AFR) | $5.1 \times 10^{-10}$ | | | | | | | | | |
| | | YRI (AFR) | $6.3 \times 10^{-10}$ | | | | | | | | | |
| 17:44-44.9 | BP44262496 | STU (SAS) | $2.6 \times 10^{-8}$ | KANSL1 | **RP11-798G7.5** | **13** | (KANSL1) | 0.03 | 0.31 | 0.6 | 0.02 | [30] (MYL4 in GBR); |
| | BP44363740 | ITU (SAS) | $1.9 \times 10^{-9}$ | (KANSL1) | **RP11-798G7.5** | **14** | | 0.03 | 0.31 | 0.63 | 0.04 | Fay & Wu's H |
| | | PJL (SAS) | $2.4 \times 10^{-10}$ | | | | | | | | | |
| 18:37.7-37.8 | rs7236000 | GWD (AFR) | $1.1 \times 10^{-9}$ | **RP11-653G8.2** | | | | 0.62 | 0.15 | 0.26 | 0.62 | – |
| | | MSL (AFR) | $5.7 \times 10^{-10}$ | | | | | | | | | |
| | rs1943603 | ESN (AFR) | $4.5 \times 10^{-9}$ | **RP11-653G8.2** | | | | 0.63 | 0.15 | 0.27 | 0.59 | |

| CHR:REGION | ID/BP | population | logP | gene by r2 | eQTL | GWAS | NSM | AFR | EUR | SAS | EAS | notes |
|---|---|---|---|---|---|---|---|---|---|---|---|---|
| 19:31.7-31.8 | BP31733665 | LWK (AFR) | $3.3 \times 10^{-11}$ | TSHZ3 | | | | 0.44 | 0.13 | 0.2 | 0.18 | – |
| | rs62101246 | ESN (AFR) | $7.4 \times 10^{-10}$ | **TSHZ3** | | | | 0.47 | 0.14 | 0.24 | 0.19 | |
| | | MSL (AFR) | $3.2 \times 10^{-9}$ | | | | | | | | | |
| 19:45.8-45.9 | rs12609631 | MSL (AFR) | $2.9 \times 10^{-8}$ | **KLC3** | | **15** | | 0.71 | 0.28 | 0.21 | 0.35 | [28] |
| | rs10853773 | ESN (AFR) | $3.5 \times 10^{-8}$ | **KLC3** | | **15** | | 0.72 | 0.27 | 0.21 | 0.34 | |
| | | YRI (AFR) | $3.5 \times 10^{-8}$ | | | | | | | | | |
| 21:17.5-17.7 | rs2823681 | GWD (AFR) | $7 \times 10^{-9}$ | **LINC00478** | | | | 0.47 | 0 | 0 | 0 | – |
| | | LWK (AFR) | $3.4 \times 10^{-10}$ | | | | | | | | | |
| | | MSL (AFR) | $2.8 \times 10^{-8}$ | | | | | | | | | |
| | | YRI (AFR) | $4 \times 10^{-8}$ | | | | | | | | | |
| 22:23-23.2 | rs2003444 | CEU (EUR) | $6.2 \times 10^{-9}$ | **IGLV3-16** | **IGLV3-12** | | | 0.49 | 0.62 | 0.65 | 0.64 | Fu & Li's D |
| | | IBS (EUR) | $1.8 \times 10^{-8}$ | | | | | | | | | |
| | | TSI (EUR) | $2 \times 10^{-9}$ | | | | | | | | | |
| | | BEB (SAS) | $8.3 \times 10^{-10}$ | | | | | | | | | |
| | | GIH (SAS) | $3.8 \times 10^{-10}$ | | | | | | | | | |
| | | ITU (SAS) | $6.6 \times 10^{-11}$ | | | | | | | | | |
| | | PJL (SAS) | $9.4 \times 10^{-9}$ | | | | | | | | | |
| | | STU (SAS) | $1.4 \times 10^{-9}$ | | | | | | | | | |
